# Single-Cell Morphological Profiling Reveals Insights into Programmed Cell Death

**DOI:** 10.1101/2025.01.15.633042

**Authors:** Benjamin Frey, David Holmberg, Petter Byström, Ebba Bergman, Polina Georgiev, Martin M Johansson, Patrick Hennig, Jonne Rietdijk, Dan Rosén, Jordi Carreras-Puigvert, Ola Spjuth

## Abstract

Analysis at the single-cell level is a powerful approach to study biological processes and responses to perturbations. However, its application in morphological profiling with phenomics remains underexplored. Here, we use the Cell Painting assay to investigate morphological effects of 53 small molecule compounds, associated with six distinct programmed cell death mechanisms, across six concentrations in MCF7 cells. To compare single-cell and aggregated analysis strategies, we conduct both supervised and unsupervised evaluations aimed at identifying features linked to programmed cell death. We apply an energy distance as a metric to quantify morphological perturbation strength, enabling efficient filtering. Among three tested feature extraction methods, self-supervised DINO embeddings applied to single-cell data captured high-resolution morphological patterns. Focused analyses of apoptosis-inducing compounds revealed biological heterogeneity attributable to specific molecular targets and concentration-dependent effects, which were not apparent in aggregated profiles. In contrast, multi-class classification models for the six programmed cell death mechanisms trained on single-cell features achieved F1 scores of 79.86%, while models trained on aggregated features reached F1 scores of up to 89.97%. Our results highlight the advantages of single-cell data for unsupervised exploration and show that aggregated representations yield more robust and accurate performance in supervised models.

## 1 Introduction

Single-cell data has become a cornerstone of modern biological research, providing unprecedented insights into cellular heterogeneity, immune responses, and disease mechanisms. Specifically in the field of transcriptomics, single-cell analysis has enabled the identification of subpopulations within heterogeneous samples, aided in uncovering the effects of perturbations, and advancing our understanding of tumor biology (Bunne et al., 2023; Lotfollahi et al., 2023; Peidli et al., 2024; Y. Wu et al., 2024; Yao et al., 2022). Single-cell sequencing technologies have proven invaluable for exploring immune responses and elucidating disease mechanisms, particularly in the context of cancer biology (Olofsson et al., 2021). Despite the growing impact of single-cell sequencing, the application of high-throughput single-cell data for characterizing specific programmed cell death (PCD) subtypes remains an open challenge.

The analysis pipeline for Cell Painting data begins with a feature extraction step that leverages tools like CellProfiler (Carpenter et al., 2006), a dedicated software that extracts pre-defined features from cellular images. CellProfiler has been instrumental in advancing morphological profiling by enabling researchers to extract detailed cellular characteristics (Harrison et al., 2023; Haslum et al., 2023; Rietdijk et al., 2021). However, the reliance on hand-crafted features has limitations, particularly as data scales and biological complexity increases. Recent advances in computer vision have introduced alternative approaches that leverage deep neural networks and optimized feature sets for automated feature extraction (V. Kim et al., 2024; Kraus et al., 2024; Stossi et al., 2024). These approaches include convolutional neural networks (CNN) (Tian et al., 2023), weakly-supervised (Moshkov et al., 2024), and self-supervised models (Caron et al., 2021;K.He et al., 2021; Kraus et al., 2024).

Self-supervised vision models, in particular, have shown great promise in morphological profiling as they do not require extensive data labeling. One example, DINO, employs two parallel networks to learn representations from different crops of the same image, enabling robust feature extraction. DINO has been successfully applied to Cell Painting data, outperforming existing methods in MOA prediction and toxicity estimation tasks (Doron et al., 2023; Gupta et al., 2024; V. Kim et al., 2024; Moshkov et al., 2024; Pfaendler et al., 2023). Similarly, Masked Autoencoders, which use pixel masking and encoder-decoder architectures, have demonstrated state-of-the-art performance on Cell Painting datasets. Notably, Recursion Pharmaceuticals recently released an open version of a Masked Autoencoder model trained on millions of Cell Painting images, a first foundation model for Cell Painting data (Kraus et al., 2024).

Despite these advancements, the majority of morphological profiling workflows where cells are perturbed using chemical or genetic perturbations aggregate features extracted from single-cell imaging into average profiles. By combining profiles from the same treatments and using mean/median cells as the main objects, outliers and data complexity can be reduced (Caicedo et al., 2017; Moshkov et al., 2024). While this approach improves data separation at the treatment level, it effectively sacrifices information about cellular heterogeneity and subpopulations at the single-cell level. This trade-off limits the deeper biological characterization of individual cells, where heterogeneity can reveal essential insights into responses to perturbations.

Applications where Cell Painting experiments have utilized single-cell data in the analysis remain relatively limited. Early efforts focused on tasks like protein co-localization analysis, classification, and the characterization of blood and immune cell types (Doron et al., 2023; Gupta et al., 2024; Moshkov et al., 2024; Pahl et al., 2023; Pfaendler et al., 2023). One specific application where single-cell analysis from Cell Painting has not been explored extensively is PCD. Previous works from Lippincott et. al and Chen et. al have applied Cell Painting and high-content imaging on single-cells to study pyroptosis and the related inflammasome respectively (D. Chen et al., 2024; Lippincott et al., 2025). In another work, Schorpp et al. classified different cell death subtypes using a CNN and larger image crops from four-channel Cell Painting experiments (Schorpp et al., 2023), covering two cell death subtypes (apoptosis, Ferroptosis, as well as non-treated cells) at seven inducing compounds per subtypes in five concentrations per compound. For their classification, they compared an end-to-end CNN with 512 × 512 crops as inputs with traditional machine learning models trained on 245-dimensional aggregated feature vectors.

In this manuscript we present a comprehensive exploration of six programmed cell death subtypes induced by 53 compounds in MCF-7 breast cancer cells, each tested at six different concentrations using five-channel Cell Painting. In order to assess the relevance of single-cell morphological profiles for studying PCD subtypes, we apply unsupervised and supervised methods and compare the performance of three feature extraction approaches: CellProfiler (Stirling et al., 2021), DeepProfiler (Moshkov et al., 2024), and the self-supervised transformer-based DINO (Figure1 A)

**Figure 1.**
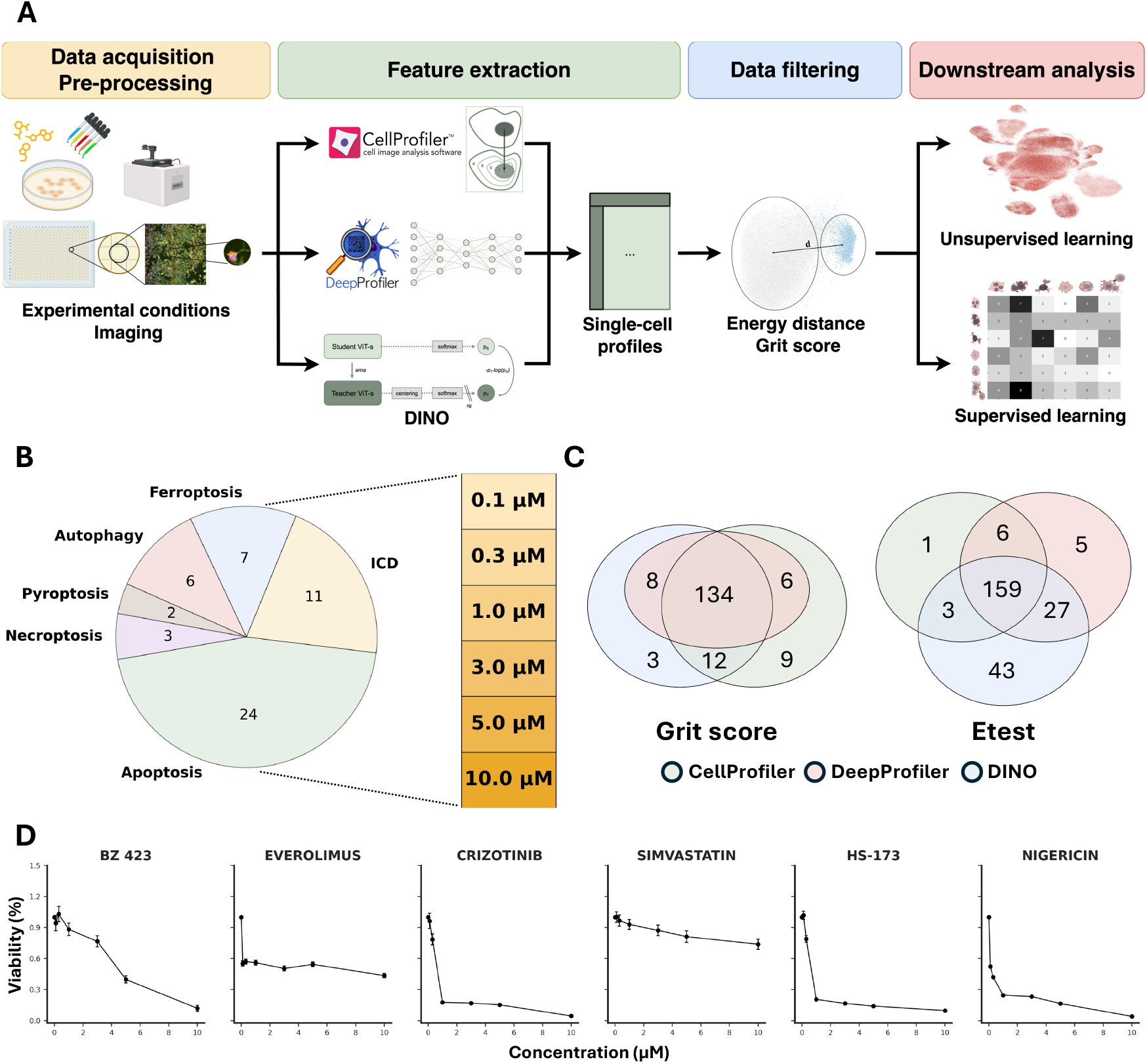
Overview of experimental and analysis pipeline and filtering outcomes. (A) Schematic of the workflow used in this work. Experimental conditions are established, followed by imaging and feature extraction using CellProfiler, DeepProfiler, or DINO. Each extracted feature set undergoes separate data filtering and downstream analysis, including metric calculations, unsupervised and supervised learning. (B) Distribution of compound counts across different of cell death subtypes. Bar chart indicates concentrations used for the different compounds. The same concentrations were used in each compound. (C) Venn diagram depicting number of significant compound concentration sets between CellProfiler, DeepProfiler and DINO. Left diagram shows filtered grit sets, right diagram shows significant etest sets. (D) Dose-dependent viability per PCD subtype (1 randomly selected compound per subtype): Apoptosis, Autophagy, ICD, Ferroptosis, Necroptosis, Pyroptosis (from left to right). Mean ± standard error (n = 4 technical replicates).

## 2 Methods and Materials

### 2.1 Compounds and programmed cell death subtypes

53 compounds were selected where annotations spanned six distinct programmed cell death subtypes: apoptosis, Necroptosis, Ferroptosis, Pyroptosis, as well as the non-regulated Autophagy and Immunogenic Cell Death (ICD) (D. Chen et al., 2024). The compounds and PCD subtype annotations are shown in Figure 1B; and with their associated targets in Supplemental Table S1.

Apoptosis, as the most extensively studied form of cell death, is characterized via cell shrinkage followed by fragmentation and development of apoptotic bodies (Kerr et al., 1972). Necroptosis describes cell death defined by membrane rupture and subsequent release of cellular contents. Necroptotic processes are induced by tumour necrosis factor (TNF) leading to a RIPK and caspase 8 activation (Conrad et al., 2016). Ferroptosis is driven by metabolic processes, specifically involving fatty acyl moieties dependent on iron (Stockwell, 2022). It has been shown to be connected to different types of degenerative diseases involving the brain, heart and kidney. Pyroptosis presents a natural cellular defense mechanism induced by pathogen-associated molecular patterns (PAMPs) and damage-associated molecular patterns (DAMP) which trigger cell swelling, lysis and the release of pro-inflammatory cytokines, and shown to have cancer-related indications (Yu et al., 2021a). Autophagy is a process driven by the lysosomal machinery, resulting in the self-degradation of cellular components (Shen et al., 2023). Immunogenic cell death, on the other hand, is a form of regulated cell death that elicits an adaptive immune response against antigens derived from dying neighbouring cells, induced for instance by DNA damaging agents (A. Tesniere et al., 2010). Immunogenic cell death (ICD), on the other hand, is a form of regulated cell death that elicits an adaptive immune response against antigens derived from dying neighboring cells, induced for instance by DNA damaging agents (Tesniere et al., 2010). ICD is characterized by the release and surface exposure of specific DAMPs, acting as endogenous adjuvants to recruit highly specialized immune cells, such as CD8+ T cells, leading to the targeted destruction of the affected cell (Kroemer et al., 2013)

### 2.2 Cell culture

MCF7 human breast cancer cells (Sigma-Aldrich #86012803) were cultured to passage +24 and confirmed to be Mycoplasma-free using the MycoAlert kit (Lonza #LT07-218). Cells were maintained at 37^°^C, 5% CO_2_, and 95% humidity in Dulbecco’s Modified Eagle Medium (DMEM; Gibco #31885-023) supplemented with 10% (v/v) heat-inactivated fetal bovine serum (FBS; Gibco #10500064) and 1% (v/v) Penicillin-Streptomycin (Gibco #15140-122). At 80-90% confluency, cells were washed with Dulbecco’s Phosphate-Buffered Saline (DPBS; Gibco #14190250), dissociated using TrypLE Select (Gibco #A1217701), and reseeded in fresh complete growth medium in T75 flasks (Thermo Scientific #156499).

### 2.3 Cell seeding and compound treatment

All test compounds were provided as DMSO solutions and diluted to stock concentrations using a 10-fold serial dilution. Phenotypic reference compounds that are known to induce a distinctive phenotype in a variety of cells (Willis et al., 2020), including Etoposide (Sigma-Aldrich #E1383), Fenbendazole (Sigma-Aldrich #F5396), and Staurosporine (Sigma-Aldrich #19-123-M), were purchased from Sigma-Aldrich and prepared as 10 mM stocks in DMSO (Sigma-Aldrich #D8418). Assay-ready plates containing test compounds, phenotypic reference compounds, and DMSO (negative controls) were prepared using an I-DOT (immediate drop-on-demand technology) liquid handler. Compounds were dispensed in precise volumes to achieve final treatment concentrations of 0.1, 0.3, 1, 3, 5, and 10 *µ*M for test compounds; 3 *µ*M for the positive controls Etoposide and Fenbendazole; 0.3 *µ*M for the Staurosporine control; and 0.1% for DMSO.

Each of the 8 compound plates included 20 or 21 test compounds at 6 concentrations and 2 technical replicates, 3 phenotypic reference compounds at 1 concentration and 2 technical replicates, and 14 wells of DMSO. The distribution of treatments and controls on the compound plates was optimized using PLAID (Plate Layouts using Artificial Intelligence Design) (Francisco Rodríguez et al., 2023), a flexible constraint-programming model that ensures a robust microplate layout.

Prior to cell seeding on the assay ready plates, the compounds pre-spotted onto the assay plates were dissolved by adding 20 *µ*L of complete growth medium per well using a Biotek Multiflo FX dispenser. The plates were then placed on a shaker at 150 rpm at room temperature for one hour to ensure complete dissolution of the compounds. Cells from passage 24 were counted and seeded on top of the plates with dissolved compounds at a density of 1100 cells per well by dispensing 20 *µ*L/well of cell suspension with Biotek Multiflo FX dispenser, resulting in a total final volume of 40 *µ*l/well. The seeded plates were left to stand at room temperature for 20 minutes before being incubated at 37^°^C in a 5% CO_2_ atmosphere for 48 hours.

### 2.4 Cell Painting assay

Cell painting experiments were conducted according to the protocol described Bray et al., 2016 with minor modifications. The fluorescent labeling process was performed in an automated laboratory setup, which included a UR10 robotic arm for handling plates, a Biotek MultiFlo FX for dispensing solutions, and a BlueWasher (Blue Cat Bio) for washing steps. In brief, MCF7 cells were cultured in 384-well plates and treated with compounds at various doses for 48 hours. Following treatment, cells were live-cell-labeled with MitoTracker Deep Red (Invitrogen) and subsequently fixed with 4% paraformaldehyde (Histolab) for 20 minutes. After fixation, cells were permeabilized with 0.1% Triton X-100 (Cytiva) for 20 minutes and stained with a solution containing Hoechst 33342, Wheat Germ Agglutinin, Phalloidin, SYTO 14, and Concanavalin A (all from Thermo Fisher Scientific) for 20 minutes. Finally, plates were washed, sealed, and stored at 4^°^C, protected from light until image acquisition.

### 2.5 Imaging and image pre-processing

Fluorescence microscopy was performed using a high-throughput ImageXpress Micro XLS (Molecular Devices), capturing nine sites per well across five fluorescence channels. Excitation and emission wavelengths were optimized for each dye. Image processing and analysis were conducted with CellProfiler and CellPose 2.0 (Stringer et al., 2021), including quality control, illumination correction, segmentation, and feature extraction. Quality control was performed via filtering out images that surpassed intensity limits in selected features measuring key intensity levels of segmented objects. Illumination correction for CellProfiler and DINO images was calculated and applied via the CellProfiler *CorrectIlluminationApply* module using the *FitPolynomial* function. For DeepProfiler, the illumination correction function inside the DeepProfiler package was used according to (Singh et al., 2014). A total of 107,405 images were processed. This number comprises of 8 384-well plates that were imaged at 9 field of views (“sites”) per well in five-channels. Outer wells were left out and QC-flagged images removed, which results in the above number. Segmentation results were saved as binary masks and later used as input for feature extraction (CellProfiler), image cropping (DeepProfiler) and training (DINO).

### 2.6 CellProfiler

A feature extraction pipeline using Cell Profiler version 4.3.4 (available at https://doi.org/10.17044/scilifelab.21378906.v2 and described in (Harrison et al., 2023)) was run and resulted in a 2,160-dimensional feature vector for each cell. These features capture fluorescence intensity, location, granularity, and other measurements of the imaged cells, grouped into cell, nucleus, and cytoplasm features.

### 2.7 DeepProfiler

DeepProfiler is a convolutional neural network trained in a weakly supervised setting, as described by (Moshkov et al., 2024). In this study, we utilized the pre-trained model CellPainting_CNN with weights from (Moshkov et al., 2024) for inference without additional training. Features were extracted as 672-dimensional vectors from the block_6a_-activation layer of the EfficientNetB0 architecture, generating one feature vector per single-cell crop. To ensure accurate representation, we used nucleus-centered image crops with a resolution of 150 × 150 pixels. The cropping size was selected to fully encapsulate 95% of all cells. Illumination correction was applied during the DeepProfiler feature extraction process, using a down_scale_factor of 8 and a median_filter_size of 48. Importantly, segmentation masks were omitted during inference to retain contextual neighborhood information surrounding the cells.

### 2.8 DINO

Self-distillation with no labels (DINO) falls under the self-supervised learning paradigm to learn representations from images without the need for explicit label annotation (Caron et al., 2021). The approach combines identical student and teacher networks to learn image features through self-distillation. We trained DINO with a ViT-small network, a patch size of 16 pixels, and five input channels. We adapted the architecture by (Doron et al., 2023), to allow for multi-channel inputs. The network was trained from scratch with a stratified and sub-sampled subset of our Cell Death dataset consisting of >185,000 single-cell images, comprising all compounds at all concentrations (including DMSO). As network inputs, we used the segmented five-channel site-level images, center-cropped around the nuclei, and stacked the images into a (150 × 150 × 5) image for each cell. We excluded cells at the borders (with a 150 pixel radius) to avoid cells being cut off. For training purposes, pixel intensities were normalized to [0, 1] and images resized to 224 × 224 pixels. Using the same set of augmentations as (Doron et al., 2023) (random channel dropping, random brightness and contrast, artificial warping), we trained the DINO network for 200 epochs with a learning rate of 0.0005. Training was performed on a single NVIDIA GTX 3090 with a batch size of 20 for 3 days. Inference was performed on the teacher network, resulting in 384-dimensional single-cell feature vectors. To evaluate the model, we inferred 625,000 stratified, randomly sampled cells from our cell death dataset that were held out during training. A full set of hyperparameters can be found in Table S3. For a full description of the augmentations, we refer to the *config_finetune_unmasked*.*yaml* on our FigShare.

### 2.9 Data post-processing

To ensure comparability, we applied the same post-processing and analysis pipeline to all three feature sets. In the following, we will also refer to the extracted feature vectors as “profiles”. Feature vectors were first standardized to the negative control (DMSO) by subtracting the mean and dividing by the standard deviation of the DMSO cells per plate. For our aggregated analysis, we calculated the median profile per field of view (“site”). Feature selection was performed by removing missing value features, highly correlated (>99%), low variance (<0.01%) and blacklisted features as defined in the pycytominer package (Way et al., 2022). Additionally, features with >25% constant values across cells were removed. This resulted in just over 1,400 features for CellProfiler, 384 features for DINO, and 672 for DeepProfiler, respectively. Additionally, we excluded cells bordering the images within a 150 pixel radius to avoid cells being cut off. (this was done post feature extraction to ensure correct performance of the neural network-based feature extractors). In short, we applied this filter: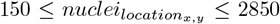. A summary of cell counts per compound concentration prior to filtering and sampling can be found in Supplemental Figure S1 as well as viability plots in Supplemental Figures S2-S4.

### 2.10 Sampling

In order to obtain cell sets more representative of the true population distribution and taking into account the over-abundance of DMSO wells in our plate layouts, we implemented a sampling strategy:

- Perform a stratified sampling by plate, well, and site (the latter iff the resulting cell count after sampling is large enough).
- To obtain the specific sampling per well/ site follow the bellow strategy which we denote as well_ratio sampling:

Let *s* ∈ (0, 1] be a user-defined sampling rate and *n* ∈ N a predefined minimum number of cells per condition. Define *W* as the set of all wells belonging to DMSO and *V*_*U*_ as all wells belonging to Treatment *U* (*T* be the set of all treatments). Then, *D*_*w*_ := {*d*_*i*_ | *d*_*i*_ ∈ *N* ∧ *d*_*i*_ ∈ *w, w* ∈ *W*} describes the set of negative control cells of a specific well *W* and {*U*_*v*_ := *t*_*i*_ | *t*_*i*_ ∈ *C*_*U*_ ∧ *t*_*i*_ ∈ *v, v* ∈ *V*_*U*_}} the treated cells of a specific well *V* and compound *U*. Then, we we define well-ratio sampling as:

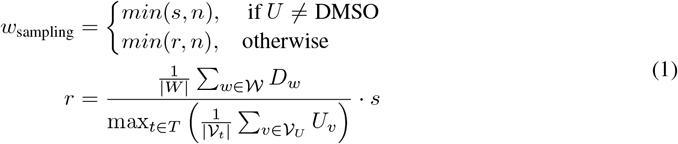

*w*_*sampling*_ can be understood as the ratio of the average DMSO cell count per well divided by the maximum average cell count per well across all treatments. In case of *r* ≥ 1, the sampling simplifies to:

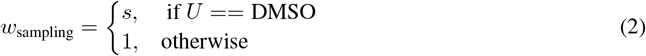

This sampling strategy was used in our unsupervised and exploratory analysis tasks as well as to define a training and inference set for our DINO training. In the unsupervised scenario, we matched our CellProfiler and DeepProfiler sets after etest filtering to match cell counts in the DINO set.

### 2.11 Energy distance

Energy distance (e-distance) (Peidli et al., 2024) was originally implemented for single-cell sequencing data as a metric for perturbation strength. We calculated e-distances per compound concentration set using single-cell profiles and the first 50 principal components.

Let *x*_1_, …*x*_*N*_ ∈ *C*_1_ and *y*_1_, …*y*_*M*_ ∈ *C*_2_ be single-cell profiles, where *N, M* represent the respective sample size and *C*_*T*_be our treatment spaces. We further define:

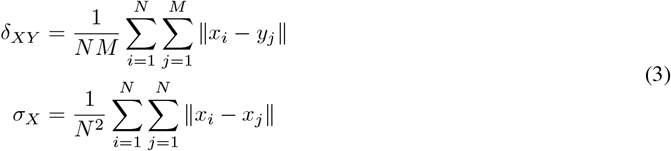

The energy distance is then defined as:

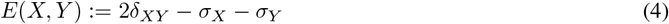

Using the e-distance as a test statistic, we performed a permutation test with 50,000 permutations to calculate a p-value for each compound concentration in relation to the negative control. We removed all compound concentrations with an adjusted p-value > 0.05. For multiple testing correction, we used the *holm-sidak* approach.

### 2.12 Grit score

Grit score measures the perturbation strength and reproducibility of a given treatment group. We calculated grit scores on a compound-concentration level using aggregated profiles. We filtered compound concentrations with a grit score below 1.5.

Let *d*_1_, …*d*_*N*_ ∈ *N* be morphological profiles of a control group and *t* ∈ *C*_*T*_ a target profile of given treatment *T* with *M* replicate profiles *r*_1_, …, *r*_*M*_. Then define *D* := {*d*_1_, …, *d*_*M*_} where *d*_*i*_ = *corr*(*t, c_i_*) and *S* := {*s*_1_, …, *s*_*M*_} where *s*_*i*_ = *corr*(*t, r_i_*). Let further *µ*_*D*_, *σ*_*D*_ be the respective statistics of *D*. Then the grit score *G* is defined as:

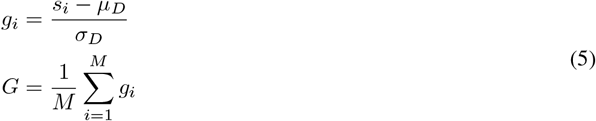

where *M* represents the number of replicates.

Both e-distance and grit score were calculated with respect to the negative control, which was then omitted for further analysis. Reference compounds in Supplemental Table S2 that were not annotated with specific cell death subtypes were omitted.

### 2.13 Mean average precision

Mean average precision (mAP) is a performance metric used in cellular profiling to assess how well a ranking method brings perturbation groups to the top of a list. It combines precision and recall into a single scalar averaging the precision values at every rank where a relevant item appears, and then averages those values across all queries (Kalinin et al., 2025). We computed mAP across perturbations by treating each unique compound concentration as a separate class, with profiles from sites treated with the same compound concentration. For each query, we calculated the cosine similarity between its profile and all others, ranked the profiles by descending similarity, recorded precision at each rank where a relevant profile occurred (the number of relevant profiles in the top k divided by k) and averaged those precision values to obtain the average precision (AP). We averaged AP over all queries to yield the mAP. For statistical evaluation, we selected a *null_size* of *n = 500,000*, referring to the Supplemental materials by Kalinin et al. (Kalinin et al., 2025). We selected a p-value cut-off of 0.05 as well as *holm-sidak* for multiple testing correction (we used these parameters to ensure comparability with our etest calculation).

### 2.14 Visualisation and clustering

We used the scanpy (Wolf et al., 2018) library to calculate PCA, neighborhood graphs, and UMAP embeddings. We trained our UMAP on single-cell profiles using PAGA for initialization and cosine similarity as the distance metric. For visualization purposes, cells extracted with CellProfiler and DeepProfiler were sampled to 10% in a stratified manner. For DINO results, we used the full data set post-filtering as it matched the sampled CellProfiler and DeepProfiler sets in size.

### 2.15 *k*^*^ Distribution Analysis

To compare our feature representations, we computed *k*^*^ distributions for each feature extractor (CellProfiler, DeepProfiler, and DINO) at both the aggregated and single-cell level. The *k*^*^ statistic captures local neighborhood consistency with respect to class membership and was proposed as a way to compare latent spaces (Kotyan et al., 2024). Given a sample *x*_*i*_ with true label *y*_*i*_, we compute pairwise cosine distances between all samples and rank them in ascending order. The *k*^*^ value is defined as the index of the first nearest neighbor whose class label differs from that of *x*_*i*_, formally expressed as

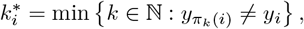

where *π*_*k*_(*i*) denotes the index of the *k*-th nearest neighbor of *x*_*i*_, and *y*_*πk*_(*i*) is its associated class label.

To account for class imbalance, 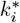 is normalized by the total number of samples *n*_*yi*_belonging to class *y*_*i*_, yielding the normalized score:

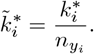

The resulting set of values 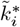 for all samples *i* ∈ *S* of a given class defines the *k*^*^ distribution for that class. We then summarize this distribution using three descriptive statistics. The mean of the distribution

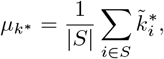

the standard deviation

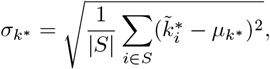

and the skewness defined as

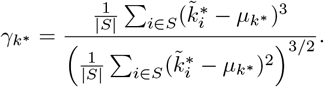

We report these statistics to characterize the local neighborhood structure in feature space. A lower *µ*_*k*_* indicates that class members tend to be surrounded by similar samples, while a higher *γ*_*k*_* reflects skewness toward tight clustering. Conversely, negative skewness suggests fragmented or overlapping class neighborhoods, where a *γ*_*k*_* <-0.5 indicates clustered distributions, a values of −0.5<*γ*_*k*_* <0.5 indicates overlapped and *γ*_*k*_* > 0.5 fractured distributions (Kotyan et al., 2024).

### 2.16 Supervised model training

The extracted profiles from CellProfiler, DeepProfiler, and DINO were used to train a classifier both on single-cell profiles and the aggregated profiles. First, profiles were split into a training and testing set (70/30), stratified over Cell Death subtypes. As our dataset contained a limited number of compounds per Cell Death subtype, we implemented a split with no overlap between seeding wells between the training and testing set. Next, a set of models was trained using the AutoGluon Tabular framework (Erickson et al., 2020). The model selection included models from the XGBoost, CatBoost, Extra Trees, Random Forest, and FastAI NN families (Breiman, 2001; T. Chen et al., 2016; Dorogush et al., 2018; Korsunsky et al., 2019). AutoGluon Tabular runs an automated stacking, combination, cross-validation and hyper-parameter tuning of models and reports a ranked list of trained models. We ran AutoGluon Tabular on the “high_quality” setting without custom models and removed the KNN model family. The training time was set to 24 hours. The two top-performing models were selected for reporting, ensuring overlap between the training scenarios. In addition to single-cell and aggregate classifications, we implemented a third strategy (‘Majority vote’) where we first predicted cell death subtypes on single-cell level followed by a majority vote over predicted labels per site. A prediction was assigned as correct if and only if the majority-voted label matched the true label. To evaluate our classification model, we reported F1, Recall, Precision as well as Accuracy. All metrics were reported on macro and micro (weighted) level. Additionally, we generated confusion matrices for all models.

### 2.17 Data and Code availability

The code to run the analysis and generate the figures can be found on Github (https://github.com/pharmbio/sc_-celldeath). The feature profiles, metrics, and splits to run the analysis and supervised training can be found on FigShare: https://doi.org/10.17044/scilifelab.28202864.

## 3 Results

### 3.1 Measuring perturbation strength using e-distance enables robust cell filtering

Before quantifying perturbation effects, we first evaluated cell viability and cell counts to determine whether dose-dependent responses were detectable and to exclude compounds or conditions that showed no activity or induced widespread cell death (Fig. 1D, Supplemental Figures S1-S4). We then calculated grit scores and e-distances for groups of single cells treated with the same compound concentration, including only those groups with at least 200 cells. Dimethyl Sulfoxide (DMSO), used as a solvent control, served as the negative control in these assessments. For all downstream analyses, DMSO-treated cells were excluded, as they did not exhibit phenotypes consistent with any of the programmed cell death subtypes under investigation.

Grit calculations resulted in 54.04% of sets kept for CellProfiler features, 53.43% for DeepProfiler features, and 51.56% for DINO features. Notably, four compound concentration sets extracted with CellProfiler showed significantly higher grit scores than in DeepProfiler. Further inspection revealed that these sets did not appear in DeepProfiler features as post-processing of the cells resulted in lower cell counts and subsequent removal (Table 1). The Venn diagram (Figure 1C) shows that 134 sets resulted in a grit > 1.5 for all feature extractors, with 20 sets only in DINO and CellProfiler features. Inspection of the distributions of grit scores further reveals that DINO features result in a significant proportion of sets with grit scores > 4. This indicates strong phenotypical changes when compared to DMSO (Supplemental Figure S5).

**Table 1.**
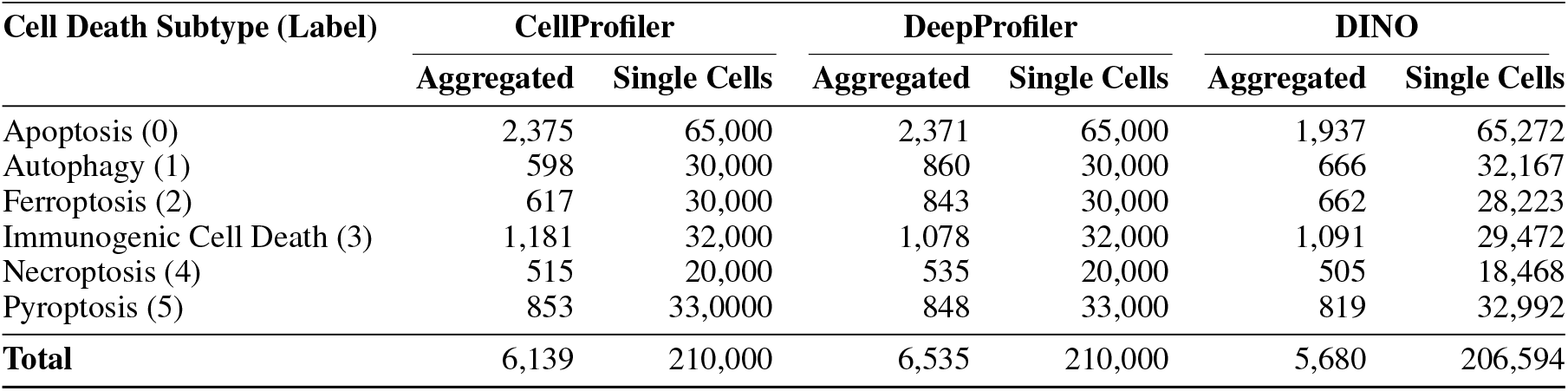
Number of samples used for aggregated and Single-Cell classification models across Feature Extractors.

Etest and e-distance results revealed similar distributions in distance and significance results between CellProfiler, DeepProfiler and DINO features (Supplemental Figure S6). For CellProfiler, higher absolute distances were observed with a maximum e-distance of 11,590 as compared to 3,090 in DeepProfiler and 420.93 for DINO (Supplemental Figure S6). Due to the non-interpretability of the distances, filtering was performed using results from the corresponding etest. Results show 74.90% of sets as significant for DeepProfiler features, 66.54% significant for CellProfiler features and 71.00% for DINO. Here, 155 common sets were observed between the three feature sets (Figure 1C). It is noteworthy that the number of sets differs between grit and e-distance calculations, as the metrics have different underlying assumptions. Therefore, the intersection of significant etest and grit-scores >1.5 was used as a cut-off for further analysis.

We further calculated mean average precisions for all compound concentrations, observing similar performance to the etest calculations with an overlap of 162 groups with etest in CellProfiler (Supplemental Figure S7A), 169 groups in DeepProfiler (Supplemental Figure S7B) and 187 groups with DINO (Supplemental Figure S7C). Additionally, we found similar levels of phenotypic activity between DeepProfiler (70% mAP, 74% etest), Supplemental Figure S7B) and DINO (79% mAP, 85% etest, Supplemental Figure S7C), with a larger difference in CellProfiler (81% mAP, 66% etest, Supplemental Figure S7A).

### 3.2 DINO features learn higher resolution morphological patterns on single-cell level

In order to characterize the selected PCD subtypes from a single-cell perspective, we used different feature extraction methods applied to single-cell data as well as aggregated (median cell) data.

We first compared the feature sets based on lower-dimensional UMAP embeddings. Our results show stronger separation of PCD subtypes at the single-cell level using DeepProfiler features compared to CellProfiler (Figure 2A). A clear separation of the Pyroptosis class is observable in both the single-cell UMAP along with the marginal kernel density estimations, as well as in the corresponding correlation matrix (Figure 2C). For CellProfiler features, we observe a positive correlation of profiles from the Pyroptosis, Ferroptosis, and Autophagy classes. For DeepProfiler features, we notice positive correlations between profiles from the Ferroptosis and Autophagy classes, but a clear separation from the Pyroptosis class. Further, a clear cluster formation of the Apoptosis class can be observed in the DeepProfiler embedding. We identified the compound FK866 to be responsible for the formation of this cluster as indicated in Supplemental Figure S8B middle. When comparing the results with the DINO embedding, more noticeable cluster formation on a single-cell level as well as in the correlation maps can be found (Figure 2A and C DINO). Clusters appear to be more dense as indicated by the marginal KDEs in the single-cell UMAP as well as by highly correlated profiles in the corresponding heatmaps (Figure 2). When taking into account the compounds, we observed a clear pattern; individual compounds corresponding to the observed clusters in morphological space (Supplemental Figure S8B). Density estimations of the PCD subtypes in our UMAP spaces revealed an overall more uniform distribution of cells in both CellProfiler and DeepProfiler features (Supplemental Figure S9A and B), while the subtypes in our DINO embedding space form multiple sub-clusters (Supplemental Figure S9C).

**Figure 2.**
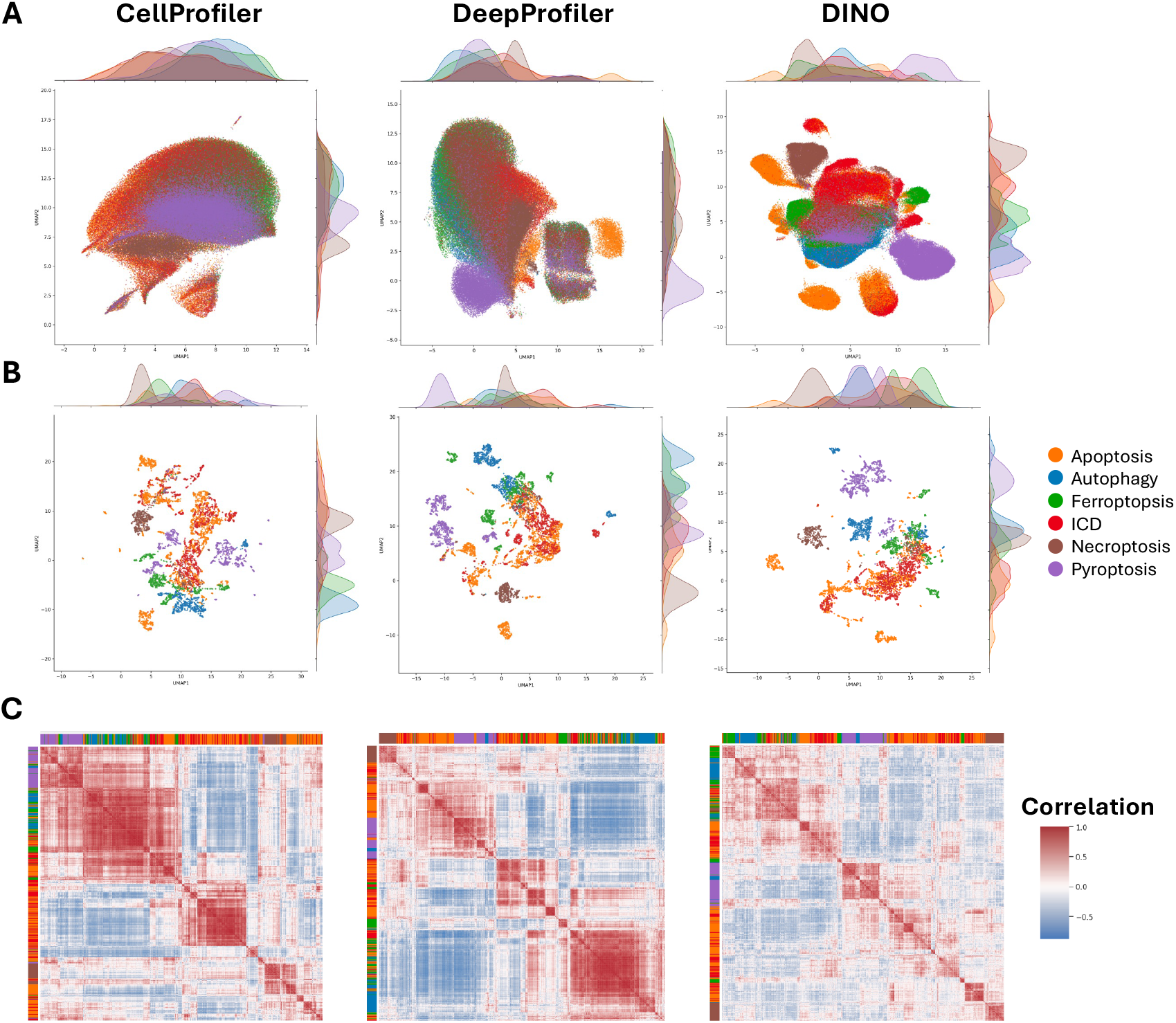
Comparison of morphological profiling across feature extraction methods. (A) UMAP embeddings of cell morphologies extracted using CellProfiler, DeepProfiler, and DINO. Each point represents a single-cell, colored by its cell death subtype. Marginal density plots along the axes display the distribution of cells for each subtype, highlighting clustering patterns unique to each method. (B) UMAP embeddings of aggregated cell morphologies extracted using CellProfiler, DeepProfiler, and DINO. Each point represents a aggregated profile, colored by its cell death subtype. Marginal density plots along the axes display the distribution of cells for each subtype, highlighting clustering patterns unique to each method. (C) Correlation heatmaps comparing pairwise feature similarities among cell treatments. Rows and columns correspond to site-level aggregated profiles, with colors representing correlation values (red for positive and blue for negative correlations). The distribution of cell death subtypes is indicated by colored bars along the heatmap axes.

Subsequent analysis of median profiles revealed a clearer separation of PCD subtypes and specific compounds in all three feature sets (Figure 1B and Supplemental Figure S8A). In all feature sets, we observed a region of strong overlap between the cell death subtypes and identified these regions as groups with lower grit and e-distance scores and less separable (Figure 2B). Overall, the differences between feature sets on an aggregated level are less apparent when compared to single-cell level.

One noticeable observation in the DeepProfiler features is the clearly separated cluster in the UMAP embedding (Figure 2A.) The cluster was identified to consist of compounds from all PCD subtypes without any distinct pattern indicative of its formation (Figure 2 DeepProfiler and Supplemental Figure S10). Analysis of the corresponding microscopy images using our *CellViewer* tool did not reveal any patterns in the images that would explain this cluster (Supplemental Figure S10C). Further analysis of the original feature intensities revealed a distinct pattern in all cells originating from that cluster (Supplemental Figure S10B), but due to the non-interpretability of the features, we were limited in our understanding of this anomaly. As a removal of the blob did not result in a change of the remaining morphological space, we kept the points in for all further analysis.

To further quantify the differences in the embeddings, we calculated the corresponding *k*^*^ distributions S11. The summary statistics reveal differences in the distributions between single-cell and aggregated profiles across all feature extractors, where overall skewness is higher in single-cell profiles (Table 2). While all single-cell distributions are positively skewed, the degree of skewness varies between cell death subtypes and feature extractors. Overall, we observe lower skewness values for DINO features (1.53-3.73) compared to CellProfiler (3.29-6.21) and DeepProfiler (2.96-6.38). Similarly, the highest degree of skewness is found in the apoptosis, ferroptosis and ICD class, while the remaining classes show overall lower skewness. For aggregated profiles, we observed a shift in distributions, specifically in the necroptosis, pyropotosis and autophagy class. For nectroptosis, we see the emergence of overlapped skewness patterns for all feature extractors, while for pyroptosis our DeepProfiler features indicate a clustered distribution as indicated by a skweness below −0.5.

**Table 2.** Summary of *k*^*^ distribution statistics for each cell death subtype, feature extractor, and level of representation. Lower *µ*_*k*_* and higher *γ*_*k*_* indicate tighter and more homogeneous class-specific neighborhoods.

| Class | Extractor | Aggregated |  |  | Single-cell |  |  |
| --- | --- | --- | --- | --- | --- | --- | --- |
| | | $\mu_{k^*}$ | $\sigma_{k^*}$ | $\gamma_{k^*}$ | $\mu_{k^*}$ | $\sigma_{k^*}$ | $\gamma_{k^*}$ |
| Apoptosis | CP | 0.017 | 0.022 | 1.47 | 0.001 | 0.001 | 4.90 |
|  | DP | 0.023 | 0.031 | 1.43 | 0.002 | 0.007 | 4.91 |
|  | DINO | 0.027 | 0.035 | 1.19 | 0.009 | 0.019 | 2.40 |
| Autophagy | CP | 0.124 | 0.104 | 0.68 | 0.001 | 0.001 | 3.29 |
|  | DP | 0.145 | 0.145 | 0.63 | 0.001 | 0.002 | 5.50 |
|  | DINO | 0.116 | 0.110 | 0.97 | 0.005 | 0.009 | 3.40 |
| Ferroptosis | CP | 0.063 | 0.055 | 1.10 | 0.001 | 0.001 | 6.26 |
|  | DP | 0.061 | 0.064 | 1.16 | 0.001 | 0.001 | 5.41 |
|  | DINO | 0.048 | 0.041 | 1.04 | 0.004 | 0.008 | 3.72 |
| ICD | CP | 0.011 | 0.013 | 1.80 | 0.001 | 0.000 | 4.21 |
|  | DP | 0.017 | 0.023 | 1.80 | 0.001 | 0.001 | 6.38 |
|  | DINO | 0.011 | 0.013 | 1.69 | 0.006 | 0.015 | 3.51 |
| Necroptosis | CP | 0.154 | 0.107 | 0.06 | 0.002 | 0.002 | 3.88 |
|  | DP | 0.253 | 0.198 | 0.05 | 0.002 | 0.003 | 3.58 |
|  | DINO | 0.329 | 0.243 | -0.14 | 0.032 | 0.046 | 2.05 |
| Pyroptosis | CP | 0.237 | 0.177 | 0.45 | 0.002 | 0.003 | 4.26 |
|  | DP | 0.446 | 0.248 | -0.54 | 0.013 | 0.020 | 2.96 |
|  | DINO | 0.286 | 0.183 | 0.16 | 0.059 | 0.071 | 1.53 |

### 3.3 Single-cell density analysis reveals biological heterogeneity within apoptotic cells

Building on the previous results, we sought to explore the heterogeneity within our PCD subtypes. For this, we focused on the DINO-derived features. We selected apoptosis for further analysis due to its high variability across compounds, providing a case for studying the cellular heterogeneity.

To investigate, a new UMAP embedding was trained on the single-cell DINO features of the apoptotic cells. The embedding revealed distinct cluster formations within the morphological space (Figure 3 and Supplemental Figure S12). To further understand the emergence of these clusters, we utilized our *CellViewer* tool to examine different regions within the UMAP. Representative cells from each region were identified using a k-medoid algorithm, and their corresponding cell images were annotated directly onto the UMAP (Figure 3A). The analysis revealed that specific regions in the morphological space were associated with distinct cellular morphologies. Notably, cells in the region highlighted in the middle exhibited morphological traits typically associated with apoptosis, including shrinkage, isolation, nuclear fragmentation, and vesicle formation. Interestingly, this region contained cells originating from a diverse array of compounds (Supplemental Figure S12). Outward from this central apoptotic region, more distinct morphological patterns emerged, highlighting morphological heterogeneity within the apoptosis subtype.

**Figure 3.**
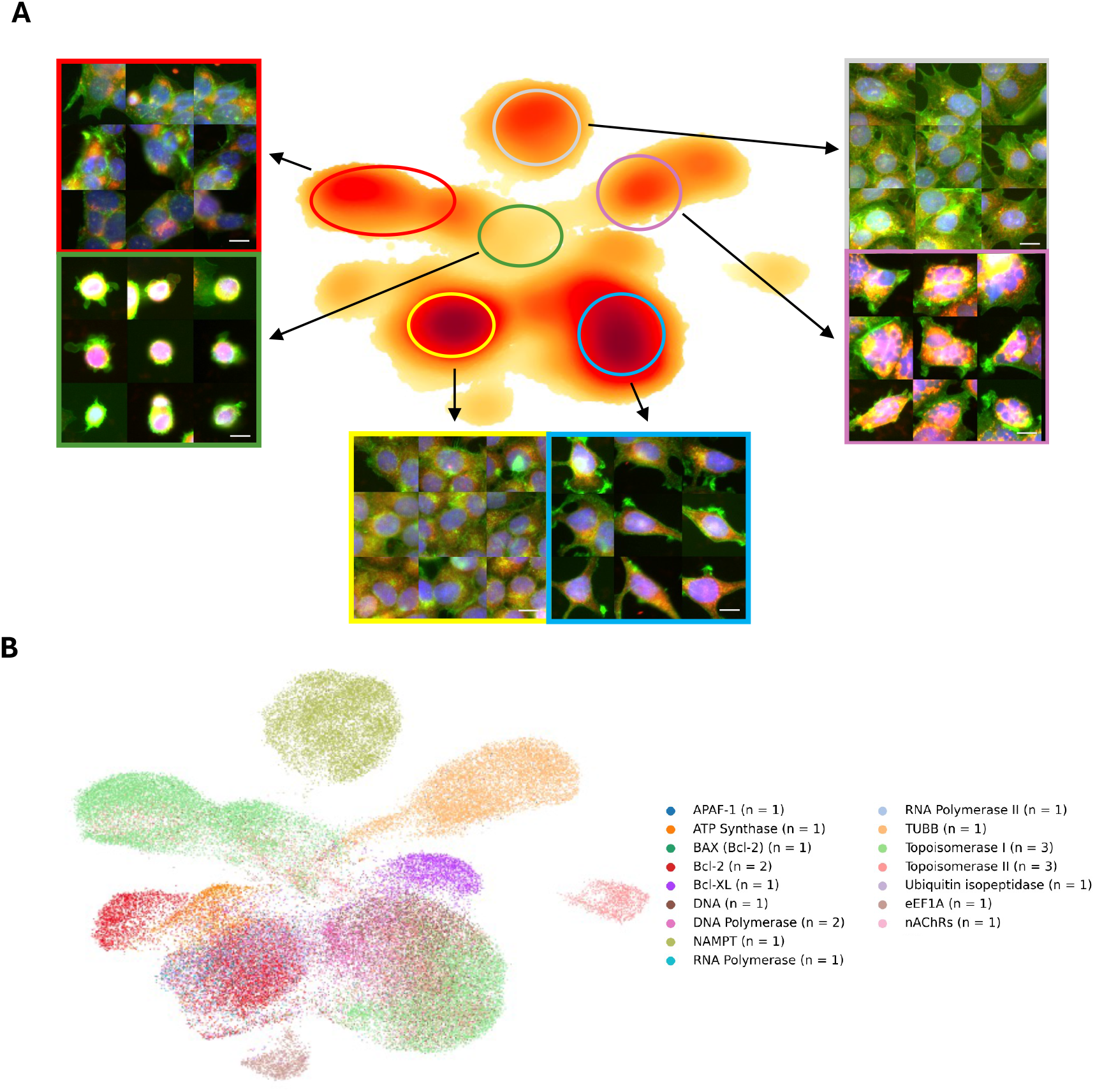
Density mapping of cells from the apoptosis subtype on DINO embedding. Scale bar corresponds to 20 µm. (A) UMAP showing the distribution of cell morphologies across treatments in the apoptosis class, with warmer colors (yellow to red) indicating higher densities. Colored circles highlight specific clusters linked to distinct phenotypes. Representative cell images adjacent to the UMAP are overlays of the Hoechst (nucleus, blue), Mitotracker (red/yellow) and Phalloidin (Cytoskeleton/ actin, green) channels. (B) Single-cell UMAP embedding of the apoptosis class colored by their annotated targets and number of unique compounds per target. Specific target, compound matching can be found in Supplemental Table S1.

Given this observed heterogeneity, we next investigated its potential origins. As outlined, cells in this experiment were treated with a wide range of compounds known to induce apoptosis. To elucidate the relationship between morphology and molecular mechanisms, compounds were next annotated with their respective targets and which were mapped onto the UMAP. This revealed a connection between distinct regions of the UMAP and specific molecular targets (Figure 3B and Supplemental Figure S13C). For instance, cells in the bottom-right region (Figure 3A) were predominantly treated with compounds targeting Topoisomerases I/II, DNA polymerases, or intercalating DNA agents (Figure 3B). The morphological characteristics in this region reflect these molecular mechanisms specifically by changes in the nucleus channel (indicated by blue color).

Analogous analyses using DeepProfiler and CellProfiler features demonstrated similar trends in morphological subspaces, albeit with less pronounced cluster formation (Supplemental Figure S13A and B, Supplemental Table S1). Overall, DINO-derived features provided the clearest differentiation of sets of cells while maintaining the most homogeneity among distinct regions in the morphological space. These findings underscore the ability of single-cell morphological features to reveal fine-grained heterogeneity within PCD subtypes and highlight the superior performance of DINO features in distinguishing regions connected to a molecular target.

### 3.4 Single-cell analysis reveal dose-dependent morphological effects

We next investigated how compound concentration influences morphological changes on single-cell level. We projected chemical perturbations at varying concentrations onto the morphological space of the corresponding cell death subtypes. More precisely, we first filtered our datasets to the respective subtypes, learned a new UMAP embedding, and subsequently projected the compound concentrations into that space and calculated their density distributions. As a showcase, we focused on the apoptosis class, which contained the highest abundance of individual compounds. We randomly selected three compounds (Cladribine, SN 38, and Topotecan) to examine their dose-dependent morphologies. We were able to detect shifts in morphology within the morphological space, driven by compound dosage (Figures 4, S14, S15). Notably, SN 38 exhibited the most prominent morphological changes, with cells shifting from a cluster in the bottom right to a region in the top left of the apoptosis-specific morphological space, based on DINO features (Figure 4B). We used the *CellViewer* tool to identify key representative cells across concentrations. Representative cells highlighted this transition, showing a progression from triangular-shaped cells with large nuclei at the lowest concentration (0.1 *µ*M) to elongated cells at the highest concentration (5.0 *µ*M) as well as cytoskeletal re-organization. A similar dose-dependent effect was observed for Cladribine (Figure 4C), where cells at the lowest concentration resembled untreated MCF-7 cells. With increasing concentrations, cells exhibited larger nuclei, stronger mitochondrial signals, and a lack of neighboring cells. However, not all compounds demonstrated pronounced changes. For instance, Topotecan showed lower morphological alterations at lower concentrations, with notable shifts only emerging at the highest tested concentration. We corroborated these observations by pair-wise e-testing between the respective compound concentrations (Figures 4D, S14D, and S15D). Our results showed significant changes in morphology between all compound concentrations. Furthermore, energy distances confirmed previous observations where SN 38 presented the most noticeable changes in morphology at higher concentrations as well as noticeable shifts between the concentrations of 0.1 and 0.3 *µ*M in Cladribine.

**Figure 4.**
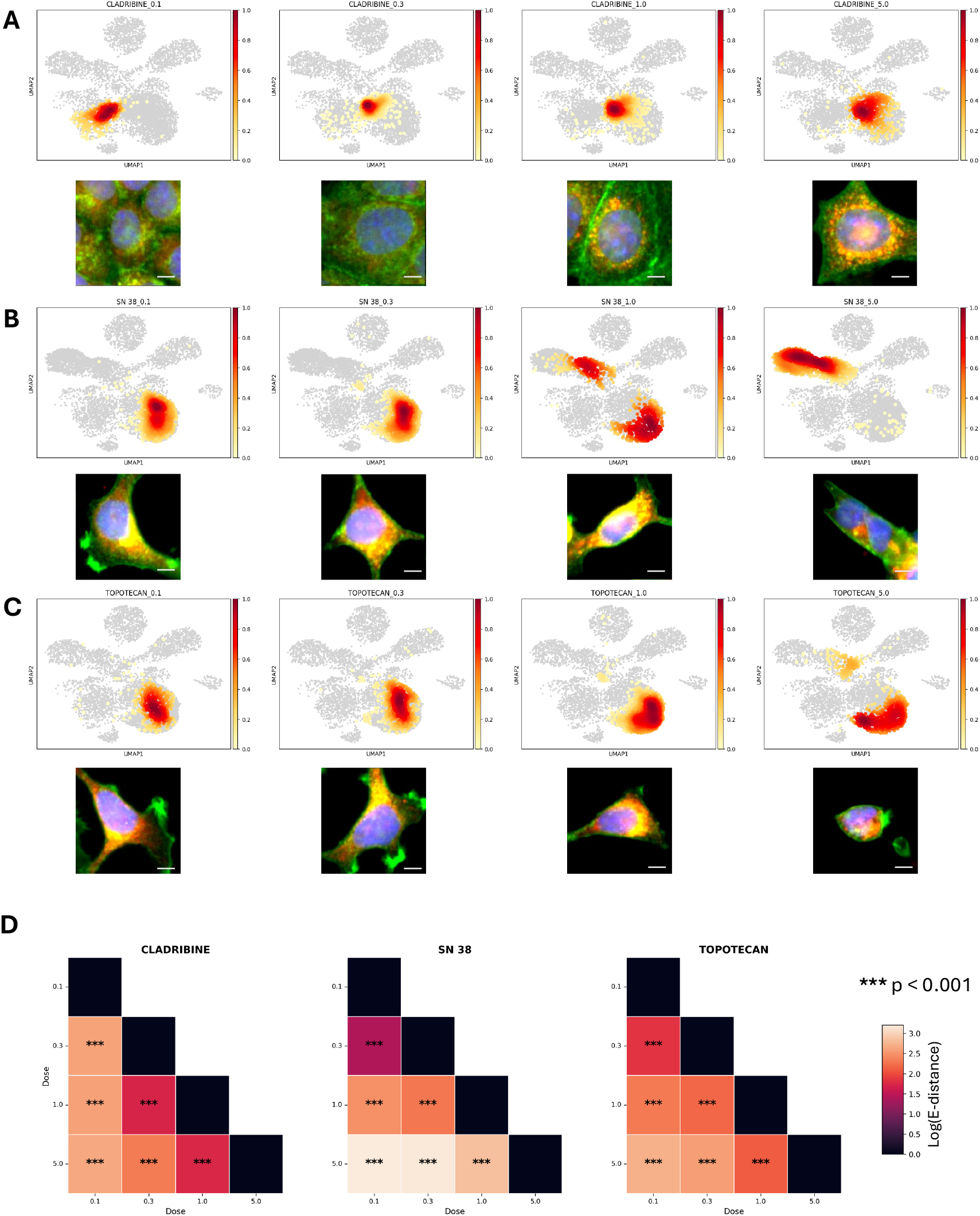
UMAP representation of single-cell morphological profiles treated with varying concentrations of compounds, accompanied by representative cell images for the apoptosis class in DINO features. Scale bar corresponds to 10 µm. Each row shows UMAP projections of cells treated with increasing concentrations (0.1, 0.3, 1.0, 3.0, and 5.0 µM). UMAP plots highlight the distribution of treated cells (colored) relative to the remaining cells in the apoptosis. Representative cell images below each UMAP depict an overlay of nuclei (blue), cytoskeleton (green), and mitochondria (red/yellow). Cells were identified with *CellViewer*. (A) UMAP projections and representative images of cells treated with Cladribine. (B) UMAP projections and representative images of cells treated with SN 38. (C) UMAP projections and representative images of cells treated with Topotecan. (D) Pair-wise energy distances (log) between concentrations (µM) for Cladribine, SN 38 and Topotecan annotated with significance level calculated via etest.

To further explore how we could use dose-response effects in our understanding of PCD, we examined features derived from the CellProfiler suite. Specific cellular features have previously been used to investigate effects relevant to cell death and the cell cycle, such as the integrated intensity of the nucleus (Matuszewski et al., 2016; Mueller et al., 2006; Olofsson et al., 2021; Roukos et al., 2015). To demonstrate its utility, we analyzed a range of compounds and their corresponding feature distributions in the normalized integrated intensity across concentrations (Supplemental Figure S16). Our findings revealed clear dose-dependent integrated DNA intensity shift, indicating a change in cell cycle such as G1 or G2 arrests, a characteristic effect known to be induced by these families of compounds. These results underscore how morphological features, such as those derived from CellProfiler, can provide quantitative insights into dose-dependent cellular responses, leveraging information provided by single cells.

### 3.5 Aggregate profiles outperform single-cells in cell death classification

After identifying distinct cell death subtypes through unsupervised analysis, we proceeded to evaluate whether single-cell morphological profiles could be utilized to accurately classify cell death subtypes. We assessed their performance via single-cell, aggregated, and majority-voted strategies to identify the most effective approaches for this classification task.

At the single-cell level, our analysis demonstrated that a bagged LightGBMXT model trained on DINO features achieved the highest classification performance, with a macro F1 score of 79.86% (Table 3) and a minimally lower performance of 79.03% macro F1 for the NeuralNetFastAI model. Comparatively, the top-performing model trained on CellProfiler features achieved a macro F1 score of 76.84%, showing a decrease in overall accuracy but comparable F1 performance. Models trained on DeepProfiler features exhibited lower performance at the single-cell level. The highest macro F1 achieved for DeepProfiler 66.12%, representing a performance drop of up to 13.74% compared to the LightGBMXT model trained on DINO features. When evaluating the classification performance on aggregated profiles, the results revealed improved metrics across all feature extraction methods. The LightGBMXT model trained on DINO features attained a macro F1 score of 88.51%, with a comparable performance to the model trained on CellProfiler features, which achieved a macro F1 score of 91.49% and the DeepProfiler model with an macro F1 score of 90.00% (Table 3).

**Table 3.**
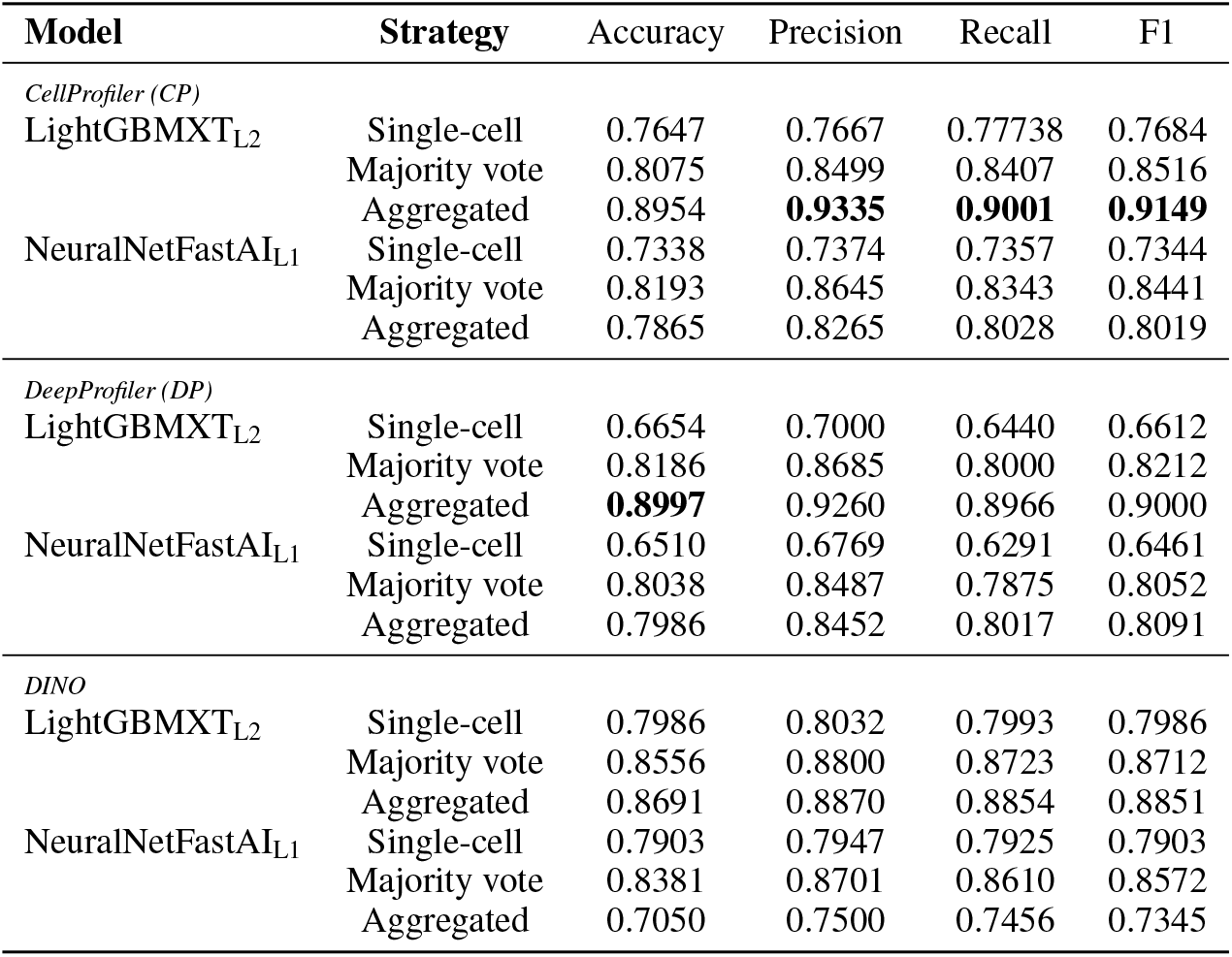
Comparison of Performance Metrics across Feature Sets with Standard, Majority Vote, and Aggregated Results. Each column depicts Accuracy, Precision, Recall, and F1 scores (all macro). Bold indicates the highest score per metric across models and strategies.

Recognizing that single-cell profiles are rarely analyzed in isolation but belong a distributions of perturbed cells, we explored an alternative strategy to enhance classification performance. We computed predictions from single-cell classifiers and aggregated these predictions using a majority-vote strategy at the site level. This approach improved the performance of our top-performing single-cell model. The LightGBMXT model trained on DINO features saw a performance boost of 7.26%, achieving an accuracy of 85.56% and a macro F1 score of 87.12% (Table 3). Similar performance deltas were observed across all remaining models, demonstrating the effectiveness of this aggregation strategy. However, despite the observed gains, none of the majority-voted models surpassed the top-performing models trained on aggregated profiles.

To further investigate model performance, we evaluated the confusion matrices for our classification models, showing the percentage of correctly predicted labels. Across all models, we observed a noticeable fraction of misclassifications between the ICD class (label 3) and apoptosis class (label 0) with up to 47.47% misclassification (Supplemental Figures S17-S19). The rate of misclassification was least pronounced in our DINO models, specifically in the single-cell models, where we observed a maximum of 21.09% misclassification between classes. As our feature extractions resulted in minimal differences in compound concentration composition and thereby train/test splitting, we further evaluated our models on an intersection of the test sets. We observed a slight increase in performance without change in the performance difference between the models as indicated by misclassification rates (Supplemental Figure S20).

Analysis of our compounds revealed structural homology between a few compounds in these classes; Etoposide (apoptosis) and Teniposide (ICD), as well as Doxorubicin/ Epirubicin (apoptosis) and Daunorubicin (ICD). We noticed similarities among their induced morphological effects, which could explain the misclassification of these specific classes (Supplemental Figure S9).

In an attempt to validate these results, we visualized the distribution of cell death classes in the UMAP spaces through density estimation (Supplemental Figure S9). We observed homogeneity among classes for both CellProfiler and DeepProfiler features (Supplemental Figure S9A and B), while DINO features exhibited significantly stronger spread, an observation we captured earlier through the k* distribution skewness (Table 2, Supplemental Figure S9C). We point out the overlap of the apoptosis and ICD classes in the UMAP spaces, further hinting at our observation of structurally similar compounds between the two classes (Supplemental Figure S9A-C).

## 4 Discussion

Effective handling of perturbations that produce little to no morphological change can be an essential tool in profiling studies. At the single-cell level, filtering out “unaffected” cells reduces noise and greatly enhances downstream detection of genuine morphological effects. In our implementation of the grit score, we noticed limitations in computing scores at the single-cell level for larger datasets, restricting its applicability in high-throughput experiments. For instance, on our cluster with 256 GB of RAM, grit scores could not be reliably computed for over 15,000 cells in one go, greatly curtailing its applicability in such experiments. To complement grit scores, we evaluated e-distance, a computationally more efficient metric suitable for large-scale single-cell datasets (Peidli et al., 2024). Our results showed that e-distance and grit scores produced comparable filtering outcomes, suggesting that either metric, or a combination, could be effectively used to identify treatments inducing significant morphological changes (Figure 1C). The choice of filtering metric ultimately depends on the experimental context and availability of computational resources, with e-distance offering scalability for single-cell data while grit scores provide robustness at the aggregated level. Additionally, we compared our results to the recently proposed mean average precision (Kalinin et al., 2025) and showed comparable results for DeepProfiler and DINO derived features (Supplemental Figure S7). For CellProfiler, we noticed a more pronounced difference in significant groups compared to etest. We leave a more thorough investigation of the effect of feature extractors on perturbation metrics open to future works.

Our unsupervised analysis illustrated that single-cell profiles offer a powerful means to study how different types of PCD emerge, revealing both subpopulation dynamics and distinct morphological characteristics (Figures 2 and 3). A notable observation across all feature extractors was the non-homogeneity of PCD subtypes. Using the apoptosis class as an example, we were able to identify distinct cellular populations within the morphological space, each associated with specific morphological traits (Figure 3). Indeed, apoptosis encompasses a wide range of intrinsic mechanisms (Dehghan et al., 2023; Kerr et al., 1972)), leading to diverse morphological effects, such as the formation of apoptotic bodies, decrease in cellular volume, enlargement of the ER, and membrane blebbing or blistering, among others (Y. Chen et al., 2024; Kerr et al., 1972), which may explain the diversity of phenotypes that we identify.

To validate the identified apoptosis subpopulations, we mapped compound targets to these regions in the morphological space. Strikingly, this revealed a clear correlation between the subpopulations and specific target annotations, suggesting that distinct molecular mechanisms drive the observed morphological diversity (Figure 3B). Compounds targeting similar pathways or molecular components clustered together within the morphological embedding, providing an additional layer of biological interpretation to the observed heterogeneity. Moreover, we demonstrated that dose-dependent changes in morphology further contribute to heterogeneity, as compounds induce distinct morphological effects only at specific concentration levels (Figure 4 and Supplemental Figures S14 and S15) and certain cell cycle stages (Supplemental Figure S16).

The choice of feature extractor plays a crucial role in shaping the observed morphological patterns. Deep learning-based feature extraction methods, such as DeepProfiler and DINO, leverage fundamentally different architectures to learn features from imaging data. DeepProfiler employs a weakly supervised approach explicitly trained to differentiate treatments (Moshkov et al., 2024), while DINO uses a self-supervised paradigm to learn representations by minimizing a loss between teacher and student networks, focusing on local and global features (Caron et al., 2021). Our results indicate that DINO tends to capture more intricate features, as evidenced by dense clustering of similar treatments in both UMAP embeddings and correlation maps (Figures 2, 3 and Supplemental Figure S8).

As deep learning based features are inherently difficult to interpret biologically, it becomes necessary to use additional approaches like our mapping of underlying imaging data to validate their relevance. To further extend on this, explainability remains a critical consideration for deep learning approaches, especially in biological and medical applications (Tanevski et al., 2022). While tools like CellProfiler provide interpretable features, neural network-based methods often require techniques like GradCAM (Selvaraju et al., 2019) or visualization of attention heads in vision transformers (Caron et al., 2021) to understand what the model is focusing on during training and inference.

However, we point to anomalies, such as the unexplained cluster in the DeepProfiler features (Supplemental Figure S10), which suggest limitations in deep-learning approaches. We hypothesized that this cluster could originate from differences in the training set (using U2OS and A549 cell lines) compared to the MCF-7 cells we used for inference here in our study (Moshkov et al., 2024). This underscores the importance of domain-specific fine-tuning when using larger pre-trained models and careful evaluation of obtained results.

Despite the strengths of single-cell profiles in uncovering subpopulation dynamics and morphological heterogeneity, several limitations must be acknowledged. While our supervised classification tasks achieved macro F1 scores of up to 79%, our single-cell classification models consistently underperformed compared to majority-voting and aggregated profile approaches. This performance gap may be attributed to an oversimplification of underlying population heterogeneity when relying solely on single-cell data and treating them as individual data points. In an attempt to consider treatments more as populations, we implemented a simple majority voting multiple-instance learning strategy, reaching similar performance levels compared to the aggregated (Table 3). Recently, more advanced modeling approaches such as optimal transport, archetype analysis, and deep sets (Baran et al., 2019; Bunne et al., 2023; Dijk et al., 2023; Dong et al., 2023) have emerged, offering valuable insights into perturbation effects and cellular responses at the same time. These methods hold promise for future addressing the challenges associated with single-cell analysis by taking underlying distributions and their changes into consideration.

In addition to the aggregation levels, smaller performance differences were observed between CellProfiler, DeepProfiler, and DINO models, with disparities of up to 13% in accuracy (Table 3). In our pipeline, we matched dataset sizes as closely as possible to minimize scaling effects, as larger datasets have been shown to improve performance by better approximating the underlying data distributions (Althnian et al., 2021; Dosovitskiy et al., 2021). Additionally, we assessed results on an intersection of our test sets ensuring equality between predicted sites. Here, we observed similar results, ensuring that model performance was not a consequence of differences in our data splitting approach (Supplemental Figure S20). Since aggregated results across feature extractors were consistent, we attribute the differences to the underlying architectures and specifically to DeepProfiler’s training paradigm and the detected anomaly in its feature space (Supplemental Figure S10). Overall however, our findings are consistent with earlier studies showing that deep-learning–derived features (specifically from DINO networks) perform on par with or exceed CellProfiler features in classification tasks (Gupta et al., 2024; V. Kim et al., 2024; Moshkov et al., 2024). Our results underscore the importance of considering both training data size and feature extraction methodology when designing models for morphological profiling. They further highlight the potential trade-offs between capturing detailed biological variability and achieving state-of-the-art performance in classification tasks. Future works could explore strategies to enhance DINO’s generalization capabilities, such as incorporating additional pre-training data, fine-tuning on class-specific labels, or using hybrid approaches that combine supervised and self-supervised learning paradigms or contrastive learning frameworks with other data modalities as seen in (Fradkin et al., 2024; Sanchez-Fernandez et al., 2023).

From a computational perspective, the scalability of single-cell morphological profiling remains a challenge. High-throughput analyses require efficient pre-processing and filtering strategies to ensure reliable downstream analyses. In this study, deep learning-based feature extraction methods not only matched or outperformed traditional tools like CellProfiler but also required fewer features to achieve these results. CellProfiler with extensive feature seleciton resulted in 1,400 hand-crafted features, while DeepProfiler and DINO had 672 and 384 features, respectively, to deliver comparable or better outcomes. Additionally, inference of features was obtainable in substantially less time with 2 hours per plate in DeepProfiler and 1 hour for DINO, both running on a single NVIDIA GTX 3090. As CellProfiler is, as of now, not capable of running feature extraction on GPUs, the inference times approached > 4 hours per plate. Overall, computational cost reductions significantly alleviate computational bottlenecks, making deep learning frameworks suitable for large-scale, high-throughput applications.

Single-cell morphological data can offer invaluable insights into cellular heterogeneity and subpopulation dynamics, complementing aggregated profiles for exploratory and hypothesis-driven analyses. However, in this work, we showed that for effective use, rigorous methodological choices are required, including feature extraction strategies and filtering metrics, tailored to the specific biological context. By integrating computationally efficient and scalable analysis methods, single-cell morphological profiling can be further refined to address its current limitations and unlock its full potential for studying complex cellular processes. Here, we presented cell death as a use case to show the utility of single-cell morphological profiling and informing drug discovery pipelines.

## 5 Authors contributions

BF: Design, methodology, analysis, visualization, interpretation, manuscript writing; DH: Methodology, analysis; PB: Methodology, analysis; EB: Methodology, interpretation; PG: Cell Painting experiments; MJ: Cell Painting experiments; PH: Methodology, interpretation; DR: Software development; JR: Cell Painting experiments, interpretation; JCP: Design, conceptualization, interpretation, manuscript writing, supervision; OS: Design, conceptualization, interpretation, manuscript writing, supervision; All of the authors reviewed, edited, and contributed to discussions on the manuscript and approved the final version of the manuscript.

## 7 Acknowledgements

Compounds were provided by Beactica Therapeutics. The computations were enabled by resources provided by the National Academic Infrastructure for Supercomputing in Sweden (NAISS), partially funded by the Swedish Research Council through grant agreement no. 2022-06725, resources provided by Uppsala University at UPPMAX, and the Berzelius resource provided by the Knut and Alice Wallenberg Foundation at the National Supercomputer Centre. The Authors acknowledge the support from the Chemical Biology Consortium Sweden (CBCS), the node at Uppsala University. CBCS is a national research infrastructure funded by the Swedish Research Council (dr.nr.2021-00179) and SciLifeLab.

## References

Aleo, Emanuela, Clare J Henderson, Alessandra Fontanini, Barbara Solazzo, and Claudio Brancolini (2006). “Identification of new compounds that trigger apoptosome-independent caspase activation and apoptosis”. 66: 18, pp. 9235–9244. DOI: 10.1158/0008-5472.CAN-06-0702.

Althnian, Alhanoof, Duaa AlSaeed, Heyam Al-Baity, Amani Samha, Alanoud Bin Dris, Najla Alzakari, Afnan Abou Elwafa, and Heba Kurdi (Jan. 2021). “Impact of Dataset Size on Classification Performance: An Empirical Evaluation in the Medical Domain”. Applied Sciences, 11: 2, p. 796.

Amiri, Mina, Ommoleila Molavi, Shahnaz Sabetkam, Sevda Jafari, and Soheila Montazersaheb (2023). “Stimulators of immunogenic cell death for cancer therapy: focusing on natural compounds”.23: 1, pp. 200–200. DOI: 10.1186/s12935-023-03058-7.

Bai, Bingjun, Lina Shan, Jianhong Wang, Jinhui Hu, Wenqian Zheng, Yiming Lv, Kangke Chen, Dengyong Xu, and Hongbo Zhu (2020). “Small molecule 2,3-DCPE induces S phase arrest by activating the ATM/ATR-Chk1-Cdc25A signaling pathway in DLD-1 colon cancer cells”. 20: 6, pp. 294–294. DOI: 10.3892/ol.2020.12157.

Balakrishnan, Kumudha, Jan A Burger, William G Wierda, and Varsha Gandhi (2009). “AT-101 induces apoptosis in CLL B cells and overcomes stromal cell-mediated Mcl-1 induction and drug resistance”. 113: 1, pp. 149–153. DOI: 10.1182/blood-2008-02-138560.

Baran, Yael, Akhiad Bercovich, Arnau Sebe-Pedros, Yaniv Lubling, Amir Giladi, Elad Chomsky, Zohar Meir, Michael Hoichman, Aviezer Lifshitz, and Amos Tanay (Oct. 2019). “MetaCell: analysis of single-cell RNA-seq data using K-nn graph partitions”. Genome Biology, 20: 1, p. 206.

Baraz, Rana, Adam Cisterne, Philip O Saunders, John Hewson, Marilyn Thien, Jocelyn Weiss, Jordan Basnett, Kenneth F Bradstock, and Linda J Bendall (2014). “mTOR inhibition by everolimus in childhood acute lymphoblastic leukemia induces caspase-independent cell death”. 9: 7, e102494–e102494. DOI: 10.1371/journal.pone.0102494.

Barry, M A, J E Reynolds, and A Eastman (1993). “Etoposide-induced apoptosis in human HL-60 cells is associated with intracellular acidification”. 53: 10 Suppl, pp. 2349–2357.

Blatt, Neal B, Anthony E Boitano, Costas A Lyssiotis, Jr Opipari Anthony W, and Gary D Glick (2008). “Bz-423 superoxide signals apoptosis via selective activation of JNK, Bak, and Bax”. 45: 9, pp. 1232–1242. DOI: 10.1016/j.freeradbiomed.2008.07.022.

Blatt, Neal B, Anthony E Boitano, Costas A Lyssiotis, Jr Opipari Anthony W, and Gary D Glick (2009). “Bz-423 superoxide signals B cell apoptosis via Mcl-1, Bak, and Bax”. 78: 8, pp. 966–973. DOI: 10.1016/j.bcp.2009.05.025.

Bownik, A, A Rymuszka, A Sierosławska, and T Skowroński (2012). “Anatoxin-a induces apoptosis of leukocytes and decreases the proliferative ability of lymphocytes of common carp (Cyprinus carpio L.) in vitro”. 15: 3, pp. 531–535. DOI: 10.2478/v10181-012-0082-7.

Bray, Mark-Anthony, Shantanu Singh, Han Han, Chadwick T. Davis, Blake Borgeson, Cathy Hartland, Maria Kost-Alimova, Sigrun M. Gustafsdottir, Christopher C. Gibson, and Anne E. Carpenter (Sept. 2016). “Cell Painting, a high-content image-based assay for morphological profiling using multiplexed fluorescent dyes”. Nature Protocols, 11: 9, pp. 1757–1774.

Breiman, Leo (Oct. 2001). “Random Forests”. Machine Learning, 45: 1, pp. 5–32.

Bugaut, Hélène, Mélanie Bruchard, Hélène Berger, Valentin Derangère, Ludivine Odoul, Romain Euvrard, Sylvain Ladoire, Fanny Chalmin, Frédérique Végran, Cédric Rébé, et al. (2013). “Bleomycin exerts ambivalent antitumor immune effect by triggering both immunogenic cell death and proliferation of regulatory T cells”. 8: 6, e65181–e65181. DOI: 10.1371/journal.pone.0065181.

Bunne, Charlotte, Stefan G. Stark, Gabriele Gut, Jacobo Sarabia del Castillo, Mitch Levesque, Kjong-Van Lehmann, Lucas Pelkmans, Andreas Krause, and Gunnar Rätsch (Nov. 2023). “Learning single-cell perturbation responses using neural optimal transport”. Nature Methods, 20: 11, pp. 1759–1768.

Cai, Ming, Jingquan He, Jian Xiong, Li Wei Rachel Tay, Ziqing Wang, Colin Rog, Jingshu Wang, Yizhao Xie, Guobin Wang, Yoshiko Banno, et al. (Nov. 2016). “Phospholipase D1-regulated autophagy supplies free fatty acids to counter nutrient stress in cancer cells”. Cell Death & Disease, 7: 11, e2448–e2448.

Caicedo, Juan C, Sam Cooper, Florian Heigwer, Scott Warchal, Peng Qiu, Csaba Molnar, Aliaksei S Vasilevich, Joseph D Barry, Harmanjit Singh Bansal, Oren Kraus, et al. (Sept. 2017). “Data-analysis strategies for image-based cell profiling”. Nature Methods, 14: 9, pp. 849–863.

Caron, Mathilde, Hugo Touvron, Ishan Misra, Hervé Jégou, Julien Mairal, Piotr Bojanowski, and Armand Joulin (May 2021). Emerging Properties in Self-Supervised Vision Transformers. arXiv, 2021. DOI: 10.48550/arXiv.2104.14294.

Carpenter, Anne E., Thouis R. Jones, Michael R. Lamprecht, Colin Clarke, In Han Kang, Ola Friman, David A. Guertin, Joo Han Chang, Robert A. Lindquist, Jason Moffat, et al. (Oct. 2006). “CellProfiler: image analysis software for identifying and quantifying cell phenotypes”. Genome Biology, 7: 10, R100.

Caserini, C, G Pratesi, M Tortoreto, B Bedogné, N Carenini, R Supino, P Perego, S C Righetti, and F Zunino (1997). “Apoptosis as a determinant of tumor sensitivity to topotecan in human ovarian tumors: preclinical in vitro/in vivo studies”. 3: 6, pp. 955–961.

Chabanon, Roman M, Gareth Muirhead, Dragomir B Krastev, Julien Adam, Daphné Morel, Marlène Garrido, Andrew Lamb, Clémence Hénon, Nicolas Dorvault, Mathieu Rouanne, et al. (2019). “PARP inhibition enhances tumor cell-intrinsic immunity in ERCC1-deficient non-small cell lung cancer”. 129: 3, pp. 1211–1228. DOI: 10.1172/JCI123319.

Chen, Daniel, Tempest Plott, Michael Wiest, Will Van Trump, Ben Komalo, Dat Nguyen, Charlie Marsh, Jarred Heinrich, Colin J. Fuller, Lauren Nicolaisen, et al. (Dec. 2024). “A combined AI and cell biology approach surfaces targets and mechanistically distinct Inflammasome inhibitors”. iScience, 27: 12, p. 111404.

Chen, Tianqi and Carlos Guestrin (Aug. 2016). “XGBoost: A Scalable Tree Boosting System” Proceedings of the 22nd ACM SIGKDD International Conference on Knowledge Discovery and Data Mining, pp. 785–794.

Chen, Yao, Xiaohua Li, Minfeng Yang, and Song-Bai Liu (May 2024). “Research progress on morphology and mechanism of programmed cell death”. Cell Death & Disease, 15: 5, pp. 1–13.

Conrad, Marcus, José Pedro Friedmann Angeli, Peter Vandenabeele, and Brent R. Stockwell (May 2016). “Regulated necrosis: disease relevance and therapeutic opportunities”. Nature Reviews Drug Discovery, 15: 5, pp. 348–366.

Cotto-Rios, Xiomaris M and Evripidis Gavathiotis (2016). “Chemical genetics: Unraveling cell death mysteries”. 12: 7, pp. 470–471. DOI: 10.1038/nchembio.2110.

Crazzolara, Roman, Kenneth F Bradstock, and Linda J Bendall (2009). “RAD001 (Everolimus) induces autophagy in acute lymphoblastic leukemia”. 5: 5, pp. 727–728. DOI: 10.4161/auto.5.5.8507.

Criollo, A, M C Maiuri, E Tasdemir, I Vitale, A A Fiebig, D Andrews, J Molgó, J Díaz, S Lavandero, F Harper, et al. (2007). “Regulation of autophagy by the inositol trisphosphate receptor”. 14: 5, pp. 1029–1039. DOI: 10.1038/sj.cdd.4402099.

Dehghan, Sepehr, Nasim Kheshtchin, Shaghayegh Hassannezhad, and Maryam Soleimani (Dec. 2023). “Cell death classification: A new insight based on molecular mechanisms”. Experimental Cell Research, 433: 2, p. 113860.

Dijk, Robert van, John Arevalo, Mehrtash Babadi, Anne E. Carpenter, and Shantanu Singh (Nov. 2023). Capturing cell heterogeneity in representations of cell populations for image-based profiling using contrastive learning. bioRxiv, 2023. DOI: 10.1101/2023.11.14.567038.

Dixon, Scott J, Kathryn M Lemberg, Michael R Lamprecht, Rachid Skouta, Eleina M Zaitsev, Caroline E Gleason, Darpan N Patel, Andras J Bauer, Alexandra M Cantley, Wan Seok Yang, et al. (2012). “Ferroptosis: an iron-dependent form of nonapoptotic cell death”. 149: 5, pp. 1060–1072. DOI: 10.1016/j.cell.2012.03.042.

Dong, Mingze, Bao Wang, Jessica Wei, Antonio H. de O. Fonseca, Curtis J. Perry, Alexander Frey, Feriel Ouerghi, Ellen F. Foxman, Jeffrey J. Ishizuka, Rahul M. Dhodapkar, et al. (Nov. 2023). “Causal identification of single-cell experimental perturbation effects with CINEMA-OT”. Nature Methods, 20: 11, pp. 1769–1779.

Dorogush, Anna Veronika, Vasily Ershov, and Andrey Gulin (Oct. 2018). CatBoost: gradient boosting with categorical features support. arXiv, 2018. DOI: 10.48550/arXiv.1810.11363.

Doron, Michael, Théo Moutakanni, Zitong S. Chen, Nikita Moshkov, Mathilde Caron, Hugo Touvron, Piotr Bojanowski, Wolfgang M. Pernice, and Juan C. Caicedo (June 2023). Unbiased single-cell morphology with self-supervised vision transformers. bioRxiv, 2023. DOI: 10.1101/2023.06.16.545359.

Dosovitskiy, Alexey, Lucas Beyer, Alexander Kolesnikov, Dirk Weissenborn, Xiaohua Zhai, Thomas Unterthiner, Mostafa Dehghani, Matthias Minderer, Georg Heigold, Sylvain Gelly, et al. (June 2021). An Image is Worth 16×16 Words: Transformers for Image Recognition at Scale. arXiv, 2021. DOI: 10.48550/arXiv.2010.11929.

Dumont, Patrick, Laurent Ingrassia, Sébastien Rouzeau, Fabrice Ribaucour, Stéphanie Thomas, Isabelle Roland, Francis Darro, Florence Lefranc, and Robert Kiss (2007). “The Amaryllidaceae isocarbostyril narciclasine induces apoptosis by activation of the death receptor and/or mitochondrial pathways in cancer cells but not in normal fibroblasts”. 9: 9, pp. 766–776. DOI: 10.1593/neo.07535.

Erickson, Nick, Jonas Mueller, Alexander Shirkov, Hang Zhang, Pedro Larroy, Mu Li, and Alexander Smola (Mar. 2020). AutoGluon-Tabular: Robust and Accurate AutoML for Structured Data. arXiv, 2020. DOI: 10.48550/arXiv.2003.06505.

Fan, Shuangbo, Qian Xu, Liang Wang, Yulin Wan, and Sheng Qiu (2020). “SMBA1, a Bax Activator, Induces Cell Cycle Arrest and Apoptosis in Malignant Glioma Cells”. 105: 3–4, pp. 164–172. DOI: 10.1159/000500292.

Fleming, Angeleen, Takeshi Noda, Tamotsu Yoshimori, and David C Rubinsztein (2011). “Chemical modulators of autophagy as biological probes and potential therapeutics”. 7: 1, pp. 9–17. DOI: 10.1038/nchembio.500.

Foti, Carmela, Cristina Florean, Antonio Pezzutto, Paola Roncaglia, Andrea Tomasella, Stefano Gustincich, and Claudio Brancolini (2009). “Characterization of caspase-dependent and caspase-independent deaths in glioblastoma cells treated with inhibitors of the ubiquitin-proteasome system”. 8: 11, pp. 3140–3150. DOI: 10.1158/1535-7163.MCT-09-0431.

Fradkin, Philip, Puria Azadi, Karush Suri, Frederik Wenkel, Ali Bashashati, Maciej Sypetkowski, and Dominique Beaini (Sept. 2024). How Molecules Impact Cells: Unlocking Contrastive PhenoMolecular Retrieval. arXiv, 2024. DOI: 10.48550/arXiv.2409.08302.

Francisco Rodríguez, María Andreína, Jordi Carreras Puigvert, and Ola Spjuth (Dec. 2023). “Designing microplate layouts using artificial intelligence”. Artificial Intelligence in the Life Sciences, 3: p. 100073.

Gaul, Leander, Sonja Mandl-Weber, Philipp Baumann, Bertold Emmerich, and Ralf Schmidmaier (2008). “Bendamustine induces G2 cell cycle arrest and apoptosis in myeloma cells: the role of ATM-Chk2-Cdc25A and ATM-p53-p21-pathways”. 134: 2, pp. 245–253. DOI: 10.1007/s00432-007-0278-x.

Gautam, Swetlana, Priyanka Singh, Manjari Singh, Subhadeep Roy, Jitendra K Rawat, Rajnish K Yadav, Uma Devi, Pushpraj S Gupta, Shubhini A Saraf, and Gaurav Kaithwas (2018). “Rifaximin, a pregnane X receptor (PXR) activator regulates apoptosis in a murine model of breast cancer”. 8: 7, pp. 3512–3521. DOI: 10.1039/c7ra09689e.

Gehrke, Iris, Eric D J Bouchard, Sara Beiggi, Armando G Poeppl, James B Johnston, Spencer B Gibson, and Versha Banerji (2014). “On-target effect of FK866, a nicotinamide phosphoribosyl transferase inhibitor, by apoptosis-mediated death in chronic lymphocytic leukemia cells”. 20: 18, pp. 4861–4872. DOI: 10.1158/1078-0432.CCR-14-0624.

Glynn, J M, T G Cotter, and D R Green (1992). “Apoptosis induced by Actinomycin D, Camptothecin or Aphidicolin can occur in all phases of the cell cycle”. 20: 1, 84S–84S. DOI: 10.1042/bst020084s.

Guo, Wei, Shuhong Wu, Li Wang, Xiaoli Wei, Xiaoying Liu, Ji Wang, Zhimin Lu, Melinda Hollingshead, and Bingliang Fang (2011). “Antitumor activity of a novel oncrasin analogue is mediated by JNK activation and STAT3 inhibition”. 6: 12, e28487–e28487. DOI: 10.1371/journal.pone.0028487.

Gupta, Ankit, Zoe Wefers, Konstantin Kahnert, Jan N. Hansen, Will Leineweber, Anthony Cesnik, Dan Lu, Ulrika Axelsson, Frederic Ballllosera Navarro, Theofanis Karaletsos, et al. (Dec. 2024). SubCell: Vision foundation models for microscopy capture single-cell biology. bioRxiv, 2024. DOI: 10.1101/2024.12.06.627299.

Hafner-Bratkovič, Iva, Petra Sušjan, Duško Lainšček, Ana Tapia-Abellán, Kosta Cerović, Lucija Kadunc, Diego Angosto-Bazarra, Pablo Pelegrin, and Roman Jerala (2018). “NLRP3 lacking the leucine-rich repeat domain can be fully activated via the canonical inflammasome pathway”. 9: 1, pp. 5182–5182. DOI: 10.1038/s41467-018-07573-4.

Harrison, Philip John, Ankit Gupta, Jonne Rietdijk, Håkan Wieslander, Jordi Carreras-Puigvert, Polina Georgiev, Carolina Wählby, Ola Spjuth, and Ida-Maria Sintorn (July 2023). “Evaluating the utility of brightfield image data for mechanism of action prediction”. PLOS Computational Biology, 19: 7, e1011323.

Haslum, Johan Fredin, Charles Lardeau, Johan Karlsson, Riku Turkki, Karl-Johan Leuchowius, Kevin Smith, and Erik Müllers (Apr. 2023). Cell Painting-based bioactivity prediction boosts high-throughput screening hit-rates and compound diversity. bioRxiv, 2023. DOI: 10.1101/2023.04.03.535328.

Hasmann, Max and Isabel Schemainda (2003). “FK866, a highly specific noncompetitive inhibitor of nicotinamide phosphoribosyltransferase, represents a novel mechanism for induction of tumor cell apoptosis”. 63: 21, pp. 7436–7442.

He, Kaiming, Xinlei Chen, Saining Xie, Yanghao Li, Piotr Dollár, and Ross Girshick (Dec. 2021). Masked Autoencoders Are Scalable Vision Learners. arXiv, 2021. DOI: 10.48550/arXiv.2111.06377.

He, Wan-ting, Haoqiang Wan, Lichen Hu, Pengda Chen, Xin Wang, Zhe Huang, Zhang-Hua Yang, Chuan-Qi Zhong, and Jiahuai Han (2015). “Gasdermin D is an executor of pyroptosis and required for interleukin-1 secretion”. 25: 12, pp. 1285–1298. DOI: 10.1038/cr.2015.139.

Ho, Tik-Shun, Yuk-Ping Ho, Wing-Yin Wong, Lawrence Chi-Ming Chiu, Yum-Shing Wong, and Vincent Eng-Choon Ooi (2007). “Fatty acid synthase inhibitors cerulenin and C75 retard growth and induce caspase-dependent apoptosis in human melanoma A-375 cells”. 61: 9, pp. 578–587. DOI: 10.1016/j.biopha.2007.08.020.

Huang, Shiu-Wen, I-Tsu Chyuan, Ching Shiue, Meng-Chieh Yu, Ya-Fen Hsu, and Ming-Jen Hsu (2020). “Lovastatin-mediated MCF-7 cancer cell death involves LKB1-AMPK-p38MAPK-p53-survivin signalling cascade”. 24: 2, pp. 1822–1836. DOI: 10.1111/jcmm.14879.

Huang, Tzu-Ching, Pu-Rong Chiu, Wen-Tsan Chang, Bau-Shan Hsieh, Yu-Ci Huang, Hsiao-Ling Cheng, Li-Wen Huang, Yu-Chen Hu, and Kee-Lung Chang (2018). “Epirubicin induces apoptosis in osteoblasts through death-receptor and mitochondrial pathways”. 23: 3–4, pp. 226–236. DOI: 10.1007/s10495-018-1450-2.

Humeau, Juliette, Allan Sauvat, Giulia Cerrato, Wei Xie, Friedemann Loos, Francesca Iannantuoni, Lucillia Bezu, Sarah Lévesque, Juliette Paillet, Jonathan Pol, et al. (2020). “Inhibition of transcription by dactinomycin reveals a new characteristic of immunogenic cell stress”. 12: 5, e11622–e11622. DOI: 10.15252/emmm.201911622.

Ingrassia, Laurent, Florence Lefranc, Janique Dewelle, Laurent Pottier, Véronique Mathieu, Sabine Spiegl-Kreinecker, Sébastien Sauvage, Mohamed El Yazidi, Mischaël Dehoux, Walter Berger, et al. (2009). “Structure-activity relationship analysis of novel derivatives of narciclasine (an Amaryllidaceae isocarbostyril derivative) as potential anticancer agents”. 52: 4, pp. 1100–1114. DOI: 10.1021/jm8013585.

Johnson, N, T T Ng, and J M Parkin (1997). “Camptothecin causes cell cycle perturbations within T-lymphoblastoid cells followed by dose dependent induction of apoptosis”. 21: 10, pp. 961–972. DOI: 10.1016/s0145-2126(97)00077-5.

Johnston, James B (2011). “Mechanism of action of pentostatin and cladribine in hairy cell leukemia”. 52 Suppl 2: pp. 43–45. DOI: 10.3109/10428194.2011.570394.

Kalinin, Alexandr A., John Arevalo, Erik Serrano, Loan Vulliard, Hillary Tsang, Michael Bornholdt, Alán F. Muñoz, Suganya Sivagurunathan, Bartek Rajwa, Anne E. Carpenter, et al. (Mar. 2025). A versatile information retrieval framework for evaluating profile strength and similarity. bioRxiv, 2025. DOI: 10.1101/2024.04.01.587631.

Kasibhatla, Shailaja, Vijay Baichwal, Sui Xiong Cai, Bruce Roth, Ira Skvortsova, Sergej Skvortsov, Peter Lukas, Nicole M English, Nilantha Sirisoma, John Drewe, et al. (2007). “MPC-6827: a small-molecule inhibitor of microtubule formation that is not a substrate for multidrug resistance pumps”. 67: 12, pp. 5865–5871. DOI: 10.1158/0008-5472.CAN-07-0127.

Kerr, J. F. R., A. H. Wyllie, and A. R. Currie (Aug. 1972). “Apoptosis: A Basic Biological Phenomenon with Wideranging Implications in Tissue Kinetics”. British Journal of Cancer, 26: 4, pp. 239–257.

Khedri, Azam, Shahnaz Khaghani, Alireza Kheirollah, Hossein Babaahmadi-Rezaei, Amir Shadboorestan, Mohammad Zangooei, Hajar Shokri Afra, Reza Meshkani, and Mohammad Hossein Ghahremani (2019). “Signaling Crosstalk of FHIT, p53, and p38 in etoposide-induced apoptosis in MCF-7 cells”. 120: 6, pp. 9125–9137. DOI: 10.1002/jcb.28188.

Kim, Seung-Hee and Kyung-Chul Choi (2013). “Anti-cancer Effect and Underlying Mechanism(s) of Kaempferol, a Phytoestrogen, on the Regulation of Apoptosis in Diverse Cancer Cell Models”. 29: 4, pp. 229–234. DOI: 10.5487/TR.2013.29.4.229.

Kim, Vladislav, Nikolaos Adaloglou, Marc Osterland, Flavio M. Morelli, Marah Halawa, Tim König, David Gnutt, and Paula A. Marin Zapata (Jan. 2024). Self-supervision advances morphological profiling by unlocking powerful image representations. bioRxiv, 2024. DOI: 10.1101/2023.04.28.538691.

Kline, Michael P, S Vincent Rajkumar, Michael M Timm, Teresa K Kimlinger, Jessica L Haug, John A Lust, Philip R Greipp, and Shaji Kumar (2008). “R-(−)-gossypol (AT-101) activates programmed cell death in multiple myeloma cells”. 36: 5, pp. 568–576. DOI: 10.1016/j.exphem.2008.01.003.

Korsunsky, Ilya, Nghia Millard, Jean Fan, Kamil Slowikowski, Fan Zhang, Kevin Wei, Yuriy Baglaenko, Michael Brenner, Po-ru Loh, and Soumya Raychaudhuri (Dec. 2019). “Fast, sensitive and accurate integration of single-cell data with Harmony”. Nature Methods, 16: 12, pp. 1289–1296.

Kotyan, Shashank, Tatsuya Ueda, and Danilo Vasconcellos Vargas (Aug. 2024). k* Distribution: Evaluating the Latent Space of Deep Neural Networks using Local Neighborhood Analysis. arXiv, 2024. DOI: 10.48550/arXiv.2312.04024.

Kraus, Oren, Kian Kenyon-Dean, Saber Saberian, Maryam Fallah, Peter McLean, Jess Leung, Vasudev Sharma, Ayla Khan, Jia Balakrishnan, Safiye Celik, et al. (Apr. 2024). Masked Autoencoders for Microscopy are Scalable Learners of Cellular Biology. arXiv, 2024. DOI: 10.48550/arXiv.2404.10242.

Kroemer, Guido, Lorenzo Galluzzi, Oliver Kepp, and Laurence Zitvogel (Mar. 2013). “Immunogenic Cell Death in Cancer Therapy”. Annual Review of Immunology, 31: Volume 31, 2013, pp. 51–72.

Lachaier, Emma, Christophe Louandre, Corinne Godin, Zuzana Saidak, Maxime Baert, Momar Diouf, Bruno Chauffert, and Antoine Galmiche (2014). “Sorafenib induces ferroptosis in human cancer cell lines originating from different solid tumors”. 34: 11, pp. 6417–6422.

Lakshmana Rao, P V, R Bhattacharya, Nidhi Gupta, M M Parida, A S B Bhaskar, and Rupa Dubey (2002). “Involvement of caspase and reactive oxygen species in cyanobacterial toxin anatoxin-a-induced cytotoxicity and apoptosis in rat thymocytes and Vero cells”. 76: 4, pp. 227–235. DOI: 10.1007/s00204-002-0330-1.

Leoni, Lorenzo M, Brandi Bailey, Jack Reifert, Heather H Bendall, Robert W Zeller, Jacques Corbeil, Gary Elliott, and Christina C Niemeyer (2008). “Bendamustine (Treanda) displays a distinct pattern of cytotoxicity and unique mechanistic features compared with other alkylating agents”. 14: 1, pp. 309–317. DOI: 10.1158/1078-0432.CCR-07-1061.

Li, Kun, Yihang Gong, Dongbo Qiu, Hui Tang, Jian Zhang, Zenan Yuan, Yingqi Huang, Yunfei Qin, Linsen Ye, and Yang Yang (2022). “Hyperbaric oxygen facilitates teniposide-induced cGAS-STING activation to enhance the antitumor efficacy of PD-1 antibody in HCC”. 10: 8, e004006. DOI: 10.1136/jitc-2021-004006.

Lippincott, Michael J., Jenna Tomkinson, Dave Bunten, Milad Mohammadi, Johanna Kastl, Johannes Knop, Ralf Schwandner, Jiamin Huang, Grant Ongo, Nathaniel Robichaud, et al. (Apr. 2025). “A morphology and secretome map of pyroptosis”. Molecular Biology of the Cell, mbc.E25–03–0119.

Liu, Junyu, Chenge Lou, Chenxiao Zhen, Yijia Wang, Peng Shang, and Huanhuan Lv (2022). “Iron plays a role in sulfasalazine-induced ferroptosis with autophagic flux blockage in K7M2 osteosarcoma cells”. 14: 5, mfac027. DOI: 10.1093/mtomcs/mfac027.

Liu, Ruyuan, Huanyu Cui, D. Li, Xuefeng Guo, Zhengbao Zhang, Shengkui Tan, and Xiaonian Zhu (2024). “Roles and Mechanisms of Ferroptosis in Sorafenib Resistance for Hepatocellular Carcinoma”. 11: pp. 2493–2504. DOI: 10.2147/JHC.S500084.

Liu, Tiancong, Xun Sun, and Zhiwei Cao (2019). “Shikonin-induced necroptosis in nasopharyngeal carcinoma cells via ROS overproduction and upregulation of RIPK1/RIPK3/MLKL expression”. 12: pp. 2605–2614. DOI: 10.2147/OTT.S200740.

Lotfollahi, Mohammad, Anna Klimovskaia Susmelj, Carlo De Donno, Leon Hetzel, Yuge Ji, Ignacio L Ibarra, Sanjay R Srivatsan, Mohsen Naghipourfar, Riza M Daza, Beth Martin, et al. (June 2023). “Predicting cellular responses to complex perturbations in high-throughput screens”. Molecular Systems Biology, 19: 6, e11517.

Martins, I., O. Kepp, F. Schlemmer, S. Adjemian, M. Tailler, S. Shen, M. Michaud, L. Menger, A. Gdoura, N. Tajeddine, et al. (Mar. 2011). “Restoration of the immunogenicity of cisplatin-induced cancer cell death by endoplasmic reticulum stress”. Oncogene, 30: 10, pp. 1147–1158.

Marzo, I, P Pérez-Galán, P Giraldo, D Rubio-Félix, A Anel, and J Naval (2001). “Cladribine induces apoptosis in human leukaemia cells by caspase-dependent and -independent pathways acting on mitochondria”. 359: Pt 3, pp. 537–546. DOI: 10.1042/0264-6021:3590537.

Matuszewski, Damian J., Carolina Wählby, Jordi Carreras Puigvert, and Ida-Maria Sintorn (Mar. 2016). “Population-Profiler: A Tool for Population Analysis and Visualization of Image-Based Cell Screening Data”. PLOS ONE, 11: 3, e0151554.

Moore, R Y (1993). “Principles of synaptic transmission”. 695: pp. 1–9. DOI: 10.1111/j.1749-6632.1993.tb23018.x.

Moshkov, Nikita, Michael Bornholdt, Santiago Benoit, Matthew Smith, Claire McQuin, Allen Goodman, Rebecca A. Senft, Yu Han, Mehrtash Babadi, Peter Horvath, et al. (Feb. 2024). “Learning representations for image-based profiling of perturbations”. Nature Communications, 15: 1, p. 1594.

Mueller, Sandra, Marcus Schittenhelm, Friedemann Honecker, Elke Malenke, Kirsten Lauber, Sebastian Wesselborg, Joerg T. Hartmann, Carsten Bokemeyer, and Frank Mayer (Aug. 2006). “Cell-cycle progression and response of germ cell tumors to cisplatin in vitro”. International Journal of Oncology, 29: 2, pp. 471–479.

Müller, I, A Jenner, G Bruchelt, D Niethammer, and B Halliwell (1997). “Effect of concentration on the cytotoxic mechanism of doxorubicin–apoptosis and oxidative DNA damage”. 230: 2, pp. 254–257. DOI: 10.1006/bbrc.1996.5898.

Nakatsu, S, S Kondo, Y Kondo, D Yin, J W Peterson, R Kaakaji, T Morimura, H Kikuchi, J Takeuchi, and G H Barnett (1997). “Induction of apoptosis in multi-drug resistant (MDR) human glioblastoma cells by SN-38, a metabolite of the camptothecin derivative CPT-11”. 39: 5, pp. 417–423. DOI: 10.1007/s002800050592.

Nguyen, Jack T and James A Wells (2003). “Direct activation of the apoptosis machinery as a mechanism to target cancer cells”. 100: 13, pp. 7533–7538. DOI: 10.1073/pnas.1031631100.

Nguyen, T V, A Jayaraman, A Quaglino, and C J Pike (2010). “Androgens selectively protect against apoptosis in hippocampal neurones”. 22: 9, pp. 1013–1022. DOI: 10.1111/j.1365-2826.2010.02044.x.

Ocadlikova, Darina, Clara Iannarone, Anna Rita Redavid, Michele Cavo, and Antonio Curti (2020). “A Screening of Antineoplastic Drugs for Acute Myeloid Leukemia Reveals Contrasting Immunogenic Effects of Etoposide and Fludarabine”. 21: 18, p. 6802. DOI: 10.3390/ijms21186802.

Olofsson, Karl, Valentina Carannante, Madoka Takai, Björn Önfelt, and Martin Wiklund (Aug. 2021). “Single cell organization and cell cycle characterization of DNA stained multicellular tumor spheroids”. Scientific Reports, 11: 1, p. 17076.

Olubodun-Obadun, Taiwo G, Ismail O Ishola, and Olufunmilayo O Adeyemi (2021). “Potentials of autophagy enhancing natural products in the treatment of Parkinson disease”. 37: 2, pp. 99–110. DOI: 10.1515/dmpt-2021-0128.

Oplustil O’Connor, Lenka, Stuart L. Rulten, Aaron N. Cranston, Rajesh Odedra, Henry Brown, Janneke E. Jaspers, Louise Jones, Charlotte Knights, Bastiaan Evers, Attilla Ting, et al. (Oct. 2016). “The PARP Inhibitor AZD2461 Provides Insights into the Role of PARP3 Inhibition for Both Synthetic Lethality and Tolerability with Chemotherapy in Preclinical Models”. Cancer Research, 76: 20, pp. 6084–6094.

Pahl, Axel, Beate Schölermann, Philipp Lampe, Marion Rusch, Mark Dow, Christian Hedberg, Adam Nelson, Sonja Sievers, Herbert Waldmann, and Slava Ziegler (June 2023). “Morphological subprofile analysis for bioactivity annotation of small molecules”. Cell Chemical Biology.

Park, I C, M J Park, C S Hwang, C H Rhee, D Y Whang, J J Jang, T B Choe, S I Hong, and S H Lee (2000). “Mitomycin C induces apoptosis in a caspases-dependent and Fas/CD95-independent manner in human gastric adenocarcinoma cells”. 158: 2, pp. 125–132. DOI: 10.1016/s0304-3835(00)00489-4.

Park, Jung Hee, Kyung Hee Jung, Soo Jung Kim, Young-Chan Yoon, Hong Hua Yan, Zhenghuan Fang, Ji Eun Lee, Joo Han Lim, Shinmee Mah, Sungwoo Hong, et al. (2019). “HS-173 as a novel inducer of RIP3-dependent necroptosis in lung cancer”. 444: pp. 94–104. DOI: 10.1016/j.canlet.2018.12.006.

Peidli, Stefan, Tessa D. Green, Ciyue Shen, Torsten Gross, Joseph Min, Samuele Garda, Bo Yuan, Linus J. Schumacher, Jake P. Taylor-King, Debora S. Marks, et al. (Jan. 2024). “scPerturb: harmonized single-cell perturbation data”. Nature Methods, pp. 1–10.

Petrazzuolo, Adriana, Maria Perez-Lanzon, Isabelle Martins, Peng Liu, Oliver Kepp, Véronique Minard-Colin, Maria Chiara Maiuri, and Guido Kroemer (2021). “Pharmacological inhibitors of anaplastic lymphoma kinase (ALK) induce immunogenic cell death through on-target effects”. 12: 8, pp. 713–713. DOI: 10.1038/s41419-021-03997-x.

Pfaendler, Ramon, Jacob Hanimann, Sohyon Lee, and Berend Snijder (Jan. 2023). Self-supervised vision transformers accurately decode cellular state heterogeneity. bioRxiv, 2023. DOI: 10.1101/2023.01.16.524226.

Pirnia, F, E Schneider, D C Betticher, and M M Borner (2002). “Mitomycin C induces apoptosis and caspase-8 and -9 processing through a caspase-3 and Fas-independent pathway”. 9: 9, pp. 905–914. DOI: 10.1038/sj.cdd.4401062.

Ranjan, Alok, Itishree Kaushik, and Sanjay K Srivastava (2020). “Pimozide Suppresses the Growth of Brain Tumors by Targeting STAT3-Mediated Autophagy”. 9: 9, p. 2141. DOI: 10.3390/cells9092141.

Rébé, Cédric, Lucie Demontoux, Thomas Pilot, and François Ghiringhelli (2019). “Platinum Derivatives Effects on Anticancer Immune Response”. 10: 1, p. 13. DOI: 10.3390/biom10010013.

Rietdijk, Jonne, Marianna Tampere, Aleksandra Pettke, Polina Georgiev, Maris Lapins, Ulrika Warpman-Berglund, Ola Spjuth, Marjo-Riitta Puumalainen, and Jordi Carreras-Puigvert (Dec. 2021). “A phenomics approach for antiviral drug discovery”. BMC Biology, 19: 1, p. 156.

Roccaro, Aldo M, Antonio Sacco, Xiaojing Jia, Ranjit Banwait, Patricia Maiso, Feda Azab, Ludmila Flores, Salomon Manier, Abdel Kareem Azab, and Irene M Ghobrial (2012). “Mechanisms of activity of the TORC1 inhibitor everolimus in Waldenstrom macroglobulinemia”. 18: 24, pp. 6609–6622. DOI: 10.1158/1078-0432.CCR-12-1532.

Romano, M F, A Lamberti, M C Turco, and S Venuta (2000). “CD40 and B chronic lymphocytic leukemia cell response to fludarabine: the influence of NF-kappaB/Rel transcription factors on chemotherapy-induced apoptosis”. 36: 3–4, pp. 255–262. DOI: 10.3109/10428190009148846.

Roukos, Vassilis, Gianluca Pegoraro, Ty C. Voss, and Tom Misteli (Feb. 2015). “Cell cycle staging of individual cells by fluorescence microscopy”. Nature Protocols, 10: 2, pp. 334–348.

Rozario, Pritisha, Miriam Pinilla, Leana Gorse, Anna Constance Vind, Kim S Robinson, Gee Ann Toh, Muhammad Jasrie Firdaus, José Francisco Martínez, Swat Kim Kerk, Zhewang Lin, et al. (2024). “Mechanistic basis for potassium efflux-driven activation of the human NLRP1 inflammasome”. 121: 2, e2309579121–e2309579121. DOI: 10.1073/pnas.2309579121.

Sanchez-Fernandez, Ana, Elisabeth Rumetshofer, Sepp Hochreiter, and Günter Klambauer (Nov. 2023). “CLOOME: contrastive learning unlocks bioimaging databases for queries with chemical structures”. Nature Communications, 14: 1, p. 7339.

Schorpp, Kenji, Alaa Bessadok, Aidin Biibosunov, Ina Rothenaigner, Stefanie Strasser, Tingying Peng, and Kamyar Hadian (July 2023). “CellDeathPred: a deep learning framework for ferroptosis and apoptosis prediction based on cell painting”. Cell Death Discovery, 9: 1, pp. 1–11.

Schwänen, C, T Hecker, G Hübinger, M Wölfle, W Rittgen, L Bergmann, and T Karakas (2002). “In vitro evaluation of bendamustine induced apoptosis in B-chronic lymphocytic leukemia”. 16: 10, pp. 2096–2105. DOI: 10.1038/sj.leu.2402651.

Selvaraju, Ramprasaath R., Michael Cogswell, Abhishek Das, Ramakrishna Vedantam, Devi Parikh, and Dhruv Batra (Dec. 2019). Grad-CAM: Visual Explanations from Deep Networks via Gradient-based Localization. arXiv, 2019. DOI: 10.48550/arXiv.1610.02391.

Shahsavari, Zahra, Fatemeh Karami-Tehrani, and Siamak Salami (2015). “Shikonin Induced Necroptosis via Reactive Oxygen Species in the T-47D Breast Cancer Cell Line”. 16: 16, pp. 7261–7266. DOI: 10.7314/apjcp.2015.16.16.7261.

Shen, Sikou, Yina Shao, and Chenghua Li (Aug. 2023). “Different types of cell death and their shift in shaping disease”. Cell Death Discovery, 9: 1, pp. 1–12.

Shimada, Kenichi, Rachid Skouta, Anna Kaplan, Wan Seok Yang, Miki Hayano, Scott J Dixon, Lewis M Brown, Carlos A Valenzuela, Adam J Wolpaw, and Brent R Stockwell (2016). “Global survey of cell death mechanisms reveals metabolic regulation of ferroptosis”. 12: 7, pp. 497–503. DOI: 10.1038/nchembio.2079.

Singh, S., M.-A. Bray, T.r. Jones, and A.e. Carpenter (2014). “Pipeline for illumination correction of images for high-throughput microscopy”. Journal of Microscopy, 256: 3, pp. 231–236.

Solimando, Antonio G and Angelo Vacca (2024). “Immunogenic therapy: new actors in myeloma”. 143: 25, pp. 2564–2565. DOI: 10.1182/blood.2024024709.

Srivastava, Shikha, Ranganatha R Somasagara, Mahesh Hegde, Mayilaadumveettil Nishana, Satish Kumar Tadi, Mrinal Srivastava, Bibha Choudhary, and Sathees C Raghavan (2016). “Quercetin, a Natural Flavonoid Interacts with DNA, Arrests Cell Cycle and Causes Tumor Regression by Activating Mitochondrial Pathway of Apoptosis”. 6: pp. 24049–24049. DOI: 10.1038/srep24049.

Stirling, David R., Madison J. Swain-Bowden, Alice M. Lucas, Anne E. Carpenter, Beth A. Cimini, and Allen Goodman (Dec. 2021). “CellProfiler 4: improvements in speed, utility and usability”. BMC Bioinformatics, 22: 1, p. 433.

Stockwell, Brent R. (July 2022). “Ferroptosis turns 10: Emerging mechanisms, physiological functions, and therapeutic applications”. Cell, 185: 14, pp. 2401–2421.

Stossi, Fabio, Pankaj K. Singh, Michela Marini, Kazem Safari, Adam T. Szafran, Alejandra Rivera Tostado, Christopher D. Candler, Maureen G. Mancini, Elina A. Mosa, Michael J. Bolt, et al. (Nov. 2024). “SPACe: an open-source, single-cell analysis of Cell Painting data”. Nature Communications, 15: 1, p. 10170.

Stringer, Carsen, Tim Wang, Michalis Michaelos, and Marius Pachitariu (Jan. 2021). “Cellpose: a generalist algorithm for cellular segmentation”. Nature Methods, 18: 1, pp. 100–106.

Tanevski, Jovan, Ricardo Omar Ramirez Flores, Attila Gabor, Denis Schapiro, and Julio Saez-Rodriguez (Apr. 2022). “Explainable multiview framework for dissecting spatial relationships from highly multiplexed data”. Genome Biology, 23: 1, p. 97.

Teng, Jin-Feng, Qi-Bing Mei, Xiao-Gang Zhou, Yong Tang, Rui Xiong, Wen-Qiao Qiu, Rong Pan, Betty Yuen-Kwan Law, Vincent Kam-Wai Wong, Chong-Lin Yu, et al. (2020). “Polyphyllin VI Induces Caspase-1-Mediated Pyroptosis via the Induction of ROS/NF-B/NLRP3/GSDMD Signal Axis in Non-Small Cell Lung Cancer”. 12: 1, p. 193. DOI: 10.3390/cancers12010193.

Tesniere, A, F Schlemmer, V Boige, O Kepp, I Martins, F Ghiringhelli, L Aymeric, M Michaud, L Apetoh, L Barault, et al. (2010). “Immunogenic death of colon cancer cells treated with oxaliplatin”. 29: 4, pp. 482–491. DOI: 10.1038/onc.2009.356.

Tesniere, A, F Schlemmer, V Boige, O Kepp, I Martins, F Ghiringhelli, L Aymeric, M Michaud, L Apetoh, L Barault, (Jan. 2010). “Immunogenic death of colon cancer cells treated with oxaliplatin”. Oncogene, 29: 4, pp. 482–491.

Tian, Guangyan, Philip J Harrison, Akshai P Sreenivasan, Jordi Carreras-Puigvert, and Ola Spjuth (Dec. 2023). “Combining molecular and cell painting image data for mechanism of action prediction”. Artificial Intelligence in the Life Sciences, 3: p. 100060.

Topçul, Mehmet R and İdil Çetin (2022). “Antitumor Activity of Tubulin-Binding Agent MPC-6827 on Different Types of Cancer Cell Lines”. 68: 4, pp. 108–112. DOI: 10.14715/cmb/2022.68.4.13.

Traganos, F, K Seiter, E Feldman, H D Halicka, and Z Darzynkiewicz (1996). “Induction of apoptosis by camptothecin and topotecan”. 803: pp. 101–110. DOI: 10.1111/j.1749-6632.1996.tb26380.x.

Ueno, Munehisa, Shoichi Nonaka, Ryuta Yamazaki, Nobuhiro Deguchi, and Masaru Murai (2002). “SN-38 induces cell cycle arrest and apoptosis in human testicular cancer”. 42: 4, pp. 390–397. DOI: 10.1016/s0302-2838(02)00321-4.

Wang, Jin, Jia-Hui Yang, Di Xiong, and Ling Chen (2025). “Activation of SIRT3/AMPK/mTOR-mediated autophagy promotes quercetin-induced ferroptosis in oral squamous cell carcinoma”. 44: pp. 9603271251323753–9603271251323753. DOI: 10.1177/09603271251323753.

Wang, Jingmiao, Haizhong Zhang, Xiaoyan Yin, and Yanrui Bian (2022). “Oxaliplatin Induces Immunogenic Cell Death in Human and Murine Laryngeal Cancer”. 2022: pp. 3760766–3760766. DOI: 10.1155/2022/3760766.

Wang, Lujue, Yuan Li, Tongxin Niu, Jing Deng, Yuxian Shi, Yating Liu, Boding Tong, Xin Qi, Dan Cao, Yongguang Tao, et al. (2025). “Simvastatin-Induced Ferroptosis in Orbital Fibroblasts in Graves’ Ophthalmopathy”. 66: 1, pp. 56–56. DOI: 10.1167/iovs.66.1.56.

Wang, Meichen, Leilei Liang, Rong Wang, Shutao Jia, Chang Xu, Yuting Wang, Min Luo, Qiqi Lin, Min Yang, Hongyu Zhou, et al. (2023). “Narciclasine, a novel topoisomerase I inhibitor, exhibited potent anti-cancer activity against cancer cells”. 13: 1, pp. 27–27. DOI: 10.1007/s13659-023-00392-1.

Wang, Suwei, Eugene A Konorev, Srigiridhar Kotamraju, Joy Joseph, Shasi Kalivendi, and B Kalyanaraman (2004). “Doxorubicin induces apoptosis in normal and tumor cells via distinctly different mechanisms. intermediacy of H(2)O(2)- and p53-dependent pathways”. 279: 24, pp. 25535–25543. DOI: 10.1074/jbc.M400944200.

Wang, Xiang, Xinxin Ren, Xu Lin, Qi Li, Yingqiong Zhang, Jun Deng, Binxin Chen, Guoqing Ru, Ying Luo, and Nengming Lin (2024). “Recent progress of ferroptosis in cancers and drug discovery”. 19: 4, pp. 100939–100939. DOI: 10.1016/j.ajps.2024.100939.

Wang, Xinyu, Liwen Fan, Xuanzhong Wang, Tianfei Luo, and Linlin Liu (2022). “Cyclophilin A contributes to shikonin-induced glioma cell necroptosis and promotion of chromatinolysis”. 12: 1, pp. 14675–14675. DOI: 10.1038/s41598-022-19066-y.

Wang, Zining, Jiemin Chen, Jie Hu, Hongxia Zhang, Feifei Xu, Wenzhuo He, Xiaojuan Wang, Mengyun Li, Wenhua Lu, Gucheng Zeng, et al. (2019). “cGAS/STING axis mediates a topoisomerase II inhibitor-induced tumor immunogenicity”. 129: 11, pp. 4850–4862. DOI: 10.1172/JCI127471.

Way et al., Gregor (2022). Pycytominer: Data processing functions for profiling perturbations. 2022.

Willis, Clinton, Johanna Nyffeler, and Joshua Harrill (Aug. 2020). “Phenotypic Profiling of Reference Chemicals across Biologically Diverse Cell Types Using the Cell Painting Assay”. SLAS DISCOVERY: Advancing the Science of Drug Discovery, 25: 7, pp. 755–769.

Wolf, F. Alexander, Philipp Angerer, and Fabian J. Theis (Feb. 2018). “SCANPY: large-scale single-cell gene expression data analysis”. Genome Biology, 19: 1, p. 15.

Wu, Jianjian, Qiang Guo, Juntao Li, Hao Yuan, Chutian Xiao, Jianguang Qiu, Qiong Wu, and Dejuan Wang (2023). “Loperamide induces protective autophagy and apoptosis through the ROS/JNK signaling pathway in bladder cancer”. 218: pp. 115870–115870. DOI: 10.1016/j.bcp.2023.115870.

Wu, Shuhong, Hongbo Zhu, Jian Gu, Lidong Zhang, Fuminori Teraishi, John J Davis, Dietmar A Jacob, and Bingliang Fang (2004). “Induction of apoptosis and down-regulation of Bcl-XL in cancer cells by a novel small molecule, 2[[3-(2,3-dichlorophenoxy)propyl]amino]ethanol”. 64: 3, pp. 1110–1113. DOI: 10.1158/0008-5472.can-03-2790.

Wu, Yueshu, Jun Yang, Gang Xu, Xiaolin Chen, and Xiaochen Qu (Apr. 2024). “Integrated analysis of single-cell and bulk RNA sequencing data reveals prognostic characteristics of lysosome-dependent cell death-related genes in osteosarcoma”. BMC Genomics, 25: 1, p. 379.

Xia, Yu, Pu Huang, Yi-Yu Qian, Zanhong Wang, Ning Jin, Xin Li, Wen Pan, Si-Yuan Wang, Ping Jin, Emmanuel Kwateng Drokow, et al. (2024). “PARP inhibitors enhance antitumor immune responses by triggering pyroptosis via TNF-caspase 8-GSDMD/E axis in ovarian cancer”. 12: 10, e009032. DOI: 10.1136/jitc-2024-009032.

Xie, Y, W Hou, X Song, Y Yu, J Huang, X Sun, R Kang, and D Tang (2016). “Ferroptosis: process and function”. 23: 3, pp. 369–379. DOI: 10.1038/cdd.2015.158.

Xin, Meiguo, Rui Li, Maohua Xie, Dongkyoo Park, Taofeek K Owonikoko, Gabriel L Sica, Patrick E Corsino, Jia Zhou, Chunyong Ding, Mark A White, et al. (2014). “Small-molecule Bax agonists for cancer therapy”. 5: pp. 4935–4935. DOI: 10.1038/ncomms5935.

Yao, Wenchao, Xuxu Liu, Yuanhang He, Maolan Tian, Shixin Lu, Qiang Wang, Yi Zheng, Zhenyi Lv, Chenjun Hao, Dongbo Xue, et al. (Dec. 2022). “ScRNA-seq and bulk RNA-seq reveal the characteristics of ferroptosis and establish a risk signature in cholangiocarcinoma”. Molecular Therapy - Oncolytics, 27: pp. 48–60.

Yu, Pian, Xu Zhang, Nian Liu, Ling Tang, Cong Peng, and Xiang Chen (Mar. 2021a). “Pyroptosis: mechanisms and diseases”. Signal Transduction and Targeted Therapy, 6: 1, pp. 1–21.

Yu, Pian, Xu Zhang, Nian Liu, Ling Tang, Cong Peng, and Xiang Chen (2021b). “Pyroptosis: mechanisms and diseases”. 6: 1, pp. 128–128. DOI: 10.1038/s41392-021-00507-5.

Zeng, Cheng-Wu, Xing-Ju Zhang, Kang-Yu Lin, Hua Ye, Shu-Ying Feng, Hua Zhang, and Yue-Qin Chen (2012). “Camptothecin induces apoptosis in cancer cells via microRNA-125b-mediated mitochondrial pathways”. 81: 4, pp. 578–586. DOI: 10.1124/mol.111.076794.

Zhang, Z W, S E Patchett, and M J Farthing (2000). “Topoisomerase I inhibitor (camptothecin)-induced apoptosis in human gastric cancer cells and the role of wild-type p53 in the enhancement of its cytotoxicity”. 11: 9, pp. 757–764. DOI: 10.1097/00001813-200010000-00013.

Zhou, Dan, Qiuhua Wu, Huajuan Qiu, Mi Li, and Yanqin Ji (2022). “Simvastatin Inhibits Endometrial Cancer Malignant Behaviors by Suppressing RAS/Mitogen-Activated Protein Kinase (MAPK) Pathway-Mediated Reactive Oxygen Species (ROS) and Ferroptosis”. 2022: pp. 6177477–6177477. DOI: 10.1155/2022/6177477.

Zhou, Qian, Yu Meng, Daishi Li, Lei Yao, Jiayuan Le, Yihuang Liu, Yuming Sun, Furong Zeng, Xiang Chen, and Guangtong Deng (Mar. 2024). “Ferroptosis in cancer: from molecular mechanisms to therapeutic strategies”. Signal Transduction and Targeted Therapy, 9: 1, pp. 1–30.

Zhu, Hanzhang, Yuqiang Shan, Ke Ge, Jun Lu, Wencheng Kong, and Changku Jia (2020). “Oxaliplatin induces immunogenic cell death in hepatocellular carcinoma cells and synergizes with immune checkpoint blockade therapy”. 43: 6, pp. 1203–1214. DOI: 10.1007/s13402-020-00552-2.

Zielke, Svenja, Nina Meyer, Muriel Mari, Khalil Abou-El-Ardat, Fulvio Reggiori, Sjoerd J L van Wijk, Donat Kögel, and Simone Fulda (2018). “Loperamide, pimozide, and STF-62247 trigger autophagy-dependent cell death in glioblastoma cells”. 9: 10, pp. 994–994. DOI: 10.1038/s41419-018-1003-1.

Zinzani, P L, M Buzzi, P Farabegoli, G Martinelli, P Tosi, E Zuffa, G Visani, N Testoni, M Salvucci, M Bendandi, et al. (1994). “Apoptosis induction with fludarabine on freshly isolated chronic myeloid leukemia cells”. 79: 2, pp. 127–131.

Zinzani, P L, M Buzzi, P Farabegoli, P Tosi, A Fortuna, G Visani, G Martinelli, A Zaccaria, and S Tura (1994). “Induction of “in vitro” apoptosis by fludarabine in freshly isolated B-chronic lymphocytic leukemia cells”. 13: 1-2, pp. 95–97. DOI: 10.3109/10428199409051657.

