## Supplementary Material for "Single-Cell Morphological Profiling Reveals Insights into Programmed Cell Death"

### 1 Supplementary

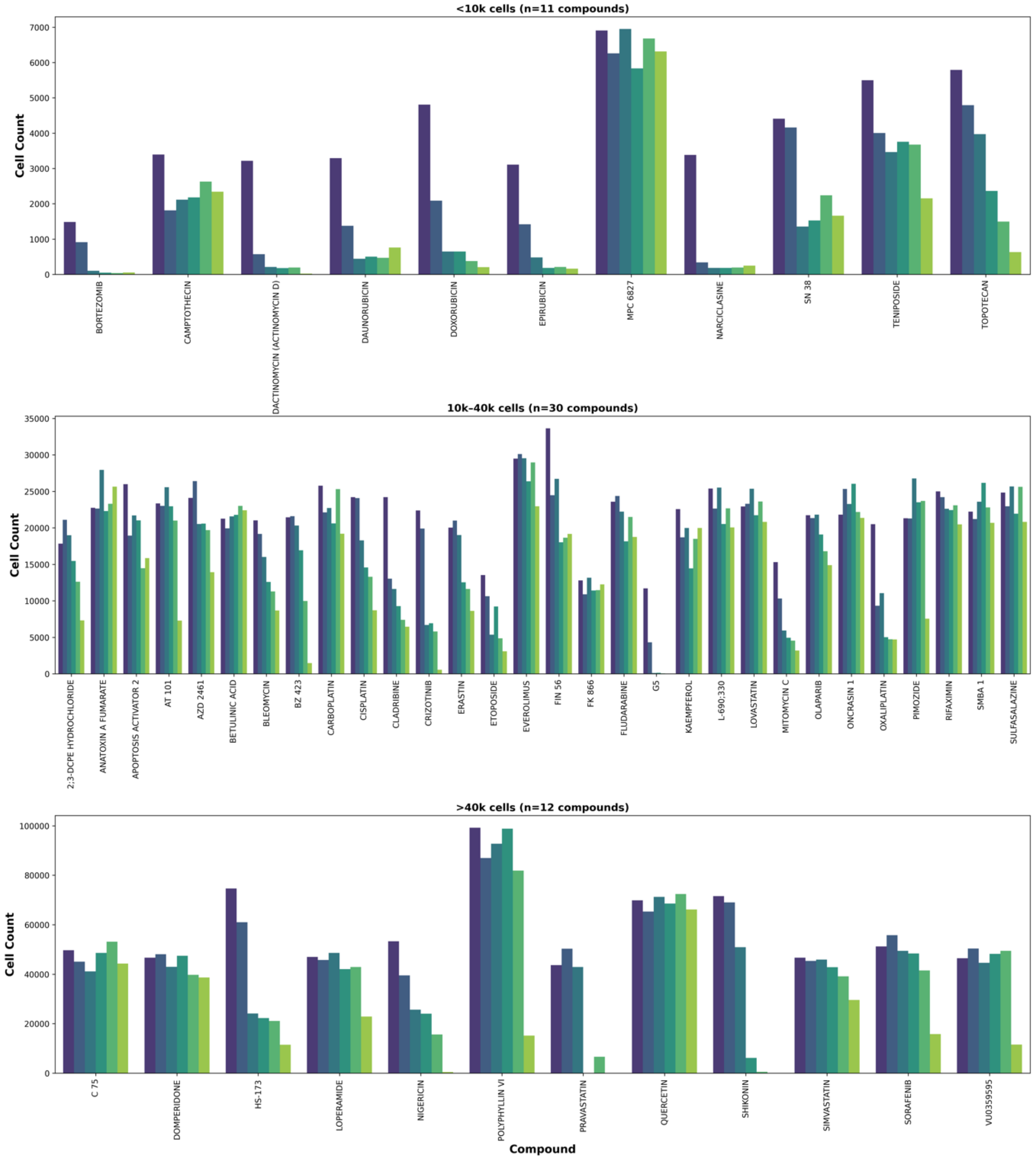


**Figure S1:** Cell counts per compound concentration of unfiltered dataset (as determined by segmented cells). Compounds are binned into compound concentrations with less than 10,000 cells, between 10,000 and 40,000 cells and more than 40,000 across all plates.

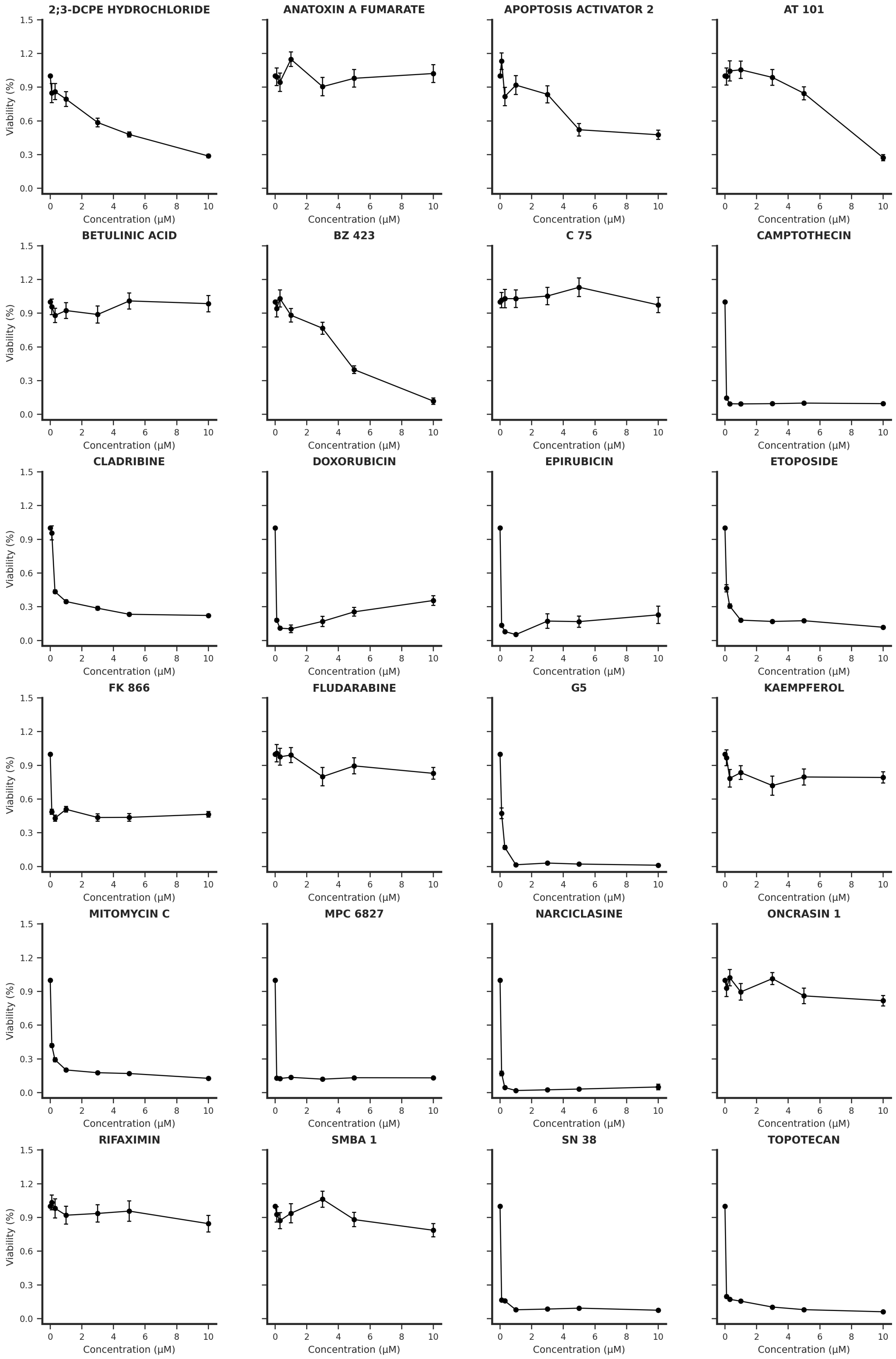


**Figure S2:** Concentration-dependent viability of compounds in apoptosis class. Viability was calculated as cell count normalized by negative control (DMSO, 1.0). Plots depict mean ± standard error (n = 4 technical replicates). X-axis depicts concentrations.

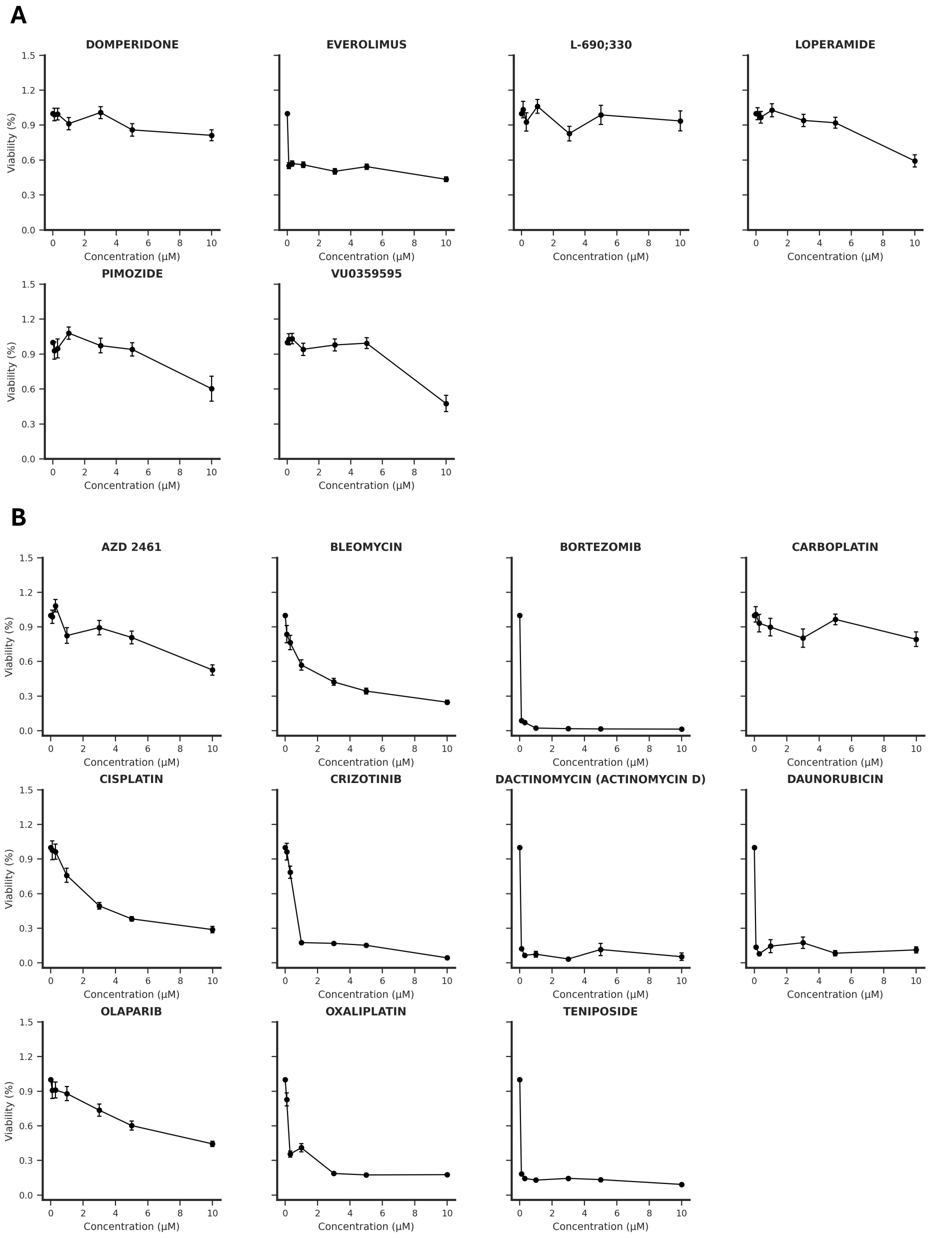


**Figure S3:** Concentration-dependent viability of compounds in autophagy and immunogenic cell death class. Viability was calculated as cell count normalized by negative control (DMSO, 1.0). Plots depict mean ± standard error (n = 4 technical replicates). X-axis depicts concentrations. (A) Autophagy compounds. (B) Immunogenic cell death compounds.

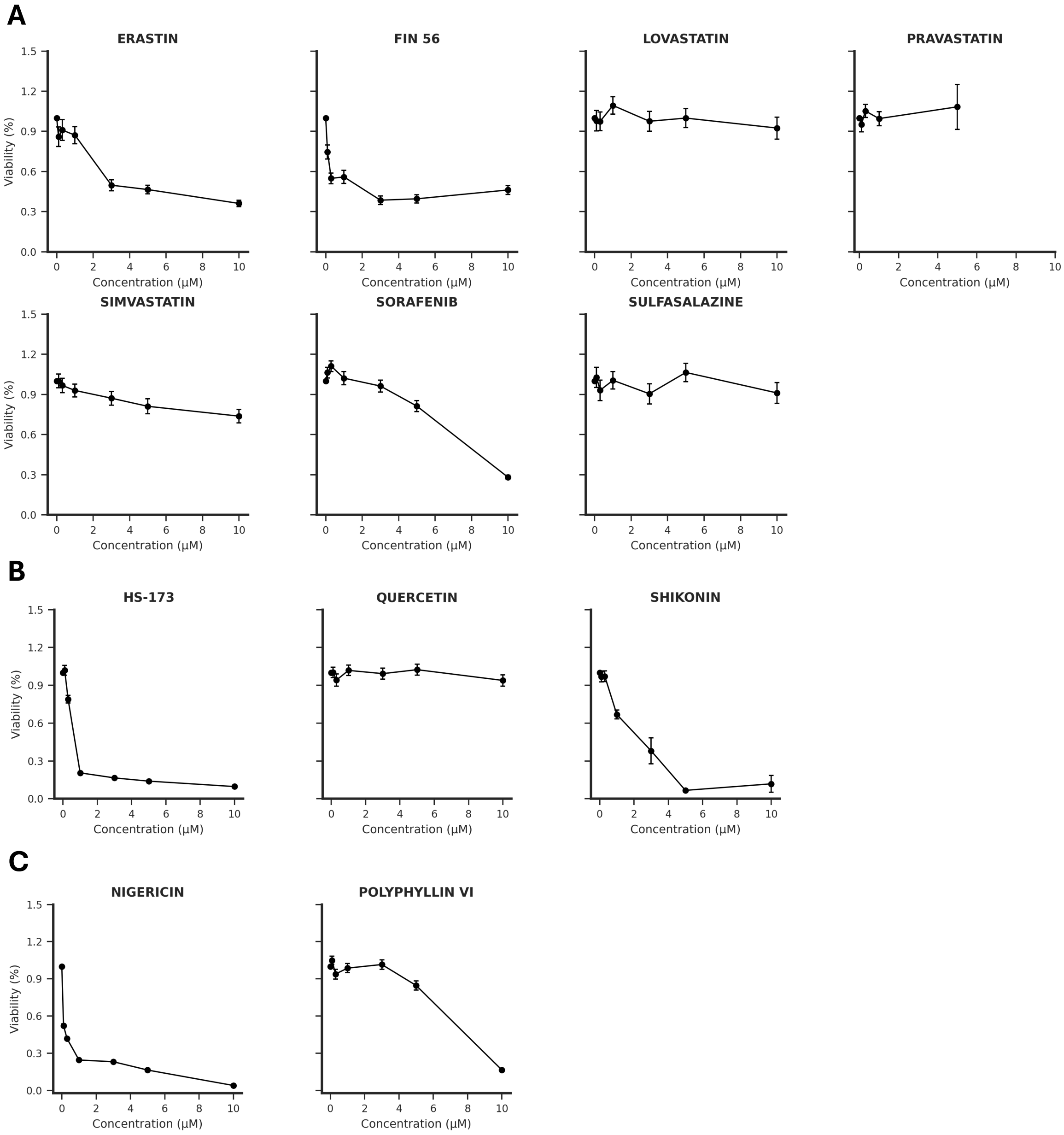


**Figure S4:** Concentration-dependent viability of compounds in ferroptosis, necroptosis and pyroptosis class. Viability was calculated as cell count normalized by negative control (DMSO, 1.0). Plots depict mean ± standard error (n = 4 technical replicates). X-axis depicts concentrations. (A) Ferroptosis compounds. (B) Necroptosis compounds (C) Pyroptosis compounds.

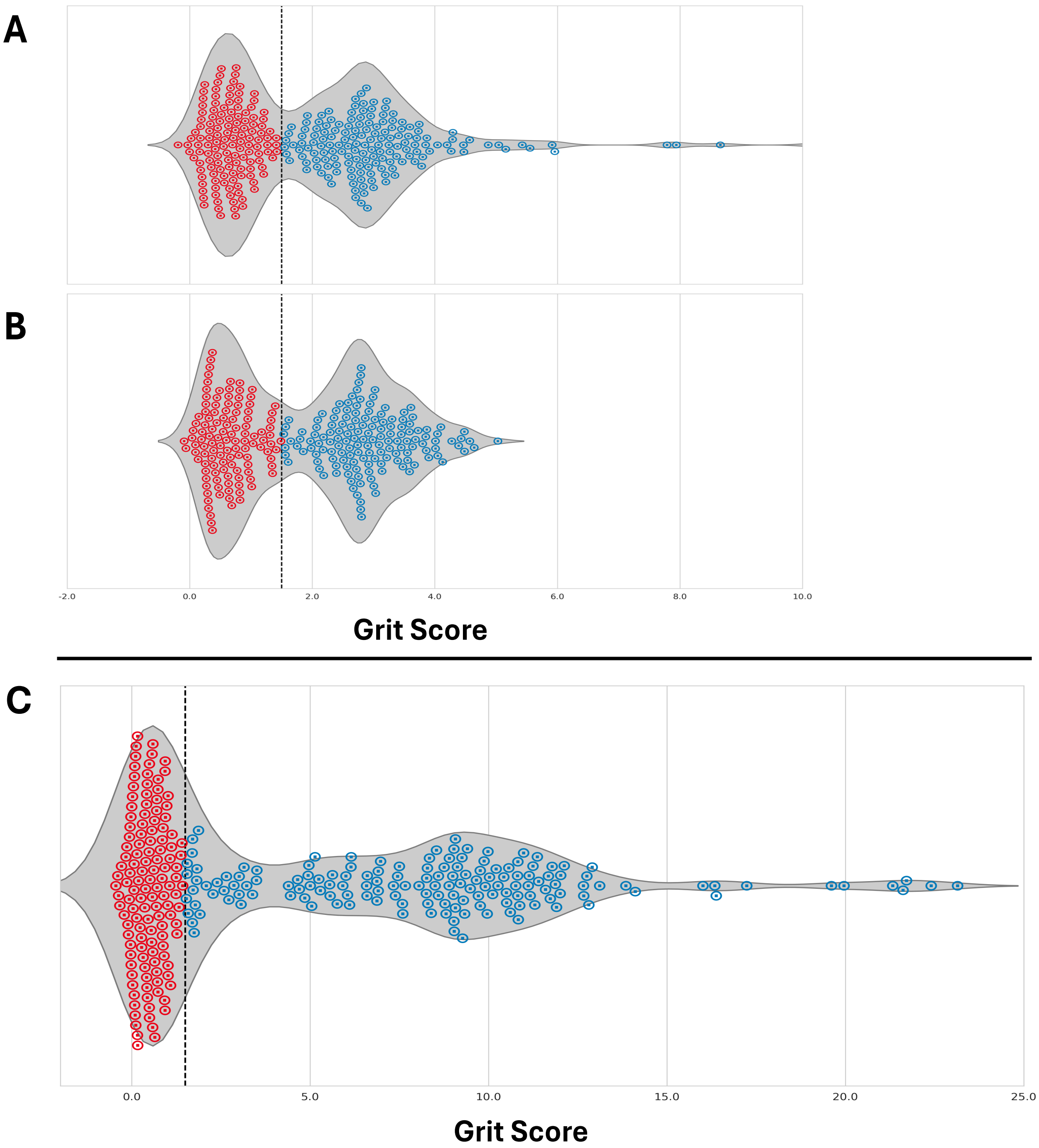


**Figure S5:** Distribution of Grit Scores with respect to DMSO. Color indicates if grit scores are below (red) or above (blue) 1.5. One point representsthe grit score per compound concentration calculated from aggregated profiles.(A) Distribution of grit scores for CellProfiler features. (B) Distribution of grit scores for DeepProfiler features. (C) Distribution of grit scores for DINO features.

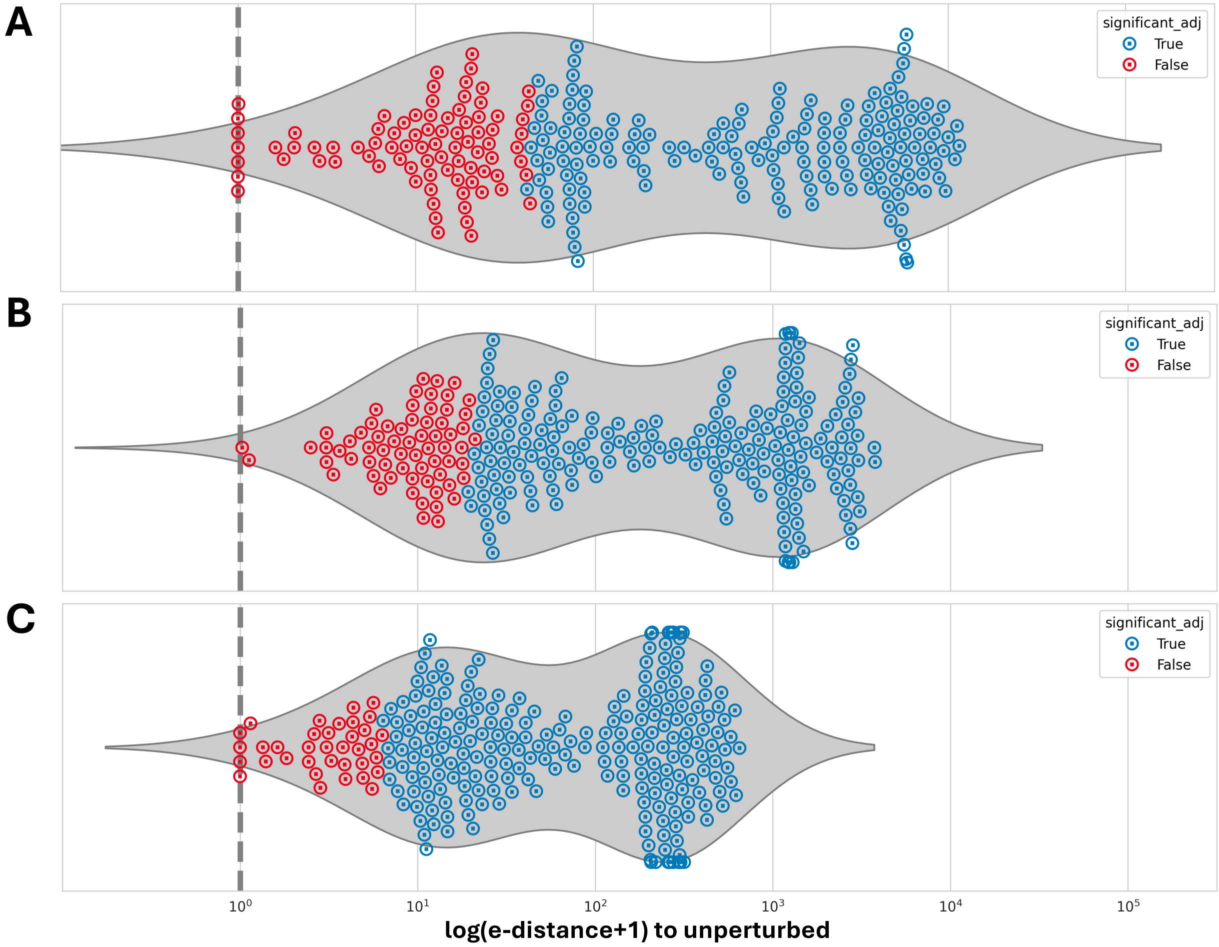


**Figure S6:** Distribution of log-transformed e-distances with respect to DMSO. Color indicates results of etest (blue significant, red not significant). One point represents e-distance calculated per compound concentration from single-cell profiles. (A) Distribution of e-distances for CellProfiler features. (B) Distribution of e-distances for DeepProfiler features. (C) Distribution of e-distances for DINO features.

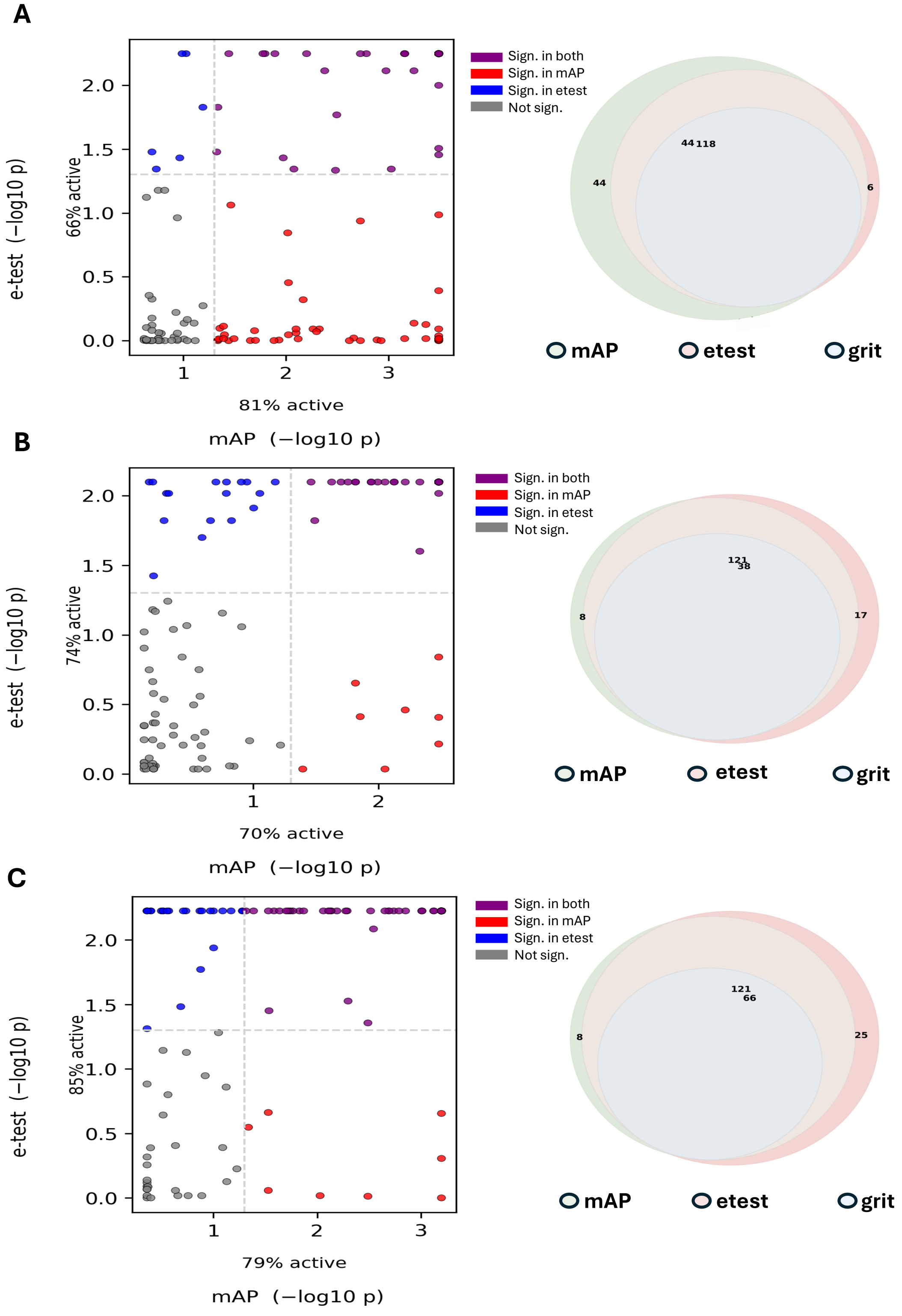


**Figure S7:** Comparison of mean average precision (mAP) with e-distance and grit scores. Each panel shows the pair-wise comparison of log-transformed p-values for mAP (x-axis) and etest (y-axis) as well as percentage of active compound concentrations on the left. The right part of the panels shows the number of significant compounds concentrations per perturbation metric (mAP, etest, grit). (A) Perturbation metrics for CellProfiler. (B) Perturbation metric for DeepProfiler. (C) Perturbation metrics for DINO.

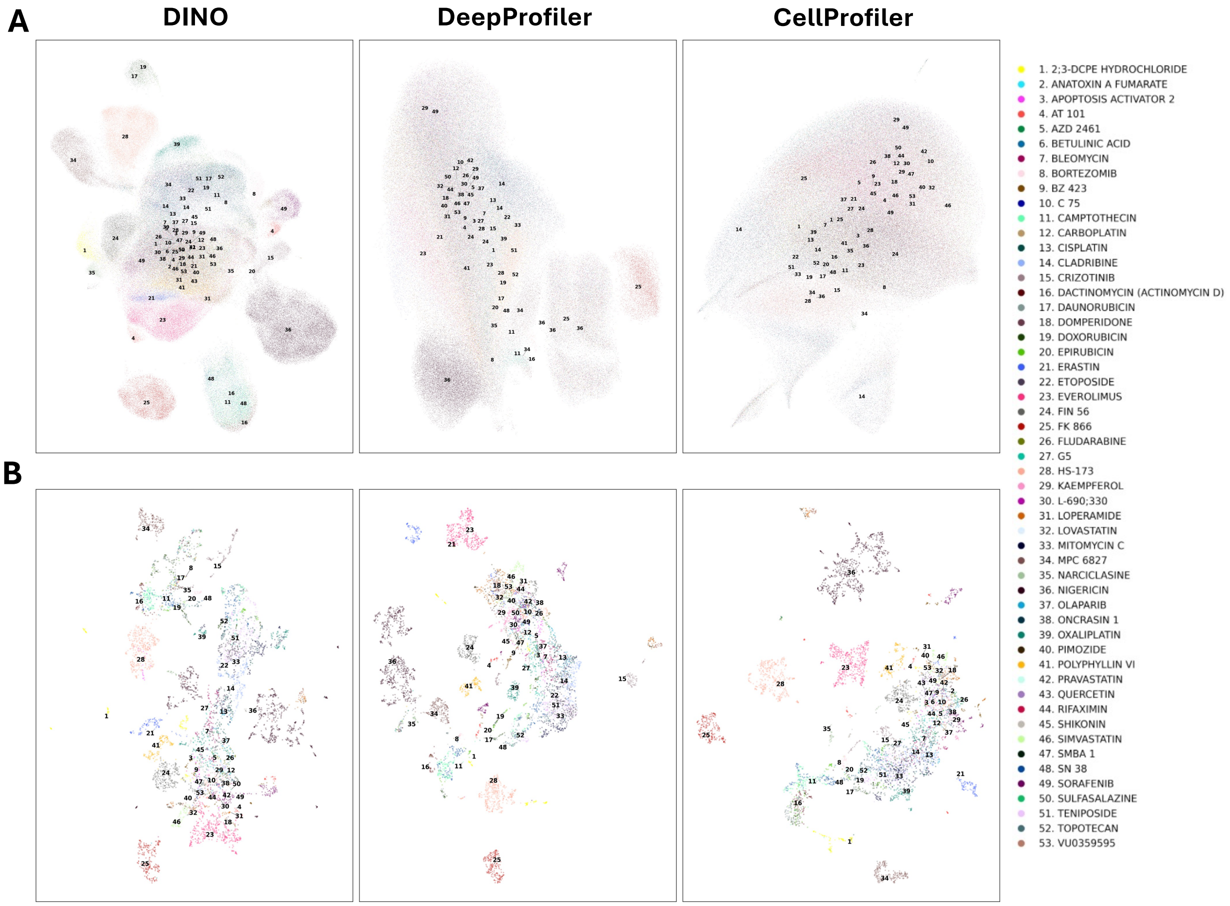


**Figure S8:** UMAP embeddings colored by compound for each feature extractor. Annotations point at compound centroids. For compounds with multiple clusters, each cluster with at least 200 cells is annotated. Left column shows DINO embeddings, middle column DeepProfiler embeddings, right column CellProfiler embeddings. (A) UMAP embedding of single-cell profiles. (B) UMAP embedding of aggregated profiles.

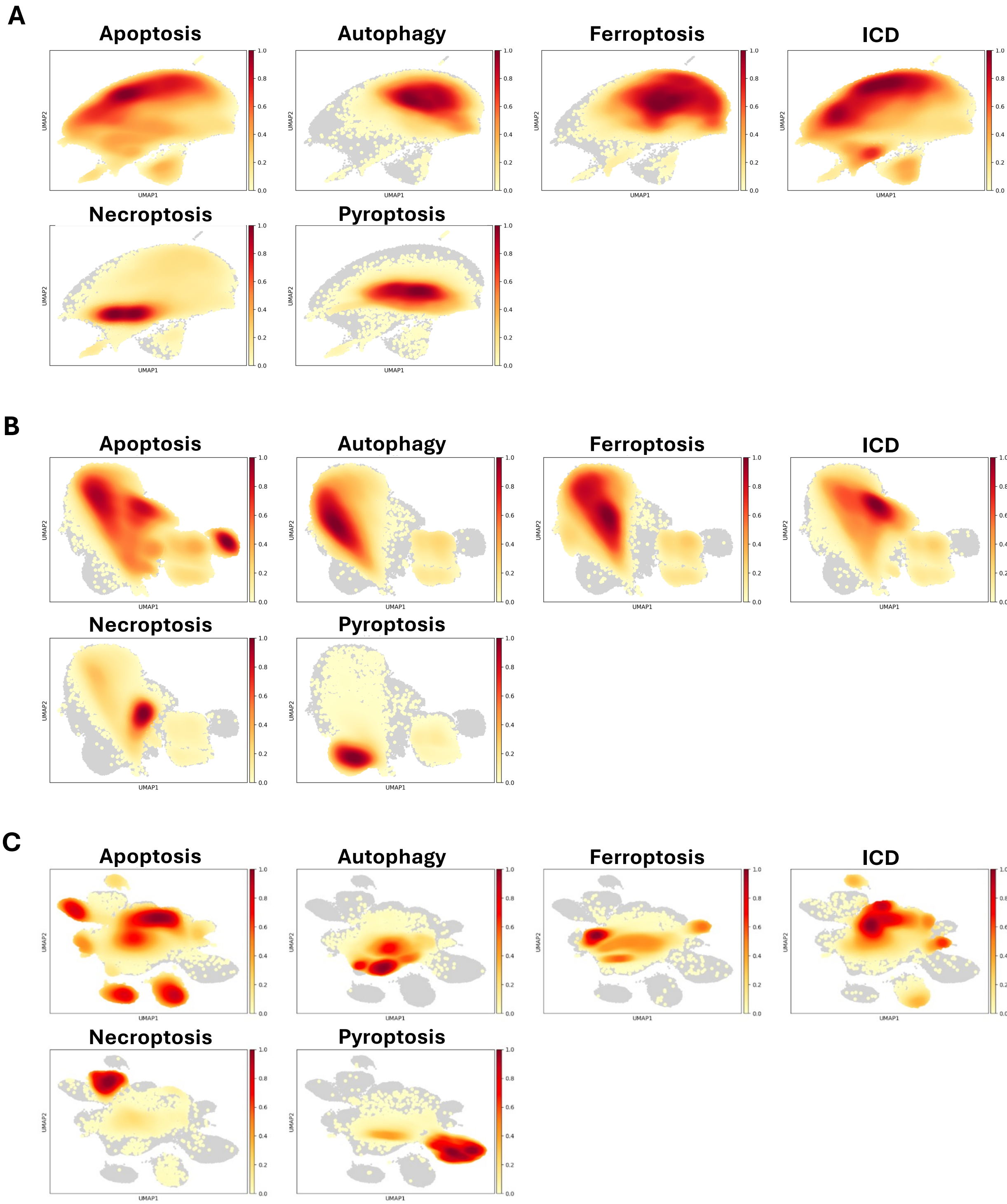


**Figure S9:** Estimated densities of cell death subtypes in UMAP space. Color indicates density regions, going from yellow (low density) to red (high density). (A) CellProfiler features. (B) DeepProfiler features. (C) DINO features.

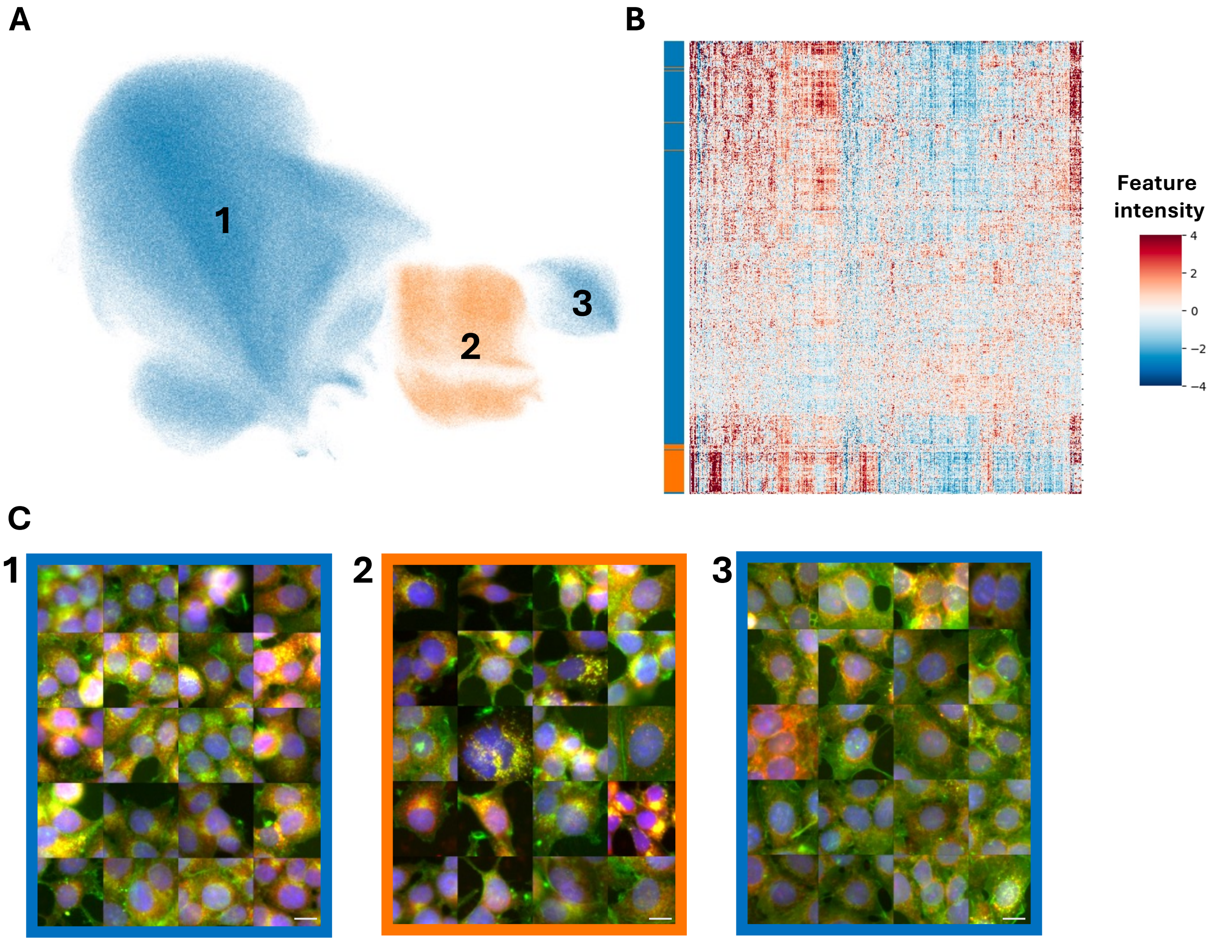


**Figure S10:** Characterization of undefined cluster in DeepProfiler features. (A) Single-cell UMAP embedding of DeepProfiler features, cluster colored in orange, remaining cells in blue, (B) Feature map of single-cell DeepProfiler features. Each column corresponds to one feature, each row to one cell. Colour indicate feature intensity. (C) Representative cells identified with *CellViewer* for each of the annotated regions in the UMAP space in A. Scale bar corresponds to 50 µm.

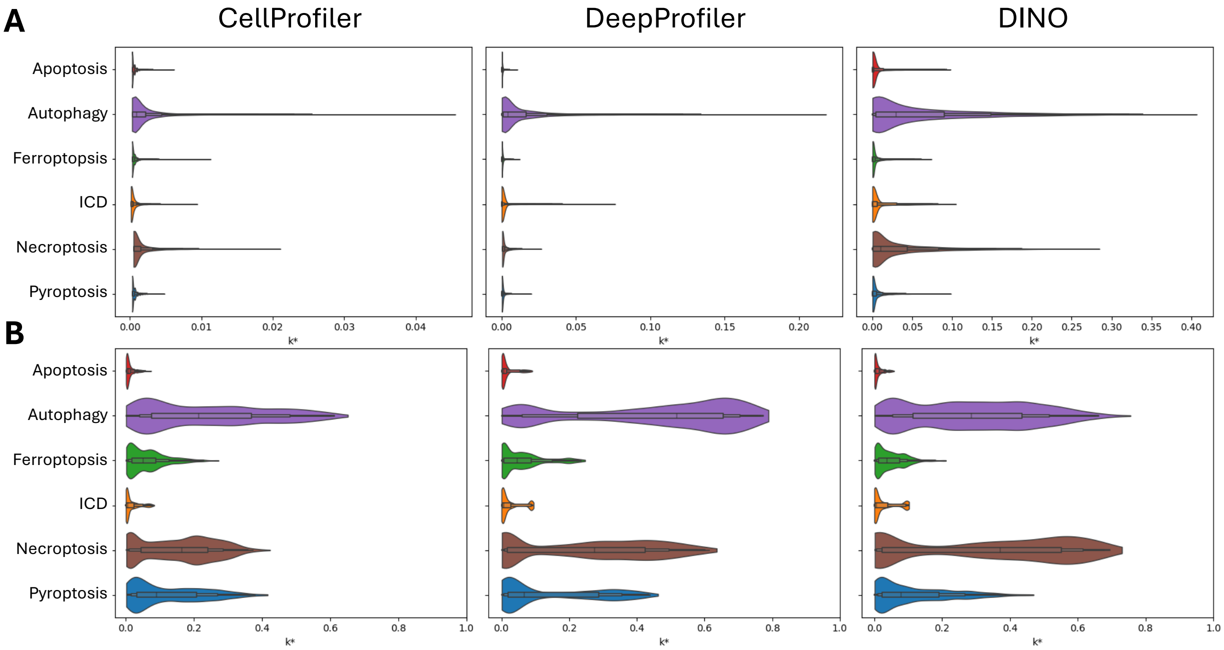


**Figure S11:** $k^*$ distributions of cell death subtypes for CellProfiler (left), DeepProfiler (middle) and DINO (right). (A) $k^*$ distributions calculated from single-cell profiles. (B) $k^*$ distributions calculated from aggregate profiles.

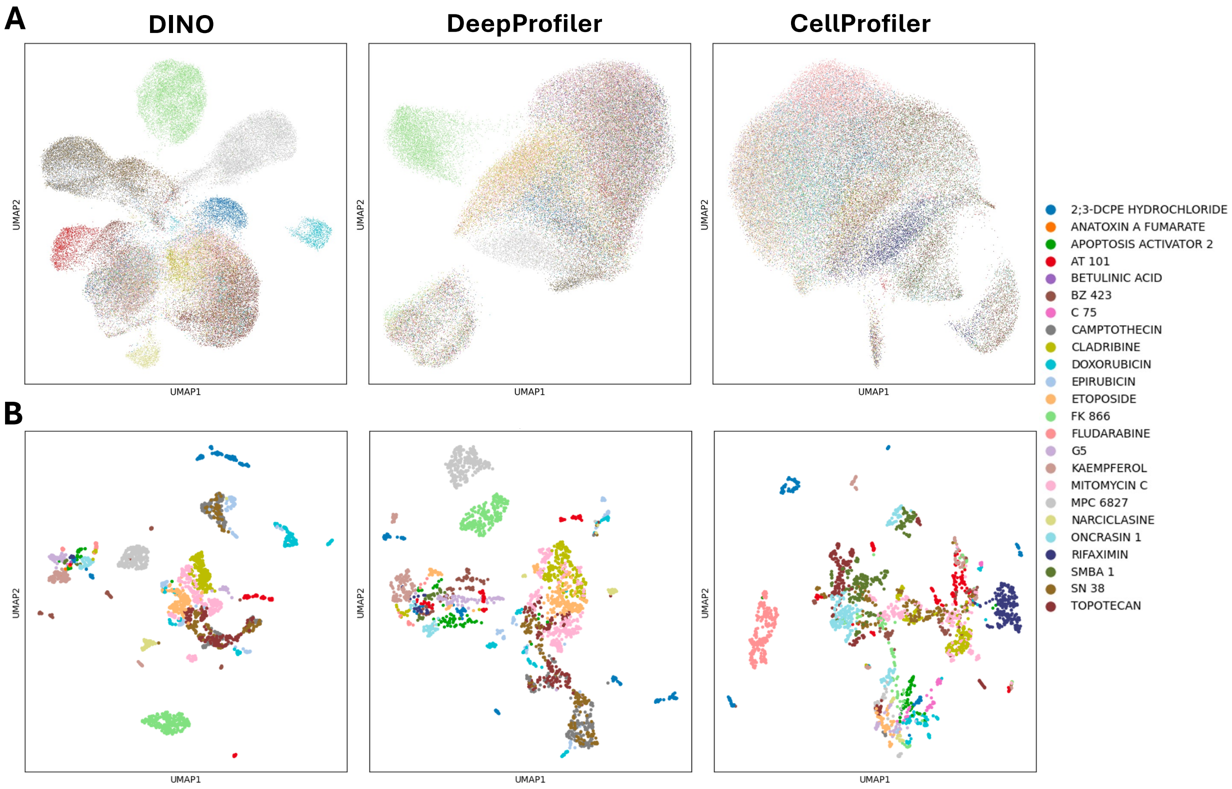


**Figure S12:** UMAP representations of apoptosis class colored by compound. (A) Single-cell UMAP embeddings for (from left to right) DINO, DeepProfiler and CellProfiler. (B) Aggregated UMAP embeddings for (from left to right) DINO, DeepProfiler and CellProfiler.

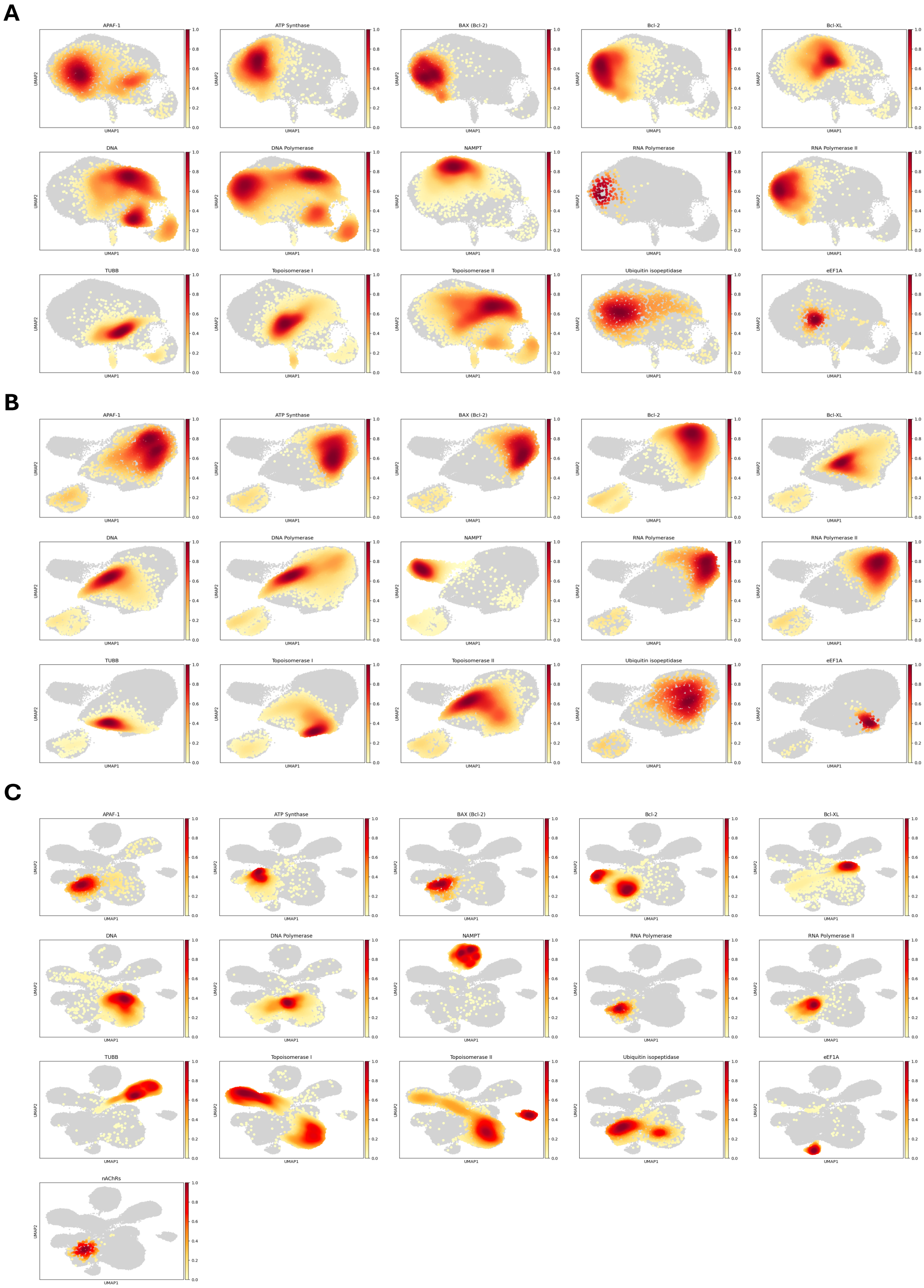


**Figure S13:** Density estimation of targets in apoptosis class. Color indicates density regions, going from yellow (low density) to red (high density). Target-compound matches can be found in Supplemental Table 1. (A) CellProfiler (B) DeepProfiler (C) DINO.

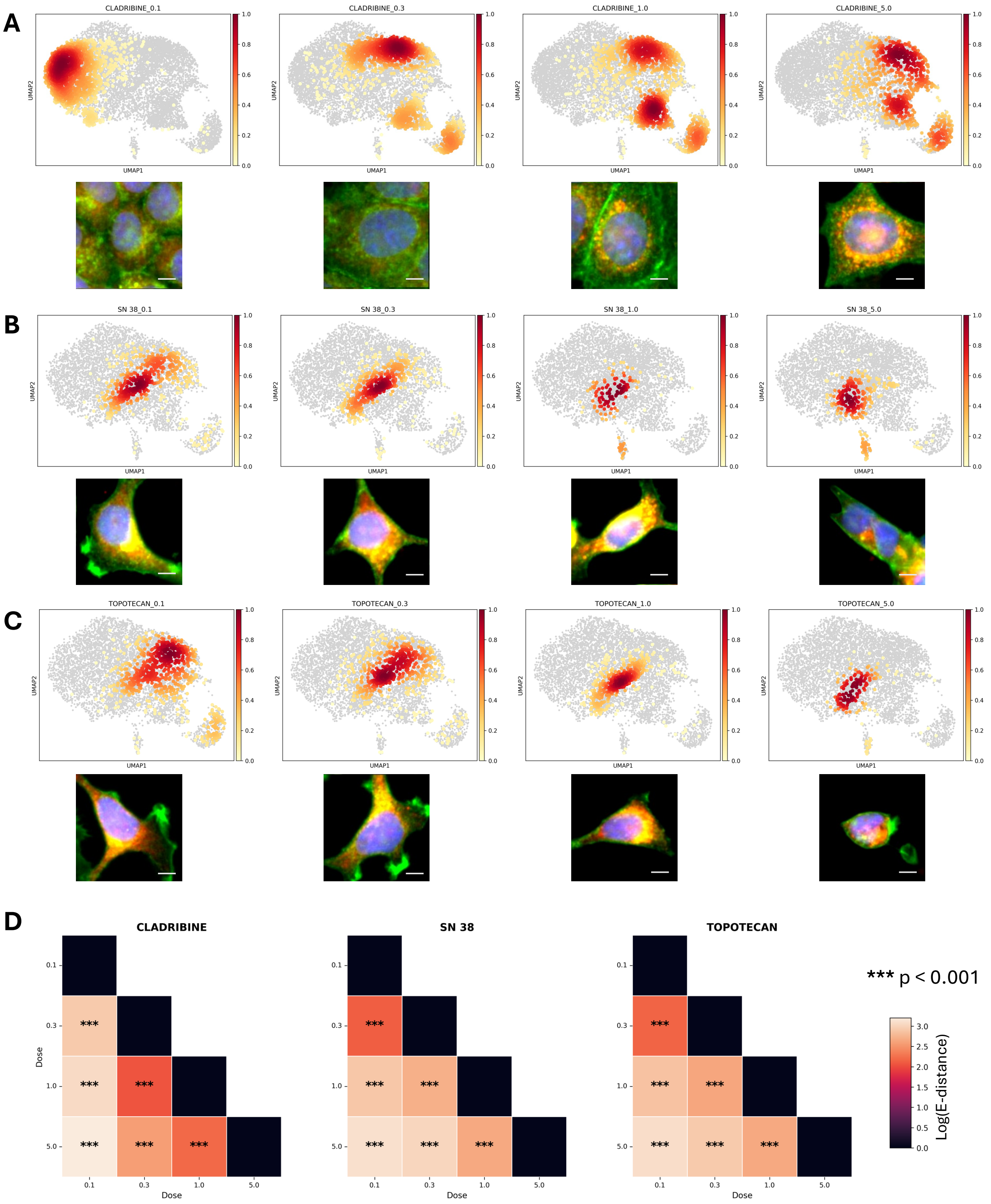


**Figure S14:** UMAP representations of selected compounds from CellProfiler with representative cells. UMAP shows density estimation for respective concentrations (0.1, 0.3, 3.0, 5.0 µM) in shared UMAP space. Bottom rows show representative cell identified via *CellViewer*. Scale bar corresponds to 10 µm. (A) Cladribine (B) SN 38 (C) Topotecan. (D) Pair-wise energy distances between concentrations (µM). Asterisk annotates significance level as calculated via etest.

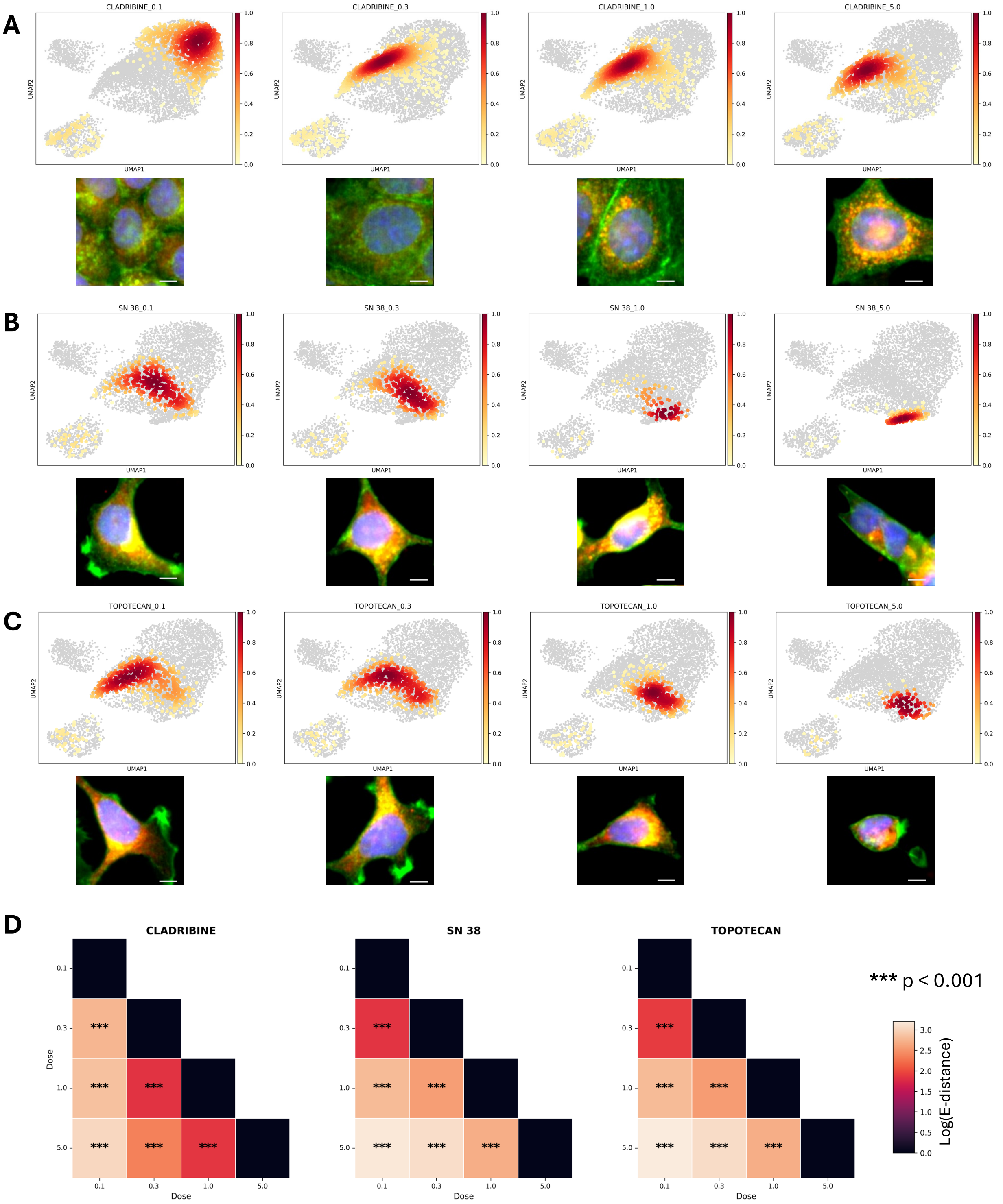


**Figure S15:** UMAP representations of selected compounds with DeepProfiler together with representative cells. UMAP shows density estimation for respective concentrations (0.1, 0.3, 3.0, 5.0 µM) in shared UMAP space. Bottom rows show representative cell identified via *CellViewer*. Scale bar corresponds to 10 µm. (A) Cladribine (B) SN 38 (C) Topotecan.(D) Pair-wise energy distances between concentrations (µM). Asterisk annotates significance level as calculated via etest.

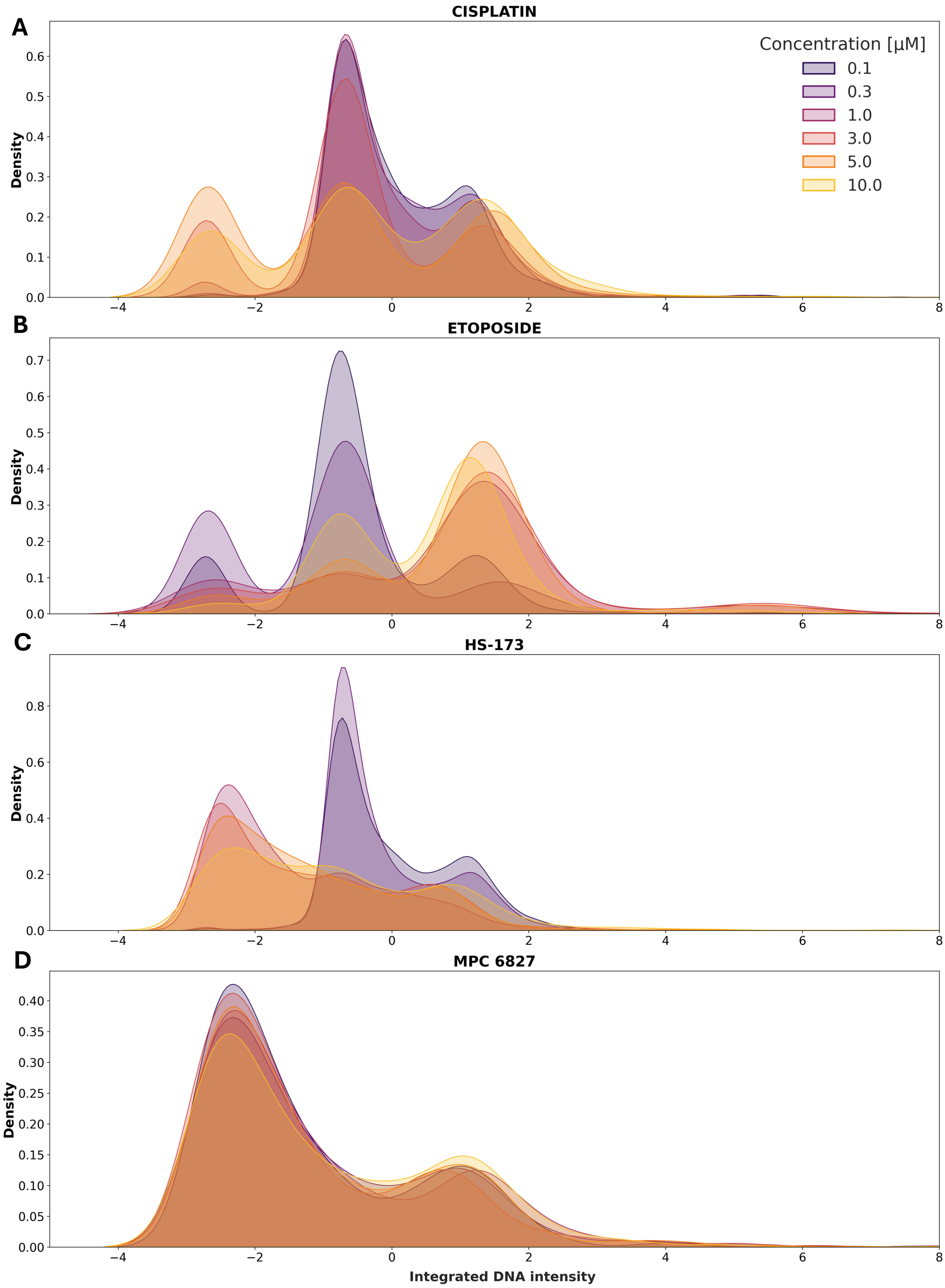


**Figure S16:** Distribution of normalized integrated DNA intensity for selected compounds. Color indicates compound concentration. (A) Cisplatin (B) Etoposide (C) HS-713 (D) MPC 6827.

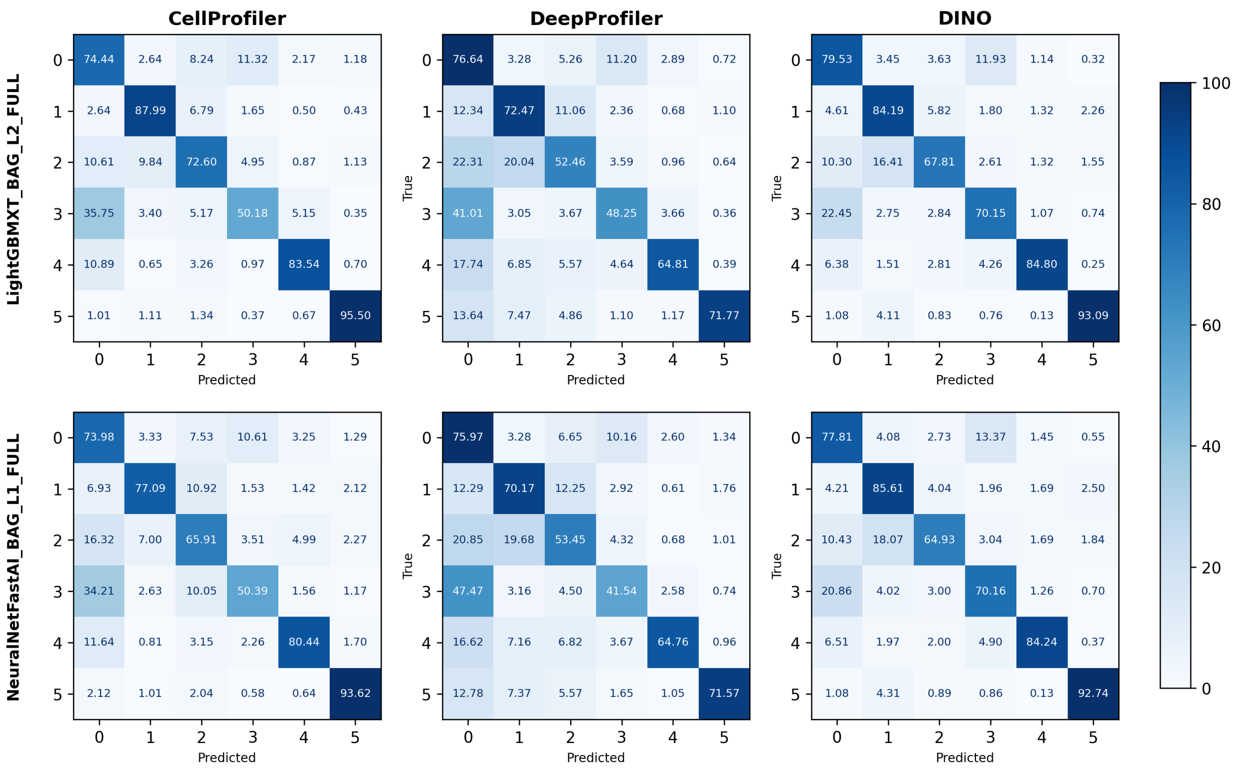


**Figure S17:** Confusion matrices of single-cell classifiers in Table 2, with label codes: 0 (apoptosis), 1 (Autophagy), 2 (Ferroptosis), 3 (Immunogenic cell death), 4 (Necroptosis), 5 (Pyroptosis). Numbers indicate percentage of correctly predicted labels in respective field. Top row: Confusion matrices for LightGBMXT_BAG_L2 models for CellProfiler, DeepProfiler and DINO (from left to right). Bottom row: Confusion matrices for NeuralNetFastAI_BAG_L1 models for CellProfiler, DeepProfiler and DINO (from left to right).

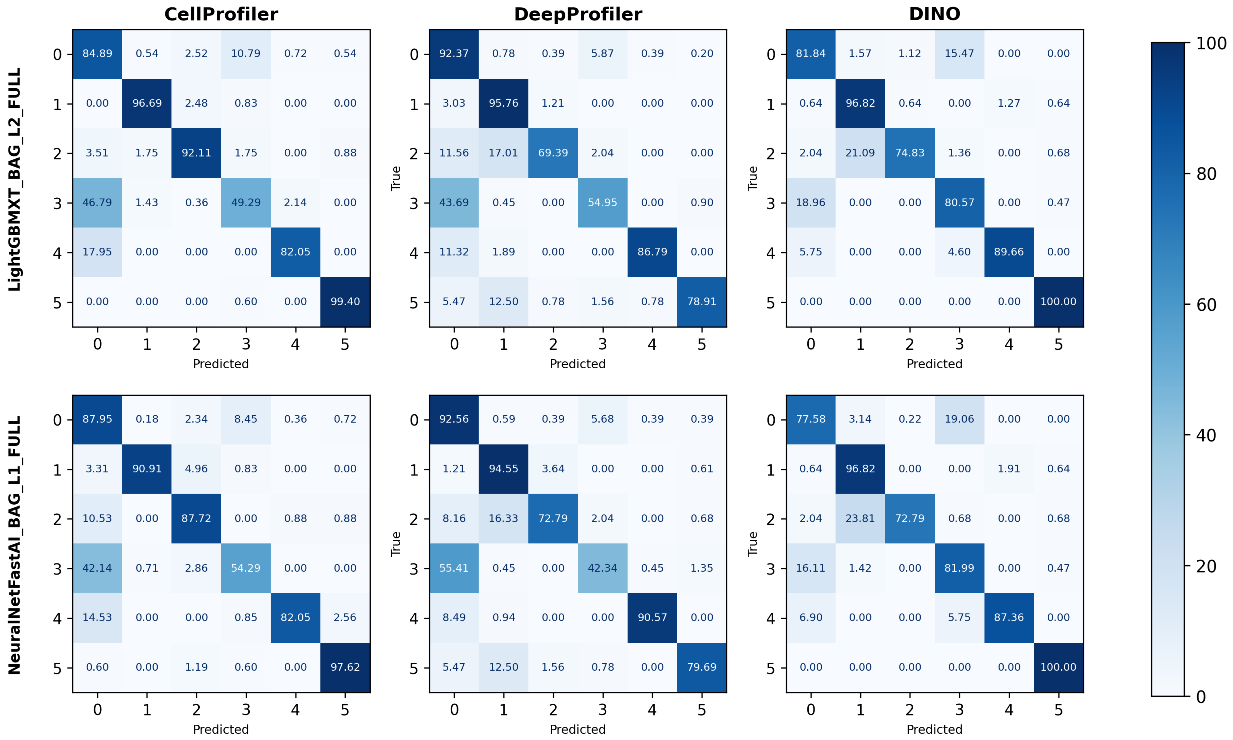


**Figure S18:** Confusion matrices of majority voted classifiers in Table 2, with label codes: 0 (apoptosis), 1 (Autophagy), 2 (Ferroptosis), 3 (Immunogenic cell death), 4 (Necroptosis), 5 (Pyroptosis). Numbers indicate percentage of correctly predicted labels in respective field. Top row: Confusion matrices for LightGBMXT_BAG_L2 models for CellProfiler, DeepProfiler and DINO (from left to right). Bottom row: Confusion matrices for NeuralNetFastAI_BAG_L1 models for CellProfiler, DeepProfiler and DINO (from left to right).

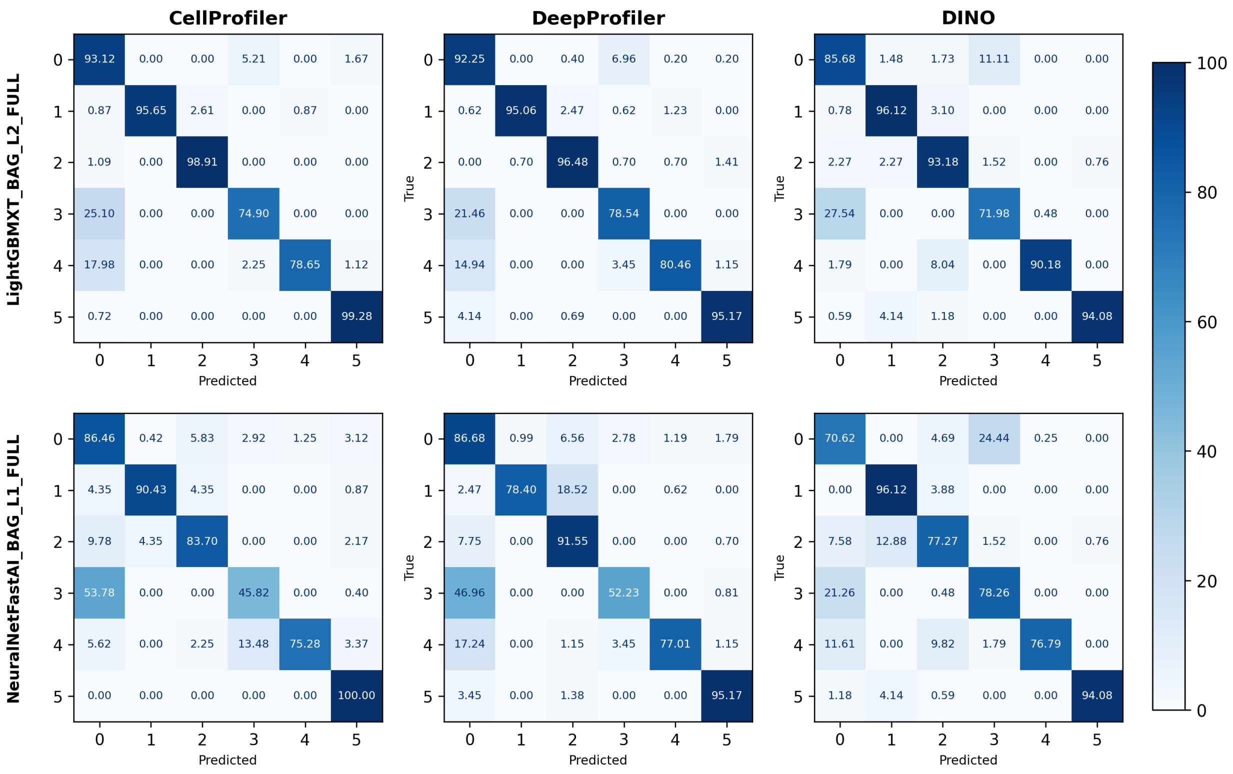


**Figure S19:** Confusion matrices of aggregated classifiers in Table 2, with label codes: 0 (apoptosis), 1 (Autophagy inducer), 2 (Ferroptosis inducer), 3 (Immunogenic cell death), 4 (Necroptosis inducer), 5 (Pyroptosis inducer). Numbers indicate percentage of correctly predicted labels in respective field. Top row: Confusion matrices for LightGBMXT_BAG_L2 models for CellProfiler, DeepProfiler and DINO (from left to right) Bottom row: Confusion matrices for NeuralNetFastAI_BAG_L1 models for CellProfiler, DeepProfiler and DINO (from left to right).

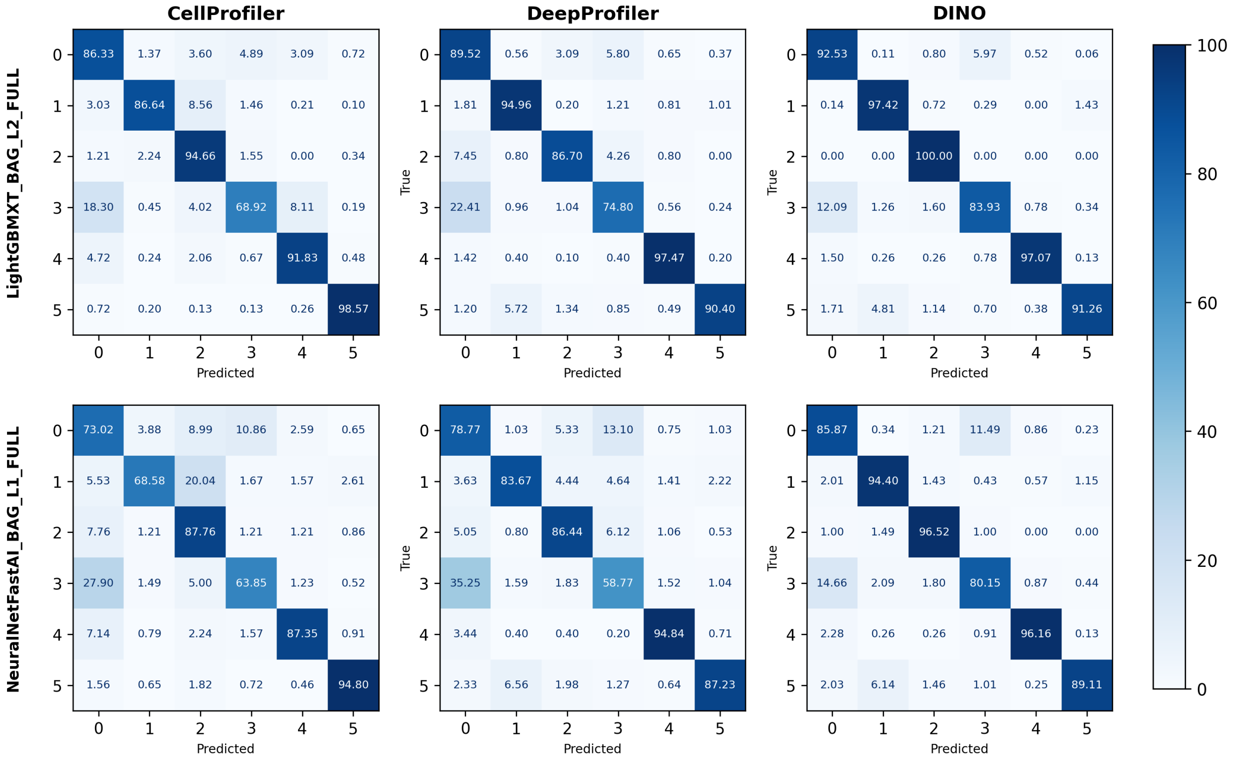


**Figure S20:** Confusion matrices of single-cell classifier on intersection of test sets, evaluated on single-cell classification models. Label codes: 0 (apoptosis), 1 (Autophagy inducer), 2 (Ferroptosis inducer), 3 (Immunogenic cell death), 4 (Necroptosis inducer), 5 (Pyroptosis inducer). Numbers indicate percentage of correctly predicted labels in respective field. Top row: Confusion matrices for LightGBMXT_BAG_L2 models for CellProfiler, DeepProfiler and DINO (from left to right) Bottom row: Confusion matrices for NeuralNetFastAI_BAG_L1 models for CellProfiler, DeepProfiler and DINO (from left to right).

**Table S1:** Table of compounds, their annotated cell death subtype, targets, and literature.

| Compound Name | Cell death subtype | Target | References |
|---|---|---|---|
| 2,3-DCPE Hydrochloride | Apoptosis | Bcl-XL | (Bai et al., 2020; S. Wang et al., 2004; S. Wu et al., 2004) |
| Anatoxin A Fumarate | Apoptosis | nAChRs | (Bownik et al., 2012; Lakshmana Rao et al., 2002) |
| Apoptosis Activator 2 | Apoptosis | APAF-1 | (J. T. Nguyen et al., 2003; T. V. Nguyen et al., 2010) |
| AT 101 | Apoptosis | Bcl-2 | (Balakrishnan et al., 2009; Kline et al., 2008) |
| AZD 2461 | Immunogenic Cell Death | PARP | (Oplustil O'Connor et al., 2016) |
| Betulinic Acid | Apoptosis | Mitochondria | (Gaul et al., 2008; Leoni et al., 2008; Schwänen et al., 2002) |
| Bleomycin | Immunogenic Cell Death | DNA | (Bugaut et al., 2013) |
| Bortezomib | Immunogenic Cell Death | 26S Proteasome | (Solimando et al., 2024) |
| Bz 423 | Apoptosis | ATP Synthase | (Blatt et al., 2008, 2009) |
| C 75 | Apoptosis | FASN | (Ho et al., 2007; Moore, 1993) |
| Camptothecin | Apoptosis | Topoisomerase I | (Glynn et al., 1992; Johnson et al., 1997; Traganos et al., 1996; Zeng et al., 2012; Zhang et al., 2000) |
| Carboplatin | Immunogenic Cell Death | DNA | (Rébé et al., 2019) |
| Cisplatin | Immunogenic Cell Death | DNA | (Martins et al., 2011) |
| Cladribine | Apoptosis | DNA Polymerase | (Johnston, 2011; Marzo et al., 2001) |
| Crizotinib | Immunogenic Cell Death | ALK, ROS1, MET | (Petrazzuolo et al., 2021) |
| Dactinomycin (Actinomycin D) | Immunogenic Cell Death | DNA | (Humeau et al., 2020) |
| Daunorubicin | Immunogenic Cell Death | Topoisomerase II | (Ocadlikova et al., 2020) |
| Domperidone | Autophagy | Dopamine Receptors | (Olubodun-Obadun et al., 2021) |
| Doxorubicin | Apoptosis | Topoisomerase II | (Müller et al., 1997; S. Wang et al., 2004) |
| Epirubicin | Apoptosis | Topoisomerase II | (T.-C. Huang et al., 2018) |
| Erastin | Ferroptosis | VDAC Modulator | (Dixon et al., 2012; Xie et al., 2016) |
| Etoposide | Apoptosis | Topoisomerase II | (Barry et al., 1993; Khedri et al., 2019) |
| Everolimus | Autophagy | mTOR | (Baraz et al., 2014; Crazzolara et al., 2009; Roccaro et al., 2012) |
| FK 866 | Apoptosis | NAMPT | (Gehrke et al., 2014; Hasmann et al., 2003) |
| FIN56 | Ferroptosis | GPX4 | (Cotto-Rios et al., 2016; Shimada et al., 2016; Xiang Wang et al., 2024) |
| Fludarabine | Apoptosis | DNA Polymerase | (Romano et al., 2000; Zinzani, Buzzi, Farabegoli, Martinelli, et al., 1994; Zinzani, Buzzi, Farabegoli, Tosi, et al., 1994) |
| G5 | Apoptosis | Ubiquitin Isopeptidase | (Aleo et al., 2006; Foti et al., 2009; Hafner-Bratkovič et al., 2018) |
| HS-173 | Necroptosis | PI3K | (J. H. Park et al., 2019) |
| Kaempferol | Apoptosis | Bcl-2 | (Kim et al., 2013) |
| L-690,330 | Autophagy | Inositol Monophosphatase | (Criollo et al., 2007; Fleming et al., 2011) |
| Loperamide | Autophagy | $\mu$-Opioid Receptors | (J. Wu et al., 2023; Zielke et al., 2018) |
| Lovastatin | Ferroptosis | HMG-CoA Reductase | (S.-W. Huang et al., 2020) |
| Mitomycin C | Apoptosis | DNA | (I. C. Park et al., 2000; Pirnia et al., 2002) |
| MPC-6827 | Apoptosis | TUBB | (Kasibhatla et al., 2007; Topçul et al., 2022) |
| Narciclasine | Apoptosis | eEF1A | (Dumont et al., 2007; Ingrassia et al., 2009; M. Wang et al., 2023) |
| Nigericin | Pyroptosis | Ion Transport | (He et al., 2015; Rozario et al., 2024) |
| Olaparib | Immunogenic Cell Death | PARP | (Chabanon et al., 2019; Xia et al., 2024) |
| Oncrasin 1 | Apoptosis | KRAS | (Guo et al., 2011) |
| Oxaliplatin | Immunogenic Cell Death | DNA | (Tesniere et al., 2010; Jingmiao Wang et al., 2022; Zhu et al., 2020) |
| Pimozide | Autophagy | Dopamine Receptors | (Ranjan et al., 2020) |
| Pravastatin | Ferroptosis | HMG-CoA Reductase | (Q. Zhou et al., 2024) |
| Polyphyllin VI | Pyroptosis | Mitochondrial Pathways | (Teng et al., 2020; Yu et al., 2021) |
| Quercetin | Necroptosis | STAT3 | (Amiri et al., 2023; Srivastava et al., 2016; Jin Wang et al., 2025) |
| Rifaximin | Apoptosis | RNA Polymerase | (Gautam et al., 2018) |
| Shikonin | Necroptosis | STAT3 | (T. Liu et al., 2019; Shahsavari et al., 2015; Xinyu Wang et al., 2022) |
| Simvastatin | Ferroptosis | HMG-CoA Reductase | (L. Wang et al., 2025; D. Zhou et al., 2022) |
| SMBA 1 | Apoptosis | BAX (Bcl-2) | (Fan et al., 2020; Xin et al., 2014) |
| SN-38 | Apoptosis | Topoisomerase I | (Nakatsu et al., 1997; Ueno et al., 2002) |
| Sorafenib | Ferroptosis | Tyrosine Kinases | (Lachaier et al., 2014; R. Liu et al., 2024) |
| Sulfasalazine | Ferroptosis | NF-$\kappa$B | (J. Liu et al., 2022; Xie et al., 2016) |
| Teniposide | Immunogenic Cell Death | Topoisomerase II | (Li et al., 2022; Z. Wang et al., 2019) |
| Topotecan | Apoptosis | Topoisomerase I | (Caserini et al., 1997; Traganos et al., 1996) |
| VU0359595 | Autophagy | M1 Muscarinic Receptors | (Cai et al., 2016) |

**Table S2:** Batch information and biological effects of selected reference compounds.

| Abbreviation | Name | Chemical Use | Biological Effect | Expected Phenotype |
|---|---|---|---|---|
| [dmso] | Dimethyl Sulfoxide | Dissolving compounds | Used as solvent control with no biological effect | No morphological changes; healthy cells |
| [etop] | Etoposide | Chemotherapeutic | Induces DNA damage leading to apoptosis | Large flat nucleoli |
| [fenb] | Fenbendazole | Anthelmintic | Disrupts microtubules and induces apoptosis | Giant multinucleated cells |
| [stau] | Staurosporine | Broad-spectrum kinase inhibitor | Induces apoptosis | Cell viability; cytotoxic control |

**Table S3:** Hyperparameters used in DINO training.

| Section | Parameter | Value |
|---|---|---|
| model | model_type | DINO |
| | arch | vit_small |
| | datatype | CellPainting |
| | image_mode | normalized_5_channels |
| | saveckp_freq | 10 |
| | batch_size_per_gpu | 100 |
| | num_channels | 5 |
| | patch_size | 16 |
| | epochs | 200 |
| | momentum_teacher | 0.996 |
| | center_momentum | 0.9 |
| | lr | 0.0005 |
| | local_crops_scale | '0.05 0.4' |
| | image_size | 224 |
| | num_workers | 20 |
